# RanBP2-dependent annulate lamellae drive nuclear pore assembly and nuclear expansion

**DOI:** 10.1101/2024.10.08.617205

**Authors:** Junyan Lin, Arantxa Agote-Arán, Yongrong Liao, Mehdi Cloarec, Leonid Andronov, Rafael Schoch, Paolo Ronchi, Victor Cochard, Rui Zhu, Erwan Grandgirard, Xiaotian Liu, Marianne Victoria Lemée, Charlotte Kleiss, Christelle Golzio, Marc Ruff, Guillaume Chevreux, Yannick Schwab, Bruno P. Klaholz, Izabela Sumara

**Affiliations:** Institute of Genetics and Molecular and Cellular Biology (IGBMC), Illkirch, France; Centre National de la Recherche Scientifique (CNRS), UMR 7104, Strasbourg, France; Institut National de la Santé et de la Recherche Médicale (INSERM), U964, Strasbourg, France; Université de Strasbourg, Strasbourg, France; Centre for Integrative Biology (CBI), Illkirch, France; European Molecular Biology Laboratory, Electron Microscopy Core Facility, Heidelberg, Germany; Université Paris Cité, CNRS, Institut Jacques Monod, F-75013 Paris, France

## Abstract

Nuclear pore complexes (NPCs) enable nucleocytoplasmic transport. While NPCs primarily localize to the nuclear envelope (NE), they also appear in cytoplasmic endoplasmic reticulum (ER) membranes called annulate lamellae (AL). Though discovered in the mid-20th century, AL’s function and biogenesis remain unclear. Previously considered exclusive to embryonic and malignant cells, we find AL in somatic mammalian cells. Under normal conditions, AL store pre-assembled NPCs (AL-NPCs) that integrate into the NE during G1 to support nuclear expansion. Upon pathological stimuli, AL transfer to the NE is impaired, leading to their cytoplasmic accumulation. RanBP2 (Nup358) is essential for AL biogenesis, with its phenylalanine-glycine (FG) repeats promoting AL-NPC scaffold oligomerization. ER-associated Climp63 (CKAP4) directs AL-NPCs to ER sheets and the NE. This AL-driven nuclear pore formation is complementary to the canonical routes, constituting a distinct NPC assembly pathway. Our work uncovers the biogenesis mechanism of AL and the nuclear function of this key cellular organelle.

## Main

Annulate lamellae (AL) are specialized subdomains within the endoplasmic reticulum (ER), characterized by a variable array of parallel-arranged layers. These assemblies feature a regular distribution of pore structures embedded in the membranes, which closely resemble nuclear pore complexes (AL-NPCs)^1,2^. AL were commonly observed in rapidly developing, differentiating, germ and malignant cells^3–11^. Despite their early discovery by McCullough in 1952^12^ and designation by Swift in 1956^13^, and subsequent reports describing AL in specific cell types^3,7,14–21^, the universal and evolutionarily conserved function as well as direct biogenesis mechanisms of AL have remained enigmatic.

It could be hypothesized that AL might be utilized by rapidly growing cells to increase the pool of nuclear envelope (NE) NPCs (NE-NPCs) and sustain optimal levels of nucleocytoplasmic transport. This process is vital for normal cellular function, and abnormal expression or localization of nucleoporins (Nups), the building units of NPCs, have been observed in cancer and neurodegenerative disorders^22–25^. However, it is currently unknown if the Nups localization defects, often visualized by low-resolution microscopy techniques as foci or granules, are linked to the presence of AL and if AL regulate nuclear function under physiological conditions.

Here, we address this long-standing knowledge gap and uncover both the universal cellular function of AL and the direct mechanisms underlying their biogenesis. We demonstrate that AL are more abundant than previously recognized and are present in the cytoplasm of various somatic cells under normal physiological conditions. In normally proliferating cells, AL contain pre-assembled AL-NPCs that contribute to nuclear pore assembly and promote nuclear expansion during G1 phase, revealing a distinct NPC assembly pathway complementary to the two previously characterized routes^26^. Under pathological conditions or stress stimuli, this supply is disrupted, leading to the clustering and expansion of AL-NPCs in the cytoplasm. Importantly, the component of the NPC cytoplasmic filaments RanBP2 (also known as Nup358) is required for the formation of AL in normal cells and for their clustering under stress conditions. N-terminal unstructured FG-repeat region of RanBP2 promotes the oligomerization of NPC scaffold components, leading to AL-NPC formation in the cytoplasm. In addition, we identify the ER-resident protein Climp63 (CKAP4) that turns out to ensure the proper localization of AL-NPCs to ER sheets and their integration into the NE. Disrupting these biogenesis mechanisms inhibits AL-NE merging, nuclear pore assembly and nuclear expansion during G1 phase, underscoring the essential role of AL in normal cycling cells.

## Results

### Single molecule localization microscopy identifies AL in a variety of somatic mammalian cells

To determine whether cytoplasmic Nup foci observed in various stressed cells represent AL, we used cellular models that have linked microtubules (MTs), fragile X-related proteins (FXRPs) (FXR1, FXR2 and FMRP), and ubiquitin-associated protein 2-like (UBAP2L) to aberrant Nup accumulation in the cytoplasm. Indeed, previous findings demonstrated an important role of dynein-mediated MT-based transport as well as chaperoning proteins FXRPs and UBAP2L in the localized assembly of Nups at the NE during interphase^25,27,28^. Nup foci were also seen in the context of a human disease such as fragile-X syndrome (FXS)^25^, which is characterized by the absence of FMRP and several other human disorders^14^. Although FXRPs-UBAP2L models do not capture all perturbations that generate cytoplasmic Nup foci, they provide a well-established framework for assessing whether these structures correspond to AL.

As expected, acute (90 minutes) MT depolymerization or deletion of UBAP2L in mCherry-Nup133 knock-in (KI) HeLa cells ((Extended Data Fig. 1a-d) led to the formation of cytoplasmic mCherry-Nup133 foci that also contained Nup62 ((Extended Data Fig. 1e). Correlative light and electron microscopy (CLEM) analysis showed these foci to be localized to parallel-aligned ER membrane structures containing stacked nuclear pore-like complexes, demonstrating that they are AL (Fig. 1a).

**Fig. 1.**
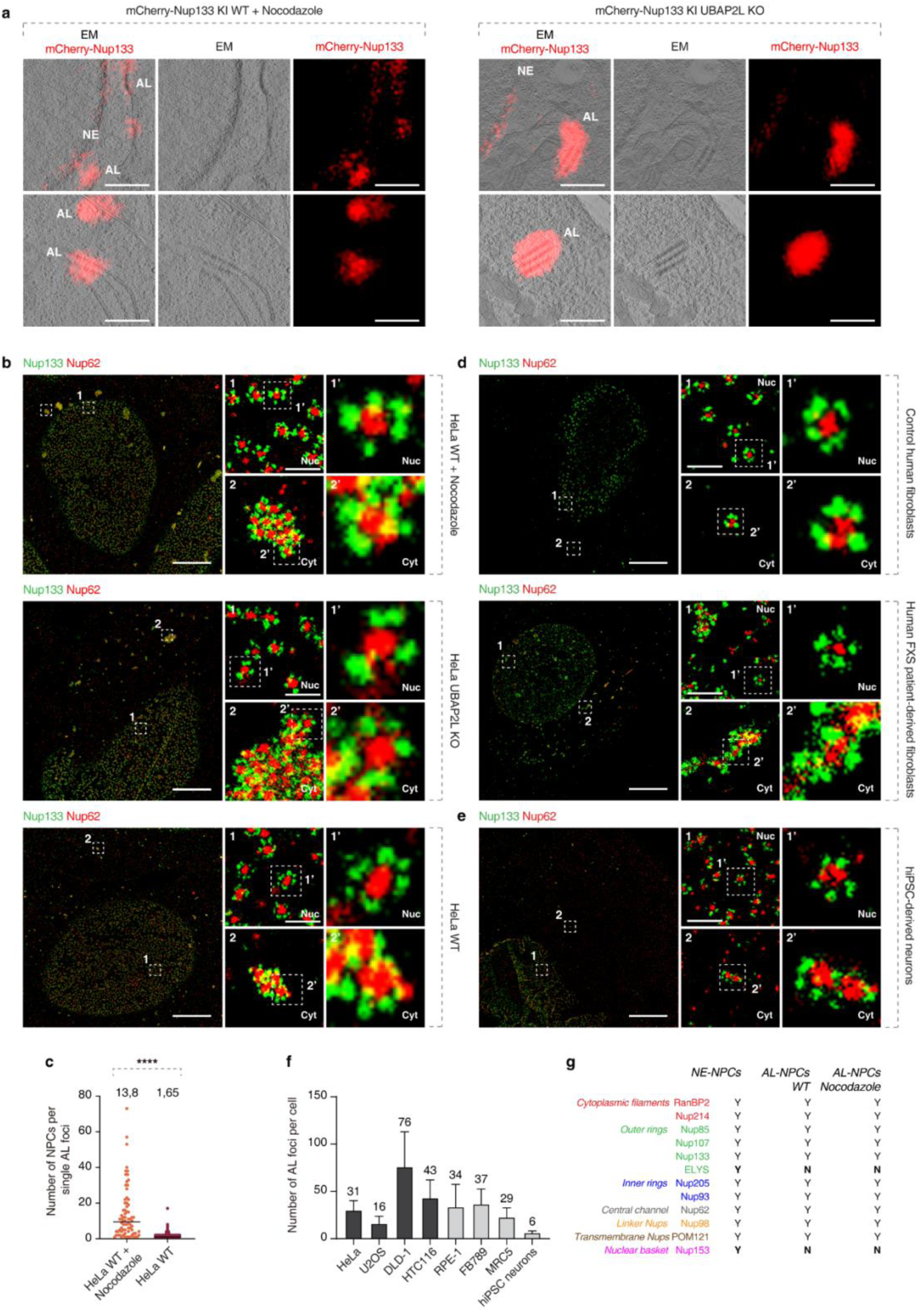
**Single molecule localization microscopy identifies AL in somatic cells.** a, Representative correlative light and electron microscopy (CLEM) of HeLa cells. The left and right graphs represent the conditions of nocodazole treatment (10 μM, 90 min) and UBAP2L knockout (KO), respectively. mCherry fluorescence is concentrated at densely packed NPCs on stacked ER sheets in the cytoplasm corresponding to AL, and is also observed at NE. Scale bars, 1 μm. b, Representative splitSMLM images depicting NPCs on the nuclear (Nuc) surface and AL-NPCs located in the cytoplasm (Cyt) in HeLa cells treated with nocodazole (10 μM, 90 min), UBAP2L KO HeLa cells and in WT HeLa cells. Nup133 signal labels the cytoplasmic and nuclear rings of the NPC, the localization of the central channel is visualized by Nup62. Note that clusters of AL-NPCs in WT cells are smaller relative to other conditions. The magnified framed regions are shown in the corresponding numbered panels. Scale bars, 3 μm (entire nuclei), 0.3 μm (zoomed regions). c, The number of single AL-NPC complexes in individual AL foci/cluster were quantified. At least 6 cells per condition were analyzed (two sample unpaired T-test, mean ± SD, *P < 0.05; ****P < 0.0001;) d, Representative splitSMLM images depicting NPCs on the nuclear (Nuc) surface and AL-NPCs located in the cytoplasm (Cyt) in normal human fibroblasts and in FXS patient-derived fibroblasts. The magnified framed regions are shown in the corresponding numbered panels. Scale bars, 3 μm (entire nuclei), 0.3 μm (zoomed regions). e, Representative splitSMLM images depicting NPCs on the nuclear (Nuc) surface and AL-NPCs located in the cytoplasm (Cyt) in hiPSC-derived neurons. The magnified framed regions are shown in the corresponding numbered panels. Scale bars, 3 μm (entire nuclei), 0.3 μm (zoomed regions). f, The number of AL-foci in individual cells in representative cancer cell lines (dark gray) and non-transformed cell lines (light grey) were quantified. At least 25 cells per cell line were analyzed. g, Summary of composition of NE-NPCs and AL-NPCs. Different colors represent specific NPC subcomplexes. “Y” indicates presence and “N” indicates absence of indicated Nups.

To overcome existing limitations in the microscopy analysis of AL, we utilized single molecule localization microscopy^29,30^ (SMLM) to generate super-resolution images of the AL foci. In particular, we applied a multi-color variant based on a dichroic image splitter and spectral demixing (splitSMLM)^31^. Via the simultaneous imaging of multiple species of fluorophores, the method enables high-precision observations of the structural organization of NPCs and other multi-component complexes, reaching a resolution of 20 nm^31^. In contrast to CLEM, this approach is easily applicable under various experimental conditions and can provide more details on the distribution and composition of AL. We confirmed the presence of highly organized cytosolic assemblies containing tightly packed NPC structures, likely corresponding to AL, upon disruption of the FXRPs-UBAP2L pathway (Fig. 1b, Extended Data Fig. 1f-h).

Surprisingly, small AL containing fewer NPCs (on average 1.65) could be observed in the cytoplasm of wild-type (WT) HeLa cells (Fig. 1b-c, Extended Data Fig. 1i), whereas in cells treated with the MT depolymerizing agent nocodazole AL were larger (average of 13.8 NPCs) (Fig. 1c). Accordingly, when imaged with conventional (diffraction-limited) fluorescence microscopy, AL-foci also appeared smaller for WT HeLa interphase cells (average size 0.090 µm^2^) compared to cells with depolymerized MTs (average size 0.213 µm^2^) (Extended Data Fig. 2a-b). Similar small cytoplasmic Nup foci were previously postulated to represent AL in normal interphasic cells^32^. We observed a relatively broad distribution range of AL foci sizes (Extended Data Fig. 2a-b) in both WT and nocodazole-treated cells, confirming the reported diversity of AL morphologies and abundance^33^. Hereafter, AL foci that were considered “large” have a size bigger than 0.3 µm^2^.

Importantly, small AL could be detected not only in a variety of common cancer cell lines including U2OS, DLD-1, HCT116, but also in the non-transformed cell lines such as RPE-1, FB789 and MRC5 (Extended Data Fig. 3a, 4a). Likewise, small AL structures were observed in normally proliferating human fibroblasts, while large AL were found in FXS patient-derived fibroblasts (Fig. 1d). Finally, AL structures were identified in human iPSCs-derived neurons (Fig. 1e, Extended Data Fig. 3a, 4a), corroborating our findings on the widespread existence of AL in somatic cells. Although AL abundance varied across different cell types (Fig. 1f), AL size consistently increased following nocodazole treatment (Extended Data Fig. 3b, 4b). These results illustrate the strength of splitSMLM as a powerful method to study AL and reveal a widespread existence of small AL in somatic, normally proliferating mammalian cells.

### AL cluster and accumulate in the cytoplasm when exposed to pathological stimuli

Small and large AL-NPCs were composed of most of the tested Nups that form different subcomplexes, with the exception of the nuclear basket Nup153 and ELYS, under various analyzed conditions (Fig. 1g, Extended Data Fig. 2c-f). Interestingly, RanBP2 was localized symmetrically on both sides of AL-NPCs (Extended Data Fig. 1f), consistent with previous reports^28,34^. In Drosophila embryos, AL-NPCs contain ELYS, but not Nup153 and Nup62^7^ and in Xenopus egg extracts, AL-NPCs did not contain ELYS, while Nup153 and Nup62 were identified^35,36^. These results suggest that the AL consist of pre-assembled NPCs, displaying composition variations across different species. Nucleocytoplasmic transport factors Exportin-1, Importin β, and Ras-related nuclear protein (Ran) likewise localized to AL foci (Extended Data Fig. 5a), as previously reported^16^, suggesting that they may be involved in AL-mediated NPC assembly, analogous to the established roles of these factors in NPC assembly pathways^10,36^. NE-resident proteins SUN1 and SUN2, but not Lamin A, Lamin B1, Nesprin, Emerin, and Lap2b, co-localized with AL foci (Extended Data Fig. 5b).

Since AL exist in various sizes and acute MT disruption lead to large AL, we studied their possible dynamic nature. Spinning disk confocal live video microscopy of WT mEGFP-Nup107 HeLa cells, synchronized in interphase, revealed a progressive increase in size and cytoplasmic clustering of AL foci over time following MT depolymerization (Extended Data Fig. 5c and Supplementary Video 1). Similar clustering events were observed upon depletion of FXR1 or in UBAP2L knockout (KO) cells (Extended Data Fig. 5d, e). These results demonstrate that small AL present in normal cells form large AL and accumulate in the cytoplasm when exposed to specific stimuli or in pathological states such as FXS (Fig. 1d).

### AL are highly dynamic and can fuse with the NE during G1 under normal conditions

Existence of small AL in normal cells prompted us to analyze their physiological role in normally proliferating cells and to address the question why pathological stimuli, such as MT disruption, lead to cytoplasmic accumulation of AL. While the conserved role of AL remains unclear, early studies in Drosophila embryos demonstrated that AL integrate into the NE to supply additional membranes and NPCs to support rapid nuclear growth, representing the first indication of the functional role of AL in the context of development^7^. To determine whether the more widespread, herein described small AL serve a similar role in non-stressed somatic cells, we first examined the abundance of AL structures during different cell cycle stages. Analysis of phosphorylated retinoblastoma (p-Rb) (G1/S phase transition marker) and cyclin B (G2 marker) revealed that small AL foci are present throughout interphase, but predominantly in the G1 and S phases (Fig. 2a, b, Extended Data Fig. 6a-d). Large AL foci induced by acute MT disruption followed the same cell cycle distribution tendency (Extended Data Fig. 6e-j). It is therefore possible that nuclear growth during the G1 phase of somatic cells is supported by AL, which primarily exist during G1 (Fig. 2a, b, Extended Data Fig. 6a-d) and which could contribute to the incorporation of new NPCs into the NE.

**Fig. 2.**
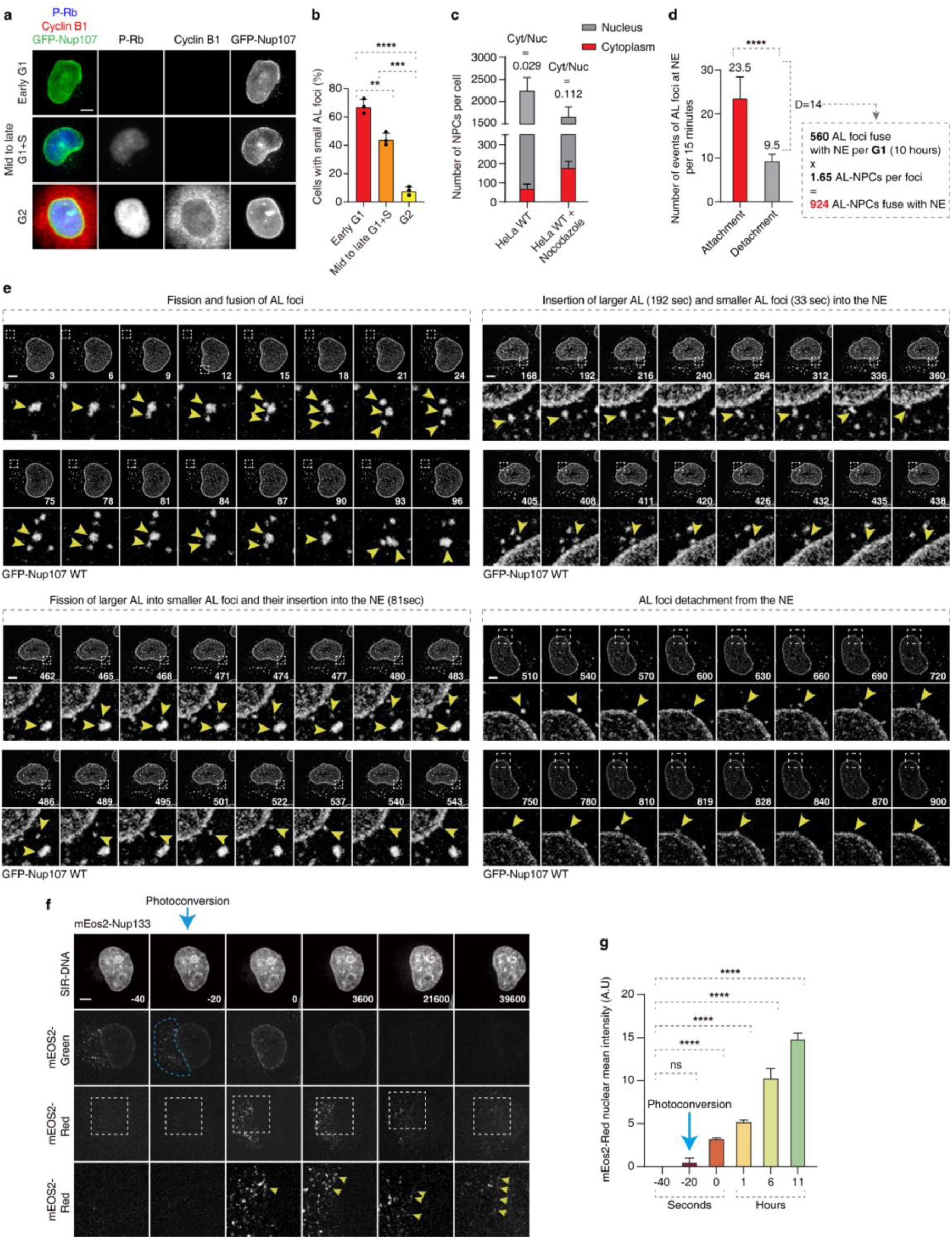
**AL are inserted into NE during G1.** a, b, Representative images of asynchronously proliferating 2xZFN-mEGFP-Nup107 HeLa cells co-labelled with anti-p-Rb (blue) and anti-cyclin B1 (red) antibodies (a). The percentage of cells with Nup foci in p-Rb and cyclin B1 negative cells (early G1), p-Rb positive and cyclin B1 negative cells (mid to late G1+S), and p-Rb and cyclin B1 positive cells (G2) was quantified in (b), and at least 200 cells per condition were analyzed (one-Way ANOVA test, mean ± SD, **P < 0.01; ***P < 0.001; ****P < 0.0001; N = 3). Scale bars, 5 μm. c, The number of NPCs in cytoplasm and nucleus of individual cells were quantified. At least 6 cells per condition were analyzed (mean ± SD) d, e, Dynamics of AL-foci in 2xZFN-mEGFP-Nup107 HeLa cells in G1 phase were analyzed by live video spinning disk confocal microscopy. Selected representative frames of the videos are depicted, and time is shown in seconds. The magnified framed regions are shown in the lower panels. Yellow arrowheads point to the movement of cytosolic AL-NPCs. Scale bars, 5 μm. The attachment and detachment events of AL foci to the NE were quantified in (d). At least 6 cells per condition were analyzed (two sample unpaired T-test, mean ± SD, ****P < 0.0001) f, g, The photoconversion of Nup foci in mEOS2-Nup133 HeLa cells were analyzed by live video spinning disk confocal microscopy. The selected representative frames of the Videos are depicted, and time is shown in seconds. The magnified framed regions of mEOS2-Red are shown in the lower panels. The blue dashed box indicates the photoconversion region, where mEOS2-Nups convert from a green to a red fluorescent state. Yellow arrowheads point to the accumulation of AL-Nup 133 at the NE. Scale bars, 5 μm. The average intensity of mEos2-Red in the nucleus at the indicated time were quantified in (g). At least 5 cells per condition were analyzed (one-Way ANOVA test, mean ± SD, ns: not significant, ****P < 0.0001)

A quantitative analysis of fixed cells estimated that AL-NPCs within the cytoplasmic AL account for approximately 3% of total nuclear NPCs, increasing to 10% following microtubule depolymerization (Fig. 2c), potentially challenging this hypothesis. However, a simple static comparison of the NPC numbers in two compartments cannot accurately determine the contribution of AL-NPC to the NE. A more time-resolved analysis of the putative dynamic nature of AL structures is required.

For this reason, we used super-resolution live-cell imaging with short time intervals between the acquisition points. We found that AL foci are highly dynamic under physiological conditions, undergoing frequent fusion and fission events (Fig. 2e, Supplementary Video 2, 3), unlike the continuous fusion observed upon microtubule depolymerization (Extended Data Fig. 5c and Supplementary Video 1). Notably, small AL foci fused more rapidly with the NE than large AL (33 and 192 seconds, respectively), and a subset of small AL originated through the fragmentation of large AL and subsequently fused with the NE (81 seconds) (Fig. 2d, e, and Supplementary Video 4). After contacting the NE, AL occasionally undergo multiple cycles of detachment and reattachment before ultimately fusing completely with the NE (Fig. 2d, e, and Supplementary Video 5).

In unstressed cells, each small AL-foci contains an average of 1.65 AL-NPCs (Fig. 1c). Given the observed net AL-NPC fusion frequency with the NE (14 events per 15 minutes) (Fig. 2d), we estimate that approximately 924 AL-NPCs are incorporated into the NE during single G1 phase of the cell cycle (10 hours). Considering that the total number of nuclear pores in HeLa cells accounts for 2000 to 4000^37^, the contribution of AL may in fact be substantial, accounting for estimated 22-45% of total nuclear NPCs.

To provide additional evidence for the AL-dependent nuclear delivery of NPCs, we generated photoconvertible mEos2-Nup133 knock-in HeLa cells (Extended Data Fig. 6k-m). The mEos2 is a green-to-red photoconvertible fluorescent protein, although its green fluorescent state lacks sufficient photostability for long-term imaging^38^, conversion of mEos2-Nup133 at the NE from green to red fluorescence at a defined interphase time point revealed no detectable diffusion of red signal from the NE into the cytoplasm (Extended Data Fig. 6n, and Supplementary Video 6). Interestingly, photoconversion of cytoplasmic AL-Nup mEos2-Nup133 foci resulted in progressive accumulation of a weak but detectable red signal at the NE (Fig.2f, g, and Supplementary Video 7), indicating that AL-NPCs can be incorporated into the nucleus. The weak nuclear signal can be likely due to progressive photobleaching of the fluorescence signal in the course of the experiment and dynamic remodeling of AL foci, which flattened and dispersed as a rim around the nuclei after migration towards NE (Fig. 2f). Future development of photoconvertible tags with improved long-term stability and brightness are required to investigate dynamics of NE incorporation events of AL in live cells in more detail. Nevertheless, taken together our data show that AL structures observed in normally proliferating cells can attach to the NE during G1.

### AL-NPC insertion into the NE through NE-ER contacts supports nuclear expansion and pore formation

To provide morphological evidence for AL incorporation into the NE, we next used splitSMLM to examine the structural organization and composition of AL-NPCs. This analysis visualized symmetric AL-NPCs in the proximity of or directly attached to the NE containing asymmetric NE-NPCs in WT interphase cells (Fig. 3a panels 1-2). AL-NPCs fusion events into the NE were observed where symmetric AL-NPCs were positioned laterally onto the NE (Fig. 3a panels 2-4). Some fusion events were accompanied by the formation of lateral NE openings containing both AL-NPCs and NE-NPCs next to each other (Fig. 3a panels 5-6).

**Fig. 3.**
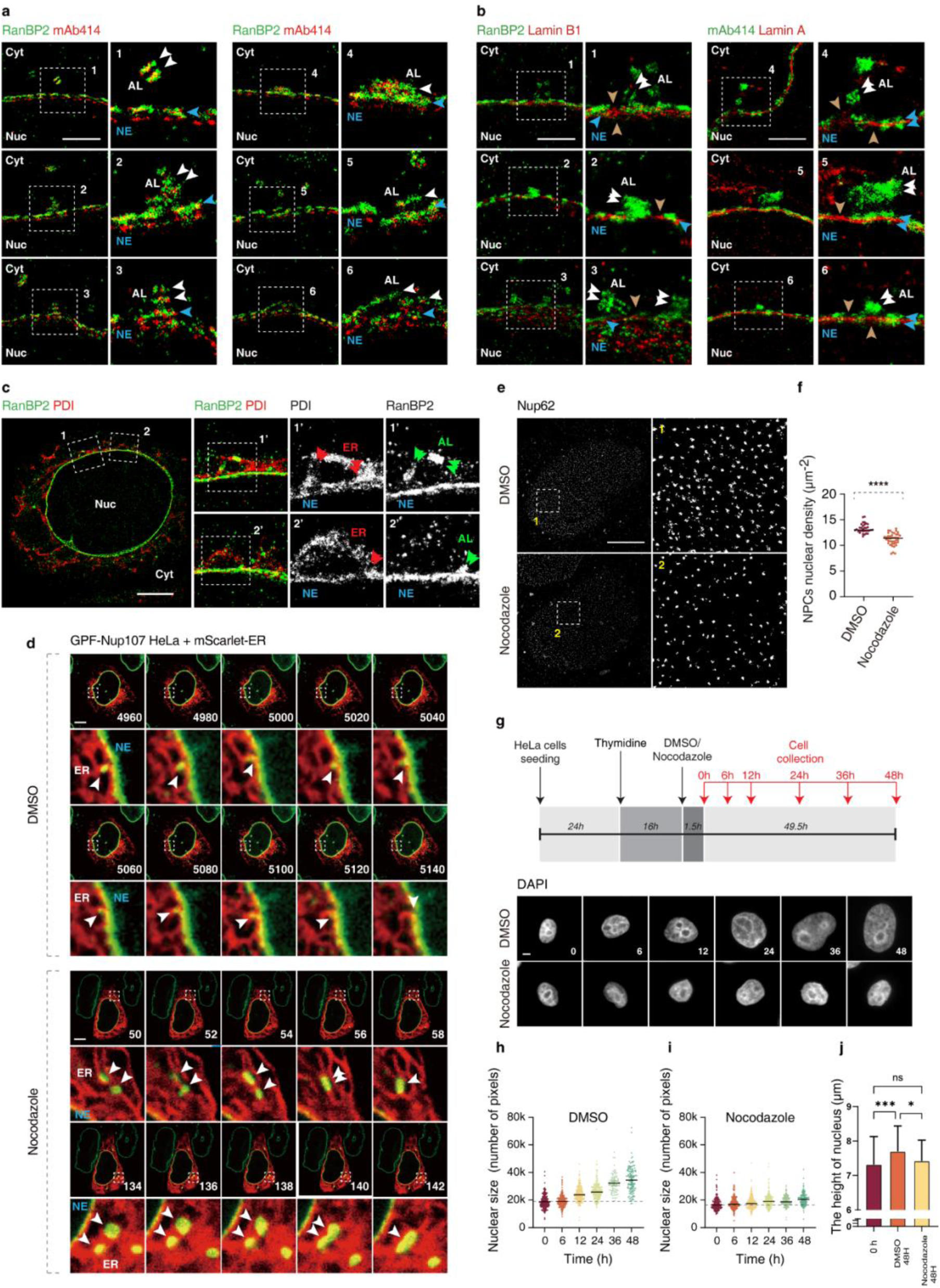
**AL-NPC insertion into the NE through NE-ER contacts supports nuclear expansion and pore formation.** a, Representative splitSMLM images depicting the nuclear (Nuc) surface and cytoplasm (Cyt) from the side view of WT HeLa cells. Central channel, cytoplasmic filaments and Nucleus basket of NPCs is labelled with mAb414 antibodies and RanBP2 antibody labels NPC cytoplasmic filaments. The magnified framed regions are shown in the corresponding numbered panels. White arrowheads point out to AL-NPC structures within AL where NPC cytoplasmic filaments are located symmetrically and blue arrowheads indicate asymmetric NE-NPCs on the nuclear envelope (NE) (panel 1). Note lateral fusion events of AL-NPCs with NE-NPCs from the cytoplasmic side (panels 2-4) and NE-openings where both AL-NPCs and NE-NPCs can be located (panels 5-6). Scale bars, 1 μm. b, Representative splitSMLM images depicting the nuclear (Nuc) surface and cytoplasm (Cyt) from the side view of WT HeLa cells. RanBP2 antibody labels NPC cytoplasmic filaments, Lamin B1 and Lamin A label NE-associated factors. Central channel, cytoplasmic filaments and nuclear basket of NPCs are labelled with mAb414 antibodies. The magnified framed regions are shown in the corresponding numbered panels. White arrowheads point out to AL-NPC structures within AL where NPC cytoplasmic filaments are located symmetrically and blue arrowheads indicate asymmetric NE-NPCs on the nuclear envelope (NE). Brown arrows indicate NE-openings where NE-associated factors remain intact. Scale bars, 1 μm. c, Representative splitSMLM images depicting the nuclear (Nuc) surface and cytoplasm (Cyt) from the side view of WT HeLa cells. RanBP2 antibody labels NPC cytoplasmic filaments and PDI labels ER. The magnified framed regions are shown in the corresponding numbered panels. Red arrows point out to ER, and green arrows indicate AL-NPCs. Scale bars, 3 μm. d, 2xZFN-mEGFP-Nup107 HeLa cells expressing ER marker plasmid mScarlet-ER were analyzed by live video spinning disk confocal microscopy. The selected representative frames of the Videos are depicted, and time is shown in seconds. The magnified framed regions are shown in the lower panels. White arrowheads point to the movement of cytosolic AL-NPC foci along the ER and their incorporation into the NE along the junction of the ER and the NE. Scale bars, 5 μm. e, f, Representative SMLM immunofluorescence images of Nup62 at the nuclear surface in DMSO or nocodazole treated (10 μM, 90 min) HeLa cells. The magnified framed regions are shown in the right panels (e). The nuclear density of NE-NPCs (Nup62) in cells was quantified in (f) (two sample unpaired T-test, mean ± SD, ****P < 0.0001, unpaired two-tailed t test; 40 cells were counted per cell line). Scale bars, 5 μm g-j, Scheme of the experimental setup (g). HeLa cells were synchronized and arrested at S phase by thymidine treatment (16h), treated with nocodazole (10 μM) and fixed and collected at indicated time points. Representative images of cell nuclei are shown in (g) and the nuclear size was quantified in (h, i). The height of nucleus was quantified in (h, i). About the nuclear size, at least 200 cells per condition were analyzed (N = 3). About the height of nucleus, at least 80 cells per condition were analyzed (N = 3). Scale bars, 5 μm.

Analysis of nuclear lamina components Lamin A and B1 by splitSMLM revealed that despite the presence of these NE incisions, the underlying nuclear membrane scaffold remained intact (Fig. 3b). Importantly, the analysis of the ER marker PDI, confirmed that the sites of AL-NPC insertion within these NE openings precisely colocalized with the ER and ER-NE contact sites (Fig. 3c). Thus, the splitSMLM analysis confirms the observed AL structures in normal cells as specialized sub-compartments of the ER, which is known to be continuous with the outer NE and contribute to its formation^39^. These observations are also in agreement with a previously proposed model suggesting “*en bloc*” insertion of AL into the NE and remodeling of AL-NPCs by GTPase Ran to become asymmetric functional NPC structures in fly embryos^7,10,34^, demonstrating evolutionary conservation of this process. Although 20 nm resolution images from splitSMLM analysis strongly suggest insertion of intact AL into NE, at this point we cannot formally exclude the possibility that AL-NPCs undergo local disassembly or that AL act solely as donors of NPC subcomplexes prior to their NE insertion.

Importantly, spinning disk confocal live video microscopy of the ER marker mScarlet-ER confirmed that AL foci move along the ER to merge with the NE in WT interphase cells (Fig. 3d, and Supplementary Video 8). MT depolymerization is predicted to modulate ER dynamics^40^, damaging reticulated ER tubules and leading to vesiculation and thickening of cisternal sheets (Extended Data Fig. 7a), which could explain the inhibition of AL foci transfer to the NE and their clustering (Fig. 3d, Extended Data Fig. 5c and Supplementary Video 9). Interestingly, AL-NPCs were also often localized near microtubules (Extended Data Fig. 7b). Live-cell imaging further revealed a consistent spatial association between AL and the microtubule network (Supplementary Video 10), suggesting that they may utilize MT-ER interactions to merge with the NE and to increase the pool of NE-NPCs. Our results indicate that MT depolymerization in somatic cells disrupts ER dynamics, thereby impairing the transfer of AL-NPCs to the NE, resulting in their cytoplasmic accumulation. However, we cannot exclude the possibility that the transport of soluble Nups along microtubules also contributes to AL-NPC assembly. In support of this, studies in *Drosophila* oocytes have shown that microtubules mediate the delivery of distinct Nup condensates to AL-NPC assembly sites a process essential for efficient AL-NPC formation^10^.

Since acute MT depolymerization inhibited AL-NPC transfer to the NE, we set out to analyze the physiological consequences of this inhibition. The density of NE-NPCs (Fig. 3e, f) and the level of Nups at NE (Extended Data Fig. 7c, d) decreased upon acute nocodazole treatment. The short treatment excluded any possible indirect effects of nocodazole on mitotic progression. Strikingly, when cells are arrested in the G1/S phase, the nucleus continues to grow beyond its normal size, and acute MT depolymerization inhibits nuclear growth in G1/S-arrested WT cells (Fig. 3g-j). Nuclear expansion is sustained by a constant supply of many proteins and lipids from the ER to the NE^41^, a process which could also be disrupted by MT depolymerization. The insertion of AL-NPCs may support nuclear expansion by potentiating nucleocytoplasmic transport rates, or the supply of lipids and proteins required for NE expansion may occur simultaneously with AL insertion into the NE. Taken together, AL are more widespread than previously anticipated and exist in WT somatic cells. AL act to supply a relatively large pool of AL-NPCs (estimated 924) to the NE, thereby promoting the nuclear expansion specifically during G1 phase of the cell cycle. AL are inserted into the NE at the ER-NE junctions through ER dynamics, and this process requires an intact microtubule network.

These findings not only establish AL as functional determinants of nuclear architecture during normal mammalian cell proliferation but also indicate existence of an additional specialized pathway for NPC assembly during normal cell cycle progression.

### RanBP2 is required for the clustering of small AL into large AL under stress conditions but does not regulate protein levels of Nups

Having demonstrated a cellular function of AL in normally growing cells, we next aimed to uncover the mechanisms driving AL biogenesis. The component of NPC cytoplasmic filaments RanBP2 (also known as Nup358) has been shown to localize to AL structures and it has been proposed to regulate a Nup condensate fusion mechanism supporting AL biogenesis during Drosophila oogenesis^10^. However, direct evidence for RanBP2’s role in AL formation is missing. It is also unknown if the role of RanBP2 is conserved in somatic cells and in other species, and what the molecular basis for possible RanBP2 function on AL is. RanBP2 was required for the formation of large AL foci induced by MT depolymerization (Fig. 4a-c, Extended Data Fig. 7e-l) and by inactivation of FXRPs-UBAP2L-dynein pathway in several human cell lines tested (Extended Data Fig. 7m-r). Another component of NPC cytoplasmic filaments, Nup214, was not required for the formation of large AL foci (Extended Data Fig. 8a-c). Higher eukaryotes utilize two NE-NPC assembly pathways, each associated with different cell cycle stages: the postmitotic and the interphase pathway^26^. We found that downregulation of ELYS (required for postmitotic NE-NPC assembly), or Nup153 or POM121 (driving interphase NE-NPC assembly) did not inhibit AL foci formation when induced by acute nocodazole treatment Extended Data Fig. 8a-f). In contrast, defects in NE-NPC biogenesis were reported to induce AL^24^ and as expected, downregulation of ELYS or Nup153 led to the formation of large AL foci in untreated cells (Extended Data Fig. 8a, d, g). Downregulation of RanBP2 strongly inhibited the formation of large AL foci induced by Nup153 or ELYS depletion (Fig. 4d-i), suggesting its universal and widespread role in AL clustering.

**Fig. 4.**
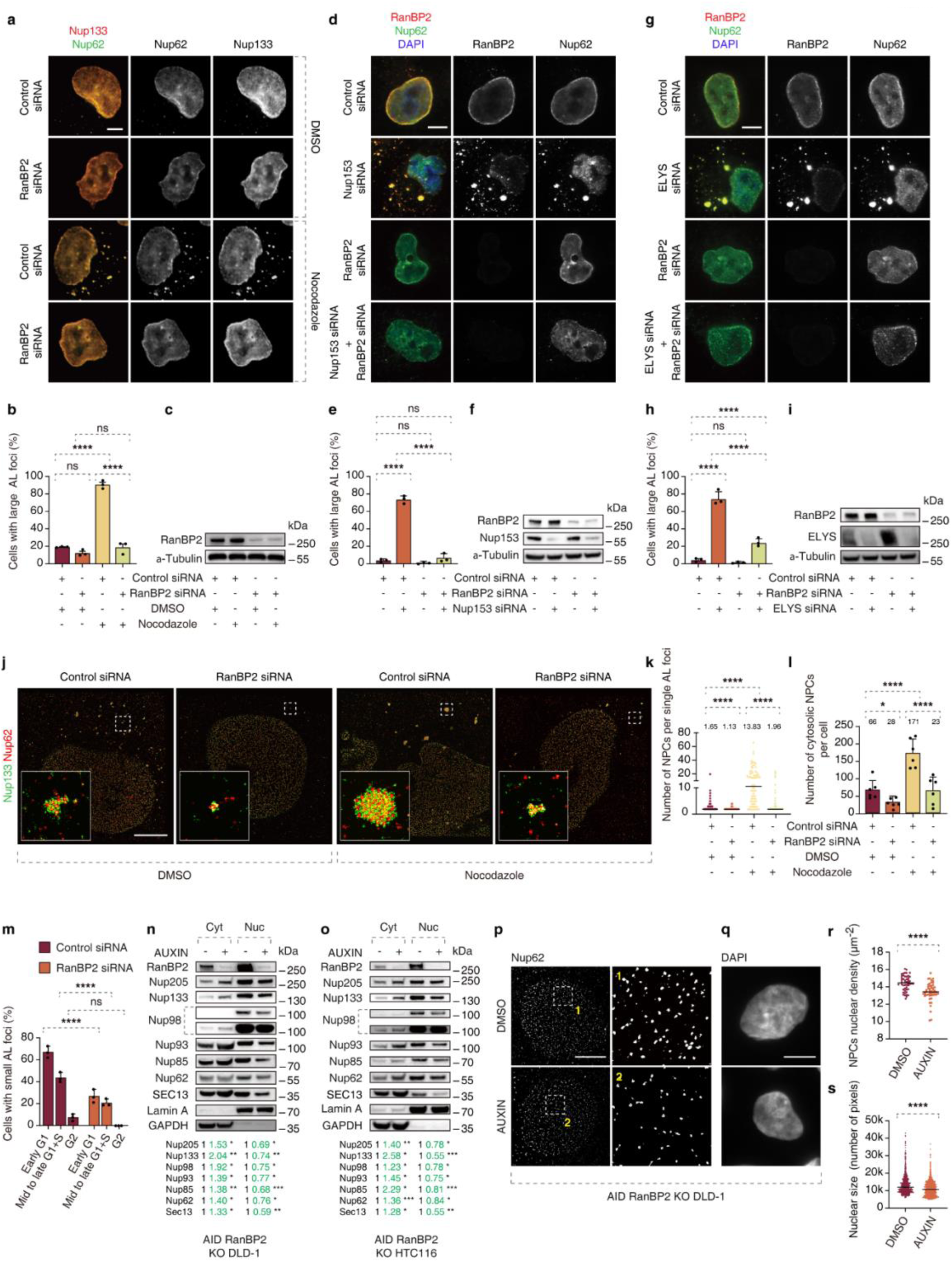
**RanBP2 is required for AL clustering, AL biogenesis and for nuclear pore formation.** a-c, Representative images of HeLa cells treated with the indicated siRNAs (48h) and subsequently by nocodazole (10 μM) or DMSO (90 min). Cells were co-labelled with anti-Nup62 (green) and anti-Nup133 (red) antibodies (a) to detect AL-NPC foci. The number of cells with big AL foci were quantified in (b), and at least 1000 cells per condition were analyzed (one-Way ANOVA test, mean ± SD, ns: not significant, ****P < 0.0001; N = 3). Scale bars, 5 μm. (c) shows the Western blot analysis. d-f, Representative images of HeLa cells treated with the indicated siRNAs (48h). Cells were co-labelled with anti-Nup62 (green) and anti-RanBP2 (red) antibodies (d) to detect AL-NPC foci. The percentage of cells with big AL foci was quantified in (e), and at least 300 cells per condition were analyzed (one-Way ANOVA test, mean ± SD, ns: not significant, **P < 0.01; ****P < 0.0001; N = 3). Scale bars, 5 μm. (f) shows the Western blot analysis. g-I, Representative images of HeLa cells treated with the indicated siRNAs (48h). Cells were co-labelled with anti-Nup62 (green) and anti-RanBP2 (red) antibodies (g) to detect AL-NPC foci. The percentage of cells with big AL foci were quantified in (h), and at least 300 cells per condition were analyzed (one-Way ANOVA test, mean ± SD, ns: not significant, **P < 0.01; ****P < 0.0001; N = 3). Scale bars, 5 μm. (i) shows the Western blot analysis. j-l, Representative splitSMLM images of HeLa cells treated with the indicated siRNAs (48h) and subsequently by nocodazole (10 μM) or DMSO (90 min) depicting AL-NPCs in the cytoplasm. Cells were co-labelled with anti-Nup62 (green) and anti-Nup133 (red) antibodies (j). The magnified framed regions are shown in the lower left corner. The number of single AL-NPC complexes per cell and the number of single AL-NPC complexes in individual AL foci/cluster were quantified in (k, l), respectively. At least 6 cells per condition were analyzed (one-Way ANOVA test, mean ± SD, *P < 0.05; ****P < 0.0001; N = 3). Scale bars, 3 μm. m, Asynchronously proliferating 2xZFN-mEGFP-Nup107 HeLa cells treated with the indicated siRNAs (48h) and co-labelled with anti-P-Rb (blue) and anti-cyclin B1 (red) antibodies. The percentage of cells with small AL foci in P-Rb and cyclin B1 negative cells (early G1), P-Rb positive and cyclin B1 negative cells (mid to late G1+S), and P-Rb and cyclin B1 positive cells (G2) was quantified in (m), and at least 200 cells per condition were analyzed (one-Way ANOVA test, mean ± SD, ****P < 0.0001; N = 3). Representative images are shown in Extended Data Fig. 11a. n, o, Auxin-inducible degron (AID) RanBP2 KO DLD-1 cells (n) and HTC116 cells(o) were treated with DMSO or auxin for 4h to deplete RanBP2. Cytoplasmic (Cyt) and nuclear (Nuc) fractions were prepared and levels of Nups were analyzed by Western blot (at least 3 independent experiments). Signal intensities were quantified; signals in the cytoplasm and nucleus were normalized according to GAPDH and Lamin A. A mean value is shown below the image (*P < 0.05, **P < 0.01, ***P < 0.001, paired one-tailed t-test; n = 3 independent experiments). p, r, Representative SMLM immunofluorescence images of Nup62 at the nuclear surface in AID RanBP2 KO DLD-1 cells treated with DMSO or auxin for 4h to deplete RanBP2. The magnified framed regions are shown in the right panels (p). The nuclear density of NPCs (Nup62) in cells was quantified in (r) (mean ± SD, ****P < 0.0001, unpaired two-tailed t test; counted 50 cells per cell line). Scale bars, 5 μm q, s, Representative images of cell nuclei in AID RanBP2 KO DLD-1 cells treated with DMSO or auxin for 4h to deplete RanBP2 are shown in (q) and the nuclear size was quantified in (s). At least 700 cells per condition were analyzed (mean ± SD, ****P < 0.0001, unpaired two-tailed t test; N = 3). Scale bars, 5 μm.

Is RanBP2 also required for the maintenance of the AL structures? Downregulation (Extended Data Fig. 9a-f) or auxin-inducible depletion (Extended Data Fig. 9g-k) of RanBP2 ^42^ after nocodazole treatment severely decreased the number of large AL foci and their clustering observed upon MT depolymerization. Thus, RanBP2 is required for the formation and the maintenance of large AL. This process is likely to be independent of reported NPC degradation mechanisms involving autophagy^43,44^, since RanBP2 downregulation inhibited large AL foci formation also in the presence of lysosomal inhibitors (Extended Data Fig. 10a-h).

Previous studies have shown that RanBP2 knockdown reduces translation of reporter constructs^45^, whereas we show that deletion or downregulation of RanBP2 did not alter the protein levels of multiple Nups across various cell lines (Extended Data Fig. 10i-k). Notably, the use of auxin-inducible RanBP2 depletion minimizes the confounding effects of chronic translational defects (Extended Data Fig. 10k). Therefore, the observed reduction in AL is unlikely to result from global translational regulation of Nups by RanBP2 under these conditions. The RanBP2 function on AL could also not be explained by possible effects on levels of FXR1 and UBAP2L, the factors driving spatial assembly of Nups during early interphase^28^, as they were not affected by auxin-inducible deletion of RanBP2 in DLD-1 (Extended Data Fig. 10l) and in HTC116 cells (Extended Data Fig. 10m). Thus, RanBP2 is required for the clustering of small AL into large AL and for AL maintenance but does not regulate protein levels of Nups or Nup-interacting factors.

### RanBP2 is required for biogenesis of small AL and their nuclear function during G1 in non-stressed cells

Is RanBP2 also required for the formation and function of small AL that we identified under normal growing conditions? The splitSMLM analysis revealed a relatively broad distribution of the number of NPC complexes in a single AL-NPC structure (Fig. 4j-l) and as expected, acute MT depolymerization increased both the number of individual AL-NPCs in single AL foci as well as the total number of AL-NPCs per cell relative to WT cells (Fig. 4j-l). Importantly,

RanBP2 was required for the formation of individual AL-NPCs, as well as their clustering both in WT and in nocodazole-treated cells (Fig. 4j-l). AL-NPCs were reported to lack nuclear basket components^7,35,36^, whereas RanBP2 appears to be localized symmetrically on both sides of AL-NPCs^28^ (Fig. 3a, b, Extended Data Fig.1f) and RanBP2 can oligomerize to form multimers^45^, providing a possible explanation for its ability to cluster individual AL-NPCs. In accordance with its role in AL-NPC biogenesis, RanBP2 was required for the formation of small AL foci in WT cells from G1 to S-phase but not in G2 stage (Fig. 4m, Extended Data Fig. 11a) and RanBP2 downregulation by siRNA decreased levels of Nups at the NE (Extended Data Fig. 11b, c). Small AL foci and the levels of Nups at the NE during interphase also decreased using auxin-inducible deletion of RanBP2 (Extended Data Fig. 11d-h), suggesting that RanBP2-mediated formation of small AL is a prerequisite for their merging with NE and increasing the pool of NE-NPCs. Short-term deletion of RanBP2 reduced levels of several Nups in isolated nuclear fractions and increased their abundance in the cytoplasm in DLD-1 (Fig. 4n) and HTC116 cells (Fig. 4o), consistent with the idea that RanBP2 does not regulate the total protein levels of Nups (Extended Data Fig. 10i-k) but their localization to the NE. Indeed, RanBP2 deletion resulted in reduced NE-NPC density (Fig. 4p, 4r).

Can RanBP2 also contribute to the nuclear pore function? We analyzed the gradient of endogenous Ran, a guanine nucleotide triphosphatase, as previously reported^28,46^, owing to the fact that Ran protein is actively imported to the nucleus with the help of transport factors where it then detaches from its cargo and is swiftly exported to begin the cycle again. Downregulation of RanBP2 (Extended Data Fig. 12a, 12b) increased the nuclear-cytoplasmic (N/C) ratio of Ran, suggesting defects in the nuclear export rate of Ran protein. Acute RanBP2 depletion likewise increased the nuclear-to-cytoplasmic (N/C) ratio of Ran and these defects were stronger in G1 cells than in G2 cells (Extended Data Fig. 12c,12e), indicating that AL-and RanBP2-dependent NPC assembly is particularly important in early interphase. Defects generated by 24-hour RanBP2 depletion were stronger in G2 cells than in G1 cells, reflecting the cumulative effect of defects (Extended Data Fig. 12d,12e). Importantly, in wild-type cells the N/C ratio of Ran was lower in G2 than in G1, suggesting an increase in the nuclear export rate of Ran as the cell cycle progresses and a hypothesis that nucleocytoplasmic transport properties change as the nucleus grows. The use of the light-inducible nuclear export system (LEXY)^47^, confirmed the export defects upon acute depletion of RanBP2 (Extended Data Fig. 12f,12g).

Our data in normally proliferating cells demonstrated a role for AL in supplying the NE-NPC pool, thereby supporting the nuclear expansion during early interphase. Since RanBP2 was required for AL-NPC formation in WT cells, we next analyzed the role of RanBP2 in nuclear function. Importantly, short-term depletion of RanBP2 was sufficient to reduce nuclear size (Fig. 4q, 4s), expanding our observations in nocodazole-treated cells (Fig. 3g-i). We conclude that RanBP2-mediated AL-NPC assembly can preserve nuclear pore density as well as nuclear growth during interphase, confirming the critical role of cytosolic NPCs in the nuclear function. Collectively, RanBP2 plays direct and widespread roles in AL, being essential for the biogenesis of single AL-NPCs, their clustering and AL maintenance in multiple stressed and normally proliferating somatic cells.

### FG repeats within the N-terminal domain of RanBP2 are required for the assembly of AL and their nuclear function

Next, we set out to study the precise molecular mechanisms of RanBP2-mediated AL biogenesis. First, to understand if the function of RanBP2 on AL is specific and which functional domains of this large protein (Fig. 5a) are required for AL formation, we performed rescue experiments using RanBP2 fragments. The full-length (FL) (aa 1-3224) and the N-terminal (NT) fragment of RanBP2 (aa 1-1171) efficiently reversed the inhibition of nocodazole-induced large AL foci observed upon downregulation of RanBP2, and the phenylalanine-glycine (FG) repeat region (832-1171) was essential for RanBP2 AL clustering function (Fig. 5a-e). The effects of auxin-inducible deletion of RanBP2 on AL formation could also be efficiently rescued by NT fragment of RanBP2 (aa 1-1171) but not by the version missing the FG repeat region (1-832) (Extended Data Fig. 13a-e).

**Fig. 5.**
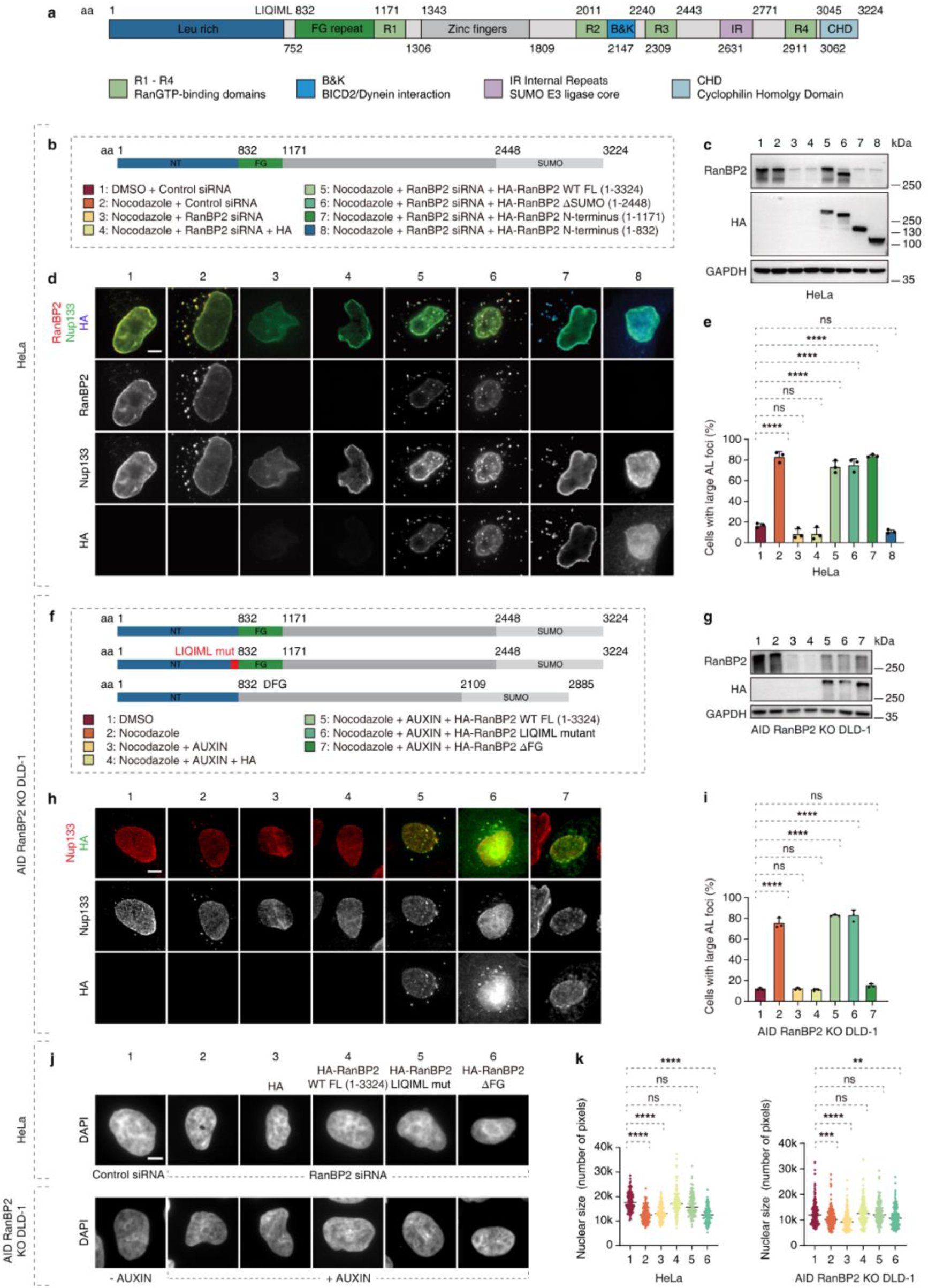
**RanBP2 is specifically required for AL clustering and biogenesis through its N-terminal FG repeat region.** a, shows the scheme of domain organization of RanBP2 protein. b-e, Rescue experiments using different RanBP2 protein fragments (b). (c) shows the Western blot analysis and (d) shows representative images of HeLa cells treated with the indicated siRNAs, transfected with different RanBP2 fragments (indicated by numbers in the schematic diagram) and treated with nocodazole (10 μM) or DMSO (90 min). Cells were co-labelled with anti-Nup133 (green), anti-RanBP2 (red) and anti-HA (blue) antibodies. The number of cells with large AL foci was quantified in (e), and at least 300 cells per condition were analyzed (one-Way ANOVA test, mean ± SD, ns: not significant, ****P < 0.0001; N = 3). Scale bars, 5 μm. f-i, Rescue experiments using different FL RanBP2 mutated versions (f). (g) Shows the Western blot analysis of RanBP2 version under different indicated conditions (numbers in the schematic diagram). (h) shows representative images of AID-RanBP2 KO DLD-1 cells treated with DMSO or Auxin, transfected with different RanBP2 versions and treated with nocodazole (10 μM) or DMSO (90 min). Cells were co-labelled with anti-Nup133 (red) and anti-HA (green) antibodies. The number of cells with large AL foci was quantified in (i), and at least 300 cells per condition were analyzed (one-Way ANOVA test, mean ± SD, ns: not significant, *P < 0.05; **P < 0.01; ****P < 0.0001; N = 3). Scale bars, 5 μm. j, k, Representative images of cell nuclei in HeLa cells and AID RanBP2 KO DLD-1 cells are shown in (j). Cells were treated with RanBP2 siRNA or Auxin to deplete RanBP2, and then rescued using different RanBP2 versions. Schematic diagram of RanBP2 versions and Western blot analysis are shown in Fig S12J-L. The nuclear size was quantified in (k). At least 120 Cell (HeLa) or 150 cells (DLD-1) per condition were analyzed (one-Way ANOVA test, mean ± SD, ns: not significant, **P < 0.01; ***P < 0.001, ****P < 0.0001; N = 3). Scale bars, 5 μm.

Importantly, full-length RanBP2 with a deletion of the FG region, which is not expected to affect other cellular functions of RanBP2, failed to restore large AL foci in RanBP2-depleted cells by either auxin-inducible degron (Fig. 5f-i) or RanBP2-specific siRNAs (Extended Data Fig. 13f-i), confirming an essential role of RanBP2 FG repeats in AL accumulation. In contrast, mutation of the LIQIML motif, which is important for RanBP2 targeting to the NE and its function on NE-NPCs assembly^45^, inhibited efficient NE localization of full-length RanBP2 leading to its nuclear and cytoplasmic localization, as expected, but successfully rescued AL defects observed under both RanBP2-deleting experimental approaches (Fig. 5f-I, Extended Data Fig. 13f-i). These results further support the conclusion that FG region of RanBP2 regulates AL formation independently of other known cellular functions of this protein and identify a mutant form of RanBP2 which can separate its functions at the NE and on AL.

These findings enabled us to further assess the role of AL in nuclear expansion in untreated WT cells. Full-length WT and RanBP2 LIQIML mutant, but not full-length RanBP2 without FG region, successfully rescued the nuclear expansion defect caused by RanBP2 depletion by auxin-inducible degron or siRNA (Extended Data Fig. 13j-l, Fig. 5j-k). Taken together, these results demonstrate the important and specific role of the FG region of RanBP2 in AL formation and AL-driven nuclear function.

### RanBP2 promotes oligomerization and assembly of AL-NPCs scaffold components through its N-terminal domain

To corroborate the role of the NT region of RanBP2, we set out to study its molecular interactions relevant for AL biology. Ectopic expression of NT RanBP2 (aa 1-1171) (Extended Data Fig. 14a-d) but not of FL Nup85 or FL Nup133 (Extended Data Fig. 14e, 14f), induced AL foci where multiple Nups, with the exception of ELYS and Nup153 (Extended Data Fig. 14c), could co-localize, relative to controls. CLEM analysis (Fig. 6a) and splitSMLM (Fig. 6b) confirmed that these foci represent AL. Ectopic expression of NT RanBP2 increased the number of individual AL-NPCs per cell and the size of AL (Fig. 6b-d). We conclude that elevated levels of NT RanBP2 fragment can induce AL under normal growth conditions.

**Fig. 6.**
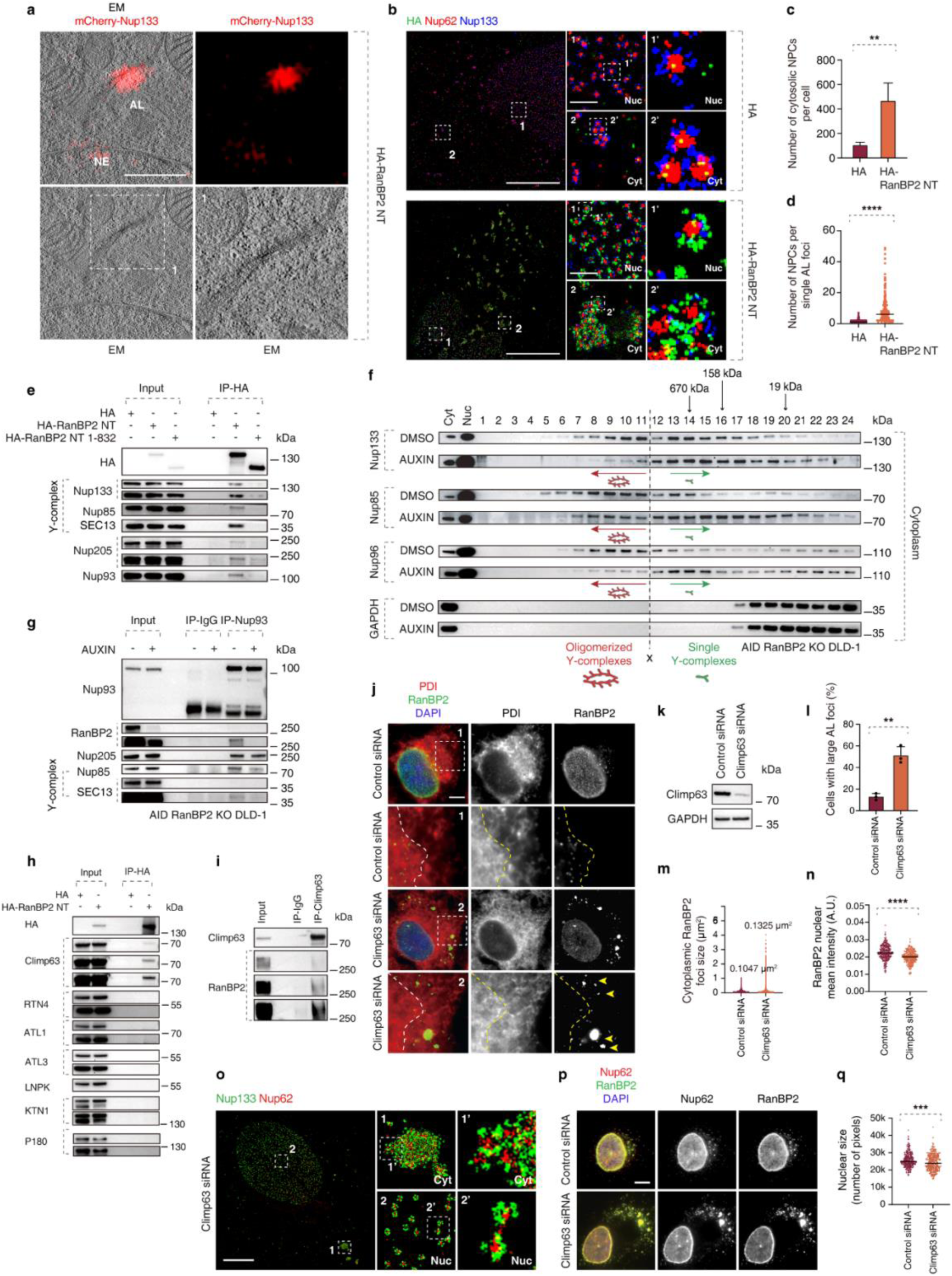
**RanBP2 drives oligomerization of AL-NPCs and interacts with Climp63 that localizes AL-NPCs to ER sheets.** a, Representative correlative light and electron microscopy (CLEM) of mCherry-Nup133-KI HeLa cells transfected with HA-RanBP2-NT fragment. The magnified framed regions are shown in the corresponding numbered panels. mCherry fluorescence is concentrated at densely packed AL-NPCs on stacked ER sheets in the cytoplasm, representing AL and is also observed at the NE. Scale bars, 1 μm. b-d, Representative splitSMLM images depicting NPCs on the nuclear surface (Nuc) and AL-NPCs in the cytoplasm (Cyt) in HeLa cells transfected with HA or HA-RanBP2-NT fragment. Nup133 signal labels the cytoplasmic and nuclear rings of the NPC and the localization of the central channel is visualized by Nup62. The magnified framed regions are shown in the corresponding numbered panels (b). The number of single AL-NPC complexes per cell and the number of single AL-NPC complexes in a single AL foci/cluster were quantified in (c, d), respectively. At least 6 cells per condition were analyzed (two sample unpaired T-test, mean ± SD, ***P < 0.001; ****P < 0.0001; N = 3). Scale bars, 3 μm, 0.3 μm (zoomed-in). e, Lysates of HeLa cells expressing HA, HA-RanBP2-NT or HA-RanBP2-NT ΔFG were immunoprecipitated using anti-HA magnetic beads (HA-IP) and analyzed by Western blot (at least 3 independent experiments). f, Auxin-inducible degron (AID) RanBP2 KO DLD-1 cells were treated with DMSO or Auxin for 4h to deplete RanBP2 and cytoplasmic fractions were prepared, separated on a Superose 6 gel filtration column and analyzed by Western blotting (at least 3 independent experiments). Fraction 1 corresponds to the void and the elution of thyroglobulin (670 kDa), g-globulin (158 kDa) and myoglobulin markers 19 kDa are indicated. Green arrows point to fractions preferentially containing single Y-complexes, and red arrows indicate fractions with higher molecular weight, oligomerized Y-complexes. Note that depletion of RanBP2 reduced the abundance of oligomerized Y-complexes (at least 3 independent experiments). g, Auxin-inducible degron (AID) RanBP2 KO DLD-1 cells were treated with Auxin for 4h to deplete RanBP2 and lysates were immunoprecipitated using Nup93 antibody or IgG and analyzed by Western blot (at least 3 independent experiments). h, Lysates of HeLa cells expressing HA or HA-RanBP2-NT were immunoprecipitated using anti-HA magnetic beads (HA-IP) and analyzed by Western blot (at least 3 independent experiments). i, Lysates of HeLa cells were immunoprecipitated using Climp63 antibody or IgG and analyzed by Western blot (at least 3 independent experiments). j-n, Representative images of HeLa cells treated with the indicated siRNAs and analyzed by Western blot (k). Cells were co-labelled with anti-RanBP2 (green) and anti-PDI (red) antibodies (j). The magnified framed regions are shown in the corresponding numbered panels. The dotted line separates the cytoplasmic area with ER sheets proximal to the nuclei where AL-NPCs are enriched in WT cells. Note that AL-NPCs foci enlarge and are localized more distally upon Climp63 downregulation. The number of cells with large AL foci were quantified in (l) where at least 400 cells per condition were analyzed (two sample unpaired T-test, mean ± SD, ns: not significant, ****P < 0.0001; N = 3). The size of cytoplasmic AL foci was quantified in (m) where at least 3500 foci per condition were analyzed (N = 3). The NE intensity of RanBP2 was quantified in (n), where at least 280 cells per condition were analyzed (two sample unpaired T-test, mean ± SD, ns: not significant, ****P < 0.0001; N = 3). Scale bars, 5 μm. o, Representative splitSMLM images depicting NPCs on the nuclear (Nuc) surface and AL-NPCs located in the cytoplasm (Cyt) in HeLa cells treated with Climp63 siRNA. Nup133 signal labels the cytoplasmic and nuclear rings of the NPC, the localization of the central channel is visualized by Nup62. The magnified framed regions are shown in the corresponding numbered panels. Scale bars, 3 μm. p, q, Representative images of HeLa cells treated with the indicated siRNAs. Cells were co-labelled with anti-RanBP2 (green) anti-Nup62(red) antibodies and DAPI (p). The size of nucleus was quantified in (q) where at least 300 cells per condition were analyzed (mean ± SD, ***P < 0.001, unpaired two-tailed t test; N = 3). Scale bars, 5 μm.

Since RanBP2 was required for the assembly of small AL foci and individual AL-NPCs in the cytoplasm (Fig. 4j-l), we next asked how RanBP2 NT regulates this crucial function at the molecular level. Interestingly, structural analysis of the NPC demonstrated that N-termini of five RanBP2 and one Nup93 molecules bind to the overlapping region of two Y complexes (also named Nup107-Nup160 complexes) in the scaffold’s outer ring^45^, and deletion of RanBP2 resulted in an outer ring that lacked Y complexes on the cytoplasmic side^48^. To test if RanBP2 can promote the oligomerization state of scaffold components of AL-NPCs in a similar fashion, we analyzed separated cytoplasmic and nuclear cellular fractions by gel-filtration chromatography (Extended Data Fig. 14g). Deletion of RanBP2 inhibited the formation of oligomers of Y complexes and Nup96-labelled scaffold in the cytoplasm (Fig. 6f) but not in the nucleus (Extended Data Fig. 14h).

Intrinsically disordered FG repeats have been reported to stabilize subcomplexes within NE-NPC^49^, therefore we examined if similar interactions between the RanBP2 NT fragment (aa 1-1171) and other Nups could play a role in AL-NPC assembly. The outer ring of the NPC is mainly composed of the evolutionarily conserved Y complex. We found that the N-terminal fragment of RanBP2 could bind the Y complex, as well as Nup205 and Nup93 (Fig. 6e), which belong to the inner ring and have recently been reported to also reside on the cytoplasmic face of NPCs^45^. The interaction between these Nups with NT RanBP2 was dependent on the FG repeat domain (Fig. 6e), suggesting that it facilitates the assembly of single AL-NPCs. Deletion of RanBP2 did not affect the formation of the Y complex (Extended Data Fig. 15a) both in the cytoplasm and in the nucleus (Extended Data Fig. 15b), but it restricted the binding of the Y complex to Nup93 (Fig. 6g, Extended Data Fig. 15c) specifically in the cytoplasm (Fig. 15c). This suggests that RanBP2 may stabilize interactions between the outer and inner rings of AL-

NPCs in agreement with reported high-resolution NPC structures^45^, and thereby promote AL-NPC assembly in the cytoplasm. Indeed, Nup93 was required for the formation of large AL-foci (Extended Data Fig. 15d-f), and the increase of AL foci abundance by ectopic expression of NT RanBP2 was dependent on Nup93 (Extended Data Fig. 15g, 15h, 15j, 15k,15m). Downregulation of Nup93 also led to the formation of Nup62-positive nuclear foci (Extended Data Fig. 15j), which did not co-localize with other Nups, suggesting that they may not represent AL. Co-localization of NT RanBP2 with Nup133 (Extended Data Fig. 15i) and Nup62 (Extended Data Fig. 15l) in the cytoplasm required Nup93, in line with reports showing that Nup93 helps to connect the central channel to the NPC scaffold^50^. We conclude that the function of the NT part of RanBP2 on AL-NPCs can be attributed to its interactions with Nup93 and its ability to stabilize connections between NPC subcomplexes. Taken together, these results identify the precise molecular mechanism of AL-NPC formation and show that RanBP2 directly drives the interaction between the Y complex and Nup93, as well as the oligomerization of scaffold AL-NPC components.

### Climp63 facilitates the localization of AL-NPCs to ER sheets and their integration with the NE

Our findings on the molecular mechanism of AL-NPC assembly and the direct role of RanBP2 in this process create a unique opportunity to identify additional factors involved in AL biology, in particular those facilitating AL-NPCs insertion into the ER membranes. Since the NT RanBP2 fragment can induce the formation of AL (Fig. 6a-d), we analyzed the proteins interacting with HA-tagged NT RanBP2 (aa 1-1171) compared to HA-tag only, using liquid chromatography-mass spectrometry (LC-MS). Of total 97 proteins specifically bound to NT RanBP2, 20 were associated with the NPC, as expected. Additionally, numerous ER-associated proteins were also identified Extended Data Fig. 16a-c). Among them, Climp63 (also known as CKAP4), a type II membrane protein, emerged as an interesting candidate for the regulation of AL because it has been implicated in maintaining ER morphology, being important for the biogenesis of ER sheets and regulation of their width, and for coordinating the formation and dynamics of ER nanoholes^51,52^. Moreover, its N-terminus binds to MTs, linking the ER to the cytoskeleton^53^. Thus, it is particularly interesting that we indeed detected an interaction between the HA-tagged NT RanBP2 and endogenous RanBP2 and Climp63 (Fig. 6h-i, Extended Data Fig. 17a). Since stringent cell lysis conditions were used for these experiments and several other ER-resident proteins were not found to interact with RanBP2, this argues against nonspecific co-purification of bulk ER membranes and supports a more specific association between RanBP2 and Climp63. In contrast to the interaction of RanBP2 with Nups (Fig. 6e), our current data do not allow us to determine whether the association of Climp63 depends on the FG-rich region of the NT part of RanBP2, and mapping the direct Climp63-RanBP2 binding sites therefore represents an important avenue for future work.

In interphase cells, RanBP2-labeled AL foci preferentially localized to ER regions positive for Climp63 (Extended Data Fig. 17e-g, 17i, j), residing closer to the nucleus relative to the entire ER network (Extended Data Fig. 17h). These Climp63-positive regions likely correspond to ER sheets, in line with published findings^52^. This suggests that, under normal growth conditions, AL may primarily reside within sheet-like regions of the ER rather than in its tubules. The tendency of the ER sheets to localize close to the nucleus may facilitate AL merging with the NE under physiological conditions.

Downregulation of Climp63 (Fig. 6j-o) led to the dispersion of ER sheets throughout the cytoplasm (Fig. 6j), in accordance with previous reports^52^, and strongly altered the morphology and distribution of AL foci. AL foci increased in both abundance (Fig. 6j, 6l) and size (Fig. 6m), relocating to the cell periphery, regions that originally corresponded to ER tubules (Fig. 6j), in Climp63 downregulated cells. Depletion of several ER-shaping and ER-associated proteins including RTN4, ATL1, ATL3, LNPK, KTN1 or P180 did not reproducibly affect AL formation (Extended Data Fig. 17b-d). Instead, their knockdown resulted in highly variable AL phenotypes, likely reflecting general perturbations of ER morphology rather than a specific role in AL biology. Thus, Climp63, and possibly a restricted subset of ER factors, but not ER-associated proteins in general, are specifically involved in the AL pathway.

Additionally, the nuclear intensity of RanBP2 was reduced upon Climp63 downregulation (Fig. 6n). SplitSMLM analysis confirmed that Nup foci in Climp63 downregulated cells correspond to AL (Fig. 6o). One can speculate that the reported Climp63 ability to connect ER to MTs could explain the observed phenotype and Climp63 absence would inhibit MT-mediated transport of AL towards NE. Alternatively, Climp63-mediated regulation of ER dynamics and nanomorphology could explain these effects. These observations also suggest that RanBP2-mediated biogenesis of AL-NPCs in the cytosol can be separated from their proper localization at ER membranes, a process directly dependent on Climp63 and that in the absence of Climp63, cells attempt to compensate these defects by increasing the formation of AL-NPCs. Irrespective of the underlying mechanisms, which could be a topic for future investigations, downregulation of Climp63 decreased Nup levels at the NE (Fig. 6j, 6n) and inhibited nuclear expansion (Fig. 6p, 6q), suggesting that Climp63-mediated ER dynamics is an important prerequisite for the integration of AL-NPCs into the NE in normal cells. Taken together, our results identify the molecular mechanism of AL biogenesis in normal cells, where RanBP2 is crucial for the formation of AL-NPCs in the cytoplasm and Climp63 acts as a factor targeting AL-NPCs to the proper membrane compartment, the ER sheets. These mechanisms ensure nuclear pore assembly and function as well as nuclear growth specifically during G1 phase of the cell cycle.

### AL-dependent NPC assembly is additive to canonical nuclear pore assembly pathways

What is the relationship of the AL-dependent nuclear pore assembly to the canonical assembly pathways acting during two stages of the cell cycle?

Indeed, the postmitotic and the interphase pathways have been primarily described to drive NPC assembly at the NE^26^. Interestingly, in contrast to RanBP2, downregulation of essential upstream components ELYS (postmitotic pathway) or Nup153 (interphase pathway) did not inhibit AL formation but instead strongly induce them (Extended Data Fig. 8d-f)^24^. Our measurements in G1 cell cycle stage estimated that the contribution of AL to pore assembly is substantial, accounting for 22-45% of total nuclear NPCs, suggesting that somatic cells may utilize an additional assembly pathway governed by RanBP2.

To test this hypothesis, we analyzed the complementarity of RanBP2-driven AL pathway with the two canonical NPC assembly pathways. Downregulation of ELYS or Nup153 reduced NE-associated Nup levels and NPC density in the nucleus (Fig. 7a-e) in HeLa cells, as expected. Strikingly, additional knockdown of RanBP2 further exacerbated these effects (Fig. 7a-e). The same results were obtained in DLD-1 cells where RanBP2 was depleted for only 4 hours using auxin-inducible degradation (Fig. 7f-j). These findings suggest that RanBP2-dependent AL do not act redundantly with the classical NPC assembly pathways but instead contribute independently to the biogenesis of a substantial pool of NPCs at the NE. Taken together, mammalian somatic cells may utilize three distinct nuclear pore assembly pathways, in contrast to the two previously described ^26,54–56^.

**Fig. 7.**
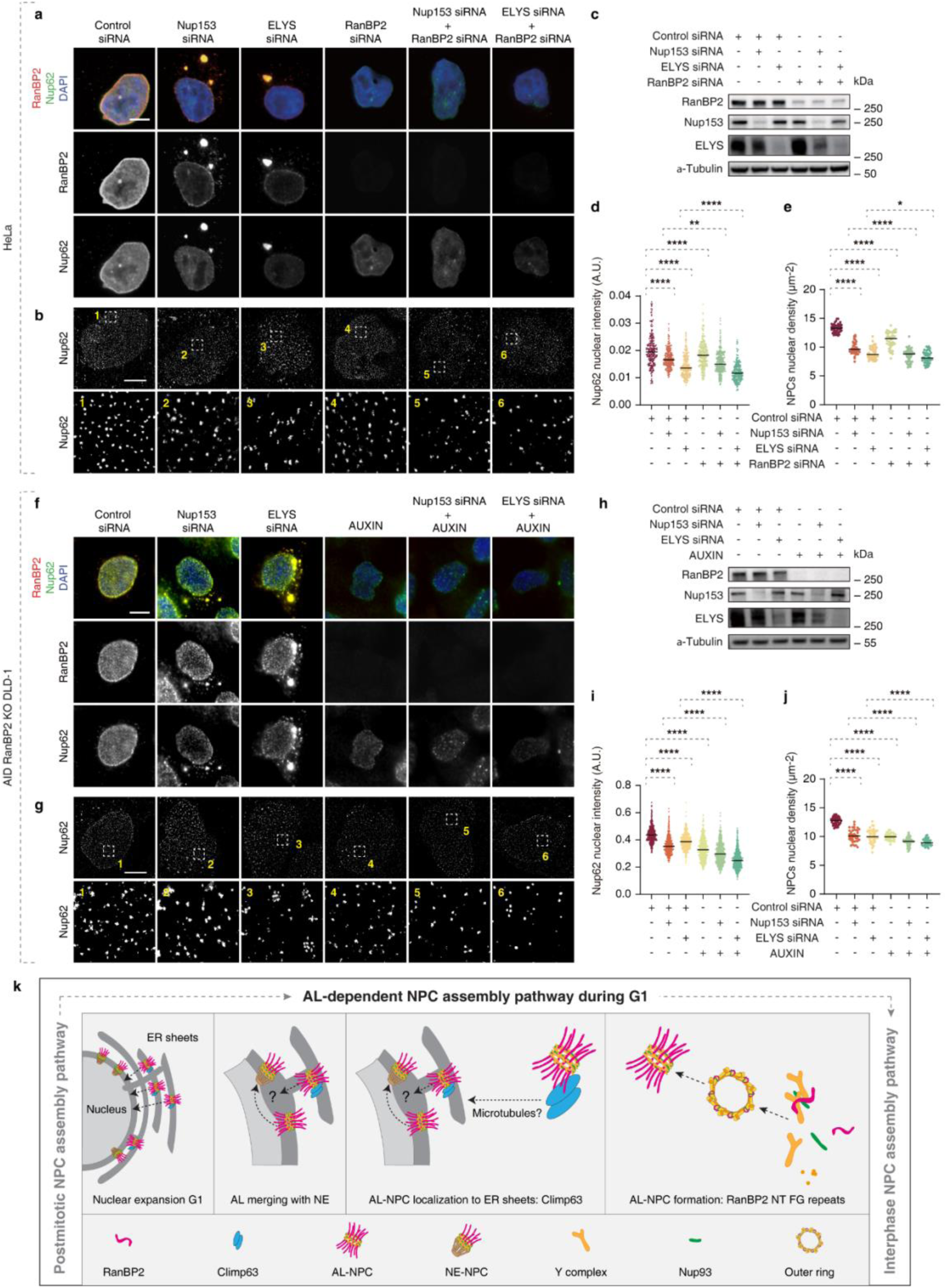
**AL-dependent NPC assembly is additive to canonical nuclear pore assembly pathways.** a-e, Representative images of HeLa cells treated with the indicated siRNAs (48h). Cells were co-labelled with anti-Nup62 (green) and anti-RanBP2 (red) antibodies (a) for immunofluorescence microscopy or labelled with anti-Nup62 for SMLM immunofluorescence (b). (c) Samples from indicated conditions were analyzed by Western blot. The intensity of Nup62 in the NE was quantified in (d), and at least 200 cell per condition were analyzed (one-Way ANOVA test, mean ± SD, **P < 0.01; ****P < 0.0001; N = 3). Scale bars, 5 μm. The nuclear density of NPCs (Nup62) in cells was quantified in (e) (one-Way ANOVA test, mean ± SD, P < 0.05; ****P < 0.0001; counted 50 cells per cell line). Scale bars, IF, 5 μm; SMLM, 3 μm. f-j, Representative images of Auxin-inducible degron (AID) RanBP2 KO DLD-1 cells treated with the Auxin or indicated siRNAs (48h). Cells were co-labelled with anti-Nup62 (green) and anti-RanBP2 (red) antibodies (f) for immunofluorescence microscopy or labelled with anti-Nup62 for SMLM immunofluorescence (g). (h) Samples from indicated conditions were analyzed by Western blot. The intensity of Nup62 in the NE was quantified in (i), and at least 200 cell per condition were analyzed (one-Way ANOVA test, mean ± SD, ****P < 0.0001; N = 3). Scale bars, 5 μm. The nuclear density of NPCs (Nup62) in cells was quantified in (j) (one-Way ANOVA test, mean ± SD, ****P < 0.0001; counted 50 cells per cell line). Scale bars, IF, 5 μm; SMLM, 3 μm. k. A hypothetical model depicting the universal function and direct biogenesis mechanisms of Annulate Lamellae (AL). AL containing few AL-NPCs are present in the cytoplasm in normally proliferating somatic cells and play a crucial role in supplying the NE with new nuclear pores. This function is essential for nucleocytoplasmic transport and for nuclear expansion during the G1 phase of the cell cycle. AL-NPCs on ER membranes can fuse with NE-NPCs to become incorporated into NE during G1 and possibly remodeled to NE-NPCs. The N-terminal part of RanBP2 is required for the formation of AL-NPCs in the cytosol across cellular models and conditions. It ensures oligomerization of NPC scaffold components (Y-complexes) specifically in the cytoplasm. Climp63 drives the insertion of pre-made AL-NPCs into ER sheets which are more proximal to the NE. This mechanism is crucial for AL merging with the NE and for nuclear expansion and constitutes a distinct NPC assembly pathway during G1 phase of normal cell cycle.

## Discussion

### Annulate lamellae (AL) exist in somatic cells under normal growth conditions

In this study, we demonstrated a universal function and direct biogenesis mechanism of Annulate Lamellae, a mysterious cellular structure known for decades to exist in the cytoplasm^12,13^ (Fig. 7k). Using splitSMLM^31^, an advanced super-resolution microscopy and spectral demixing method for colocalization and image analysis, we show that AL are more abundant than previously anticipated and exist in the cytoplasm of a variety of somatic cells under physiological conditions. Our analysis provides unique insights into the biogenesis of AL, which is a key cellular organelle involved in the regulation of nuclear function. Compared to healthy state, dysregulation of AL biogenesis occurs in diseased cells, where AL accumulate and cluster in the cytoplasm as exemplified by our data in the fragile-X syndrome patient-derived samples (Fig. 1e). Under physiological conditions during the G1 cell cycle stage, AL merge with the NE and supply it with pre-formed NPCs, a process that is particularly active during periods of rapid nuclear growth (Fig. 7k). Inhibition of AL merging with the NE, either by blocking their MT-based movement (Fig. 3d-f) or by interfering with AL biogenesis through RanBP2 deletion (Fig. 4n-s) or by preventing their Climp63-mediated localization to ER membranes (Fig. 6j-q), reduces the density of nuclear pores and inhibits nuclear expansion during G1. These findings identify a specific physiological situation that drives the use of AL in normal cells, which were not amenable to AL analysis previously.

Our data may also explain why, in the past, AL have been commonly observed in rapidly developing, cancerous, or challenged cells, which are likely to depend more crucially on rapid nuclear growth ^3–11^. Under these conditions, cells might actively increase their cytoplasmic AL pool by upregulating RanBP2-dependent mechanisms. In addition, our findings show that AL vary widely in size and abundance and are highly dynamic undergoing continuous fusion and fission prior to the merging with the NE (Fig. 2d, 2e). This dynamic behavior may further explain why previous static analyses failed to detect AL in somatic cells. Given variable abundance of AL in different cell types (Fig. 1f), we cannot exclude the possibility that they may rely on the AL pathway to varying degrees.

### AL-dependent NPC assembly during G1 constitutes a distinct pathway in somatic cells

Blocks of AL structures inserted into the NE have been previously observed in *Drosophila* embryos^7^, while our study demonstrates that this process is evolutionarily conserved and operates in somatic cells in general. AL have been also implicated in merging with the NE in postmitotic cells^15^, however, in contrast to our work, this phenomenon seemed independent of ER membranes, and rather involved direct interactions with decondensing chromosomes during late anaphase. Time-resolved analysis of AL dynamics allowed us to estimate the net contribution of AL-NPCs to the NE that accounts for more than 900 new pores per one G1 phase of the cell cycle, representing a substantial portion (22-45%) of the total present NE-NPCs (Fig. 2d, 2E). Analysis of the NPC assembly kinetic revealed that postmitotic NPC assembly is completed within 15 minutes, rapidly adding around two thousand NPCs, while by contrast, interphase NPC assembly is slower and more sporadic, plateauing after 100 minutes, with unknown NPC assembly rate^57^. In comparison, our AL-NPC insertion data show a continuous rate of 1.5 NPCs per minute throughout entire G1, as supported by the persistent presence of AL foci from early G1 through mid-to-late G1/S (Fig. 2a, 2b) and by photoconversion experiments (Fig. 2f, 2g). Thus, although slower than the postmitotic assembly, AL-NPC insertion occurs at a rate comparable to canonical interphase assembly and represents a kinetically significant and sustained mechanism of NPC addition.

Strikingly, AL-driven NPC assembly acts complementarily to the two canonical NPC assembly pathways and the effects on pore density observed upon simultaneous depletion of RanBP2 and ELYS (postmitotic pathway) or Nup153 (interphase pathway) are additive (Fig. 7a-j). Thus, RanBP2- and AL-dependent NPC increase occurs through an independent mechanism (Fig. 7a-j). Supporting this, our data identify a RanBP2 mutant (ΔFG region mutant) that clearly separates its AL-specific function from its other roles at the NE and in the cytoplasm. Taken together, our study positions the AL-insertion mechanism as a distinct, possibly third NE-NPC biogenesis pathway that operates specifically during interphase and acts complementarily to the two well-described pathways in higher eukaryotes^26,54–56^. In the future, it will be important to understand why downregulation of factors responsible for postmitotic and interphase NPC assembly results in the accumulation of AL and how these defects are linked to the regulation of RanBP2.

Our findings on the existence of small AL structures represent an important first step toward future studies aimed at understanding how normal somatic cells acquire pre-assembled AL-NPCs during nuclear growth. During development, the simultaneous insertion of numerous AL-NPCs (arranged within several parallel ER membranes in the cytoplasm) into the nuclear envelope (NE) involves the formation of large NE openings^7^, which likely serve to avoid topological constraints associated with AL insertion. It is unknown if a similar mechanism operates in somatic cells. Studies by Otsuka and Ellenberg^58^ indicate that the reforming NE is fenestrated at mitotic exit and largely seals by early G1, however, a small fraction of openings may plausibly persist into G1. Such residual discontinuities could, in principle, serve as entry sites for AL-derived NPCs or for membrane fusion events contributing to NE growth, which should be studied in the future.

Our splitSMLM analysis reveals that small, symmetric AL-NPCs laterally merge with asymmetric NE-NPCs. Some of these fusion events are accompanied by the formation of small NE openings containing ER membranes, with both AL-NPCs and NE-NPCs positioned adjacent to each other, without disrupting the underlying lamina components (Fig. 3a-c). While a recently proposed model suggests that the GTPase Ran remodels AL-NPCs into asymmetric, functional NPCs in fly embryos^34^, it will be crucial in future work to define the exact remodeling steps that occur during AL membrane insertion into the NE in somatic cells, ideally using high-resolution ultrastructural approaches. Although our data support the insertion of intact AL structures into the NE, we cannot yet rule out the possibility that AL-NPCs undergo partial disassembly prior to insertion, or that AL act primarily as donors of NPC subcomplexes rather than inserting as complete structures.

Interestingly, the bleb-like AL-NE intermediates observed on the NE by our splitSMLM analysis may resemble NE herniations described in *Saccharomyces cerevisiae* and in human cells carrying mutations in Brl1/6 or TorsinA^59–63^. These factors have been implicated in interphase NPC assembly, possibly through roles in membrane fusion and lipid homeostasis at the NE. Whether Brl1/6 or TorsinA also contribute to AL insertion into the NE and the regulation of AL-NPCs remains an important avenue for future research.

Yet, our work not only demonstrates that small AL are clearly beneficial for cellular homeostasis by preserving nuclear function during G1, but also lays a solid foundation for important future studies on molecular topology of nuclear pore formation to the NE.

### The molecular mechanisms of AL biogenesis in somatic cells

In addition to elucidating the universal cellular function of AL during G1, the direct molecular mechanisms of AL formation identified in our study explain how AL-NPCs are made and how they could localize to the proper compartment of the ER (Fig. 7k). We show that RanBP2 is indispensable for the assembly of cytosolic AL-NPCs across cell types and conditions, serving as an assembly platform that specifically regulates the oligomerization of Y complexes within the AL-NPC scaffold. While this process may involve other structurally important Nups, such as Nup93 (Extended Data Fig. 15), the N-terminal region of RanBP2 containing FG repeats is required for AL biogenesis. Intrinsically disordered FG repeats have been shown to stabilize subcomplexes within NE-NPC^49^, providing a possible explanation for their function on AL-NPCs. The role of cohesive FG repeats could also help to understand previous observations in fly embryos, where RanBP2 and intact microtubules were implicated in recruiting various condensed precursor Nup particles, predicted to lead to AL biogenesis^10^. Although we occasionally observe cytoplasmic RanBP2 foci under normal conditions (Fig. 3c), we are unable to clearly classify them as membrane-free. Interestingly, our results reveal a spatial association of AL to the microtubule network (Extended Data Fig. 7b) and RanBP2 has previously been reported to stabilize MT bundles and recruit dynein motor protein^64,65^. Whether RanBP2 mediates the interaction between AL and microtubules and if this process involves the formation of Nup condensate particles that facilitate AL assembly in somatic cells, remain important questions for the future. Here, we provide direct evidence for the role of RanBP2 in AL biogenesis and clarify the detailed molecular requirements for AL-NPC formation that is functionally analogous but mechanistically distinct from that described in fly oocytes.

On the membrane side, we identify Climp63 as a key factor localizing AL-NPCs to ER sheets proximal to the nucleus, a process important for their subsequent integration into the NE and nuclear expansion. To our knowledge, this is the first evidence directly linking AL to specific ER substructure so far. By analysing the interaction and function of several other ER factors, we support the idea that Climp63, and possibly a restricted subset of ER proteins, but not ER-associated factors in general, is specifically involved in the AL pathway. Since Climp63 was shown to ensure not only the formation of ER sheets but also the biogenesis of ER nanoholes^51,52^, which were discussed to be similar in size to that of nuclear pores^51^, it would be exciting to study the roles of ER dynamics and nanomorphology in AL biogenesis. It would also be important to understand if the reported role of Climp63 in connecting ER to MTs could explain its role in positioning AL in proximity to the NE.

One of the most abundant proteins in the AL proteome belongs to the RGPD family (Extended Data Fig. 16a), which shares significant sequence similarity with the N-terminal domain of RanBP2. Although RGPD proteins are predicted to localize to the Golgi apparatus, they may retain MT-binding capacity^66^ and it will be interesting to study this protein family in the context of AL biology. Likewise, a possible role of NE-resident proteins SUN1 and SUN2, which we found to co-localize with AL foci (Extended Data Fig. 5b) and which are components of LINC complexes^67^, could be during early steps of NPC interphase assembly^68^, which opens possibilities for future investigations.

Undoubtedly, our findings not only clarify the fundamental mechanisms of AL biogenesis but also create an important ground for future investigations aiming to identify AL-specific factors, which could be utilized by pathologically challenged cells such as during malignancy. Indeed, maintaining NPC function is crucial for normal cellular physiology, as disruptions in nucleocytoplasmic transport, mis-localization of Nups, and elevated levels of AL have been observed in various diseased and infected cells^23,25,46,69–74^. The biological principles of AL biology identified by our study may create a basis for future approaches to restore NPC homeostasis in human diseases such as fragile-X syndrome and numerous other neurological and neurodegenerative diseases as well as cancer ^14^.

## Acknowledgements

We thank members of the Sumara, I. and Ricci, R. laboratories for helpful discussions on the manuscript and IGBMC facilities for their help. We are grateful to Valérie Doye, Bernhard Hempoelz and participants of the “Rembo” workshop for stimulating discussions on data interpretation. We are grateful to Arnaud Echard and Anakine Prizins for helpful support on photoconversion experiments and comments on the manuscript. We acknowledge the Virus and Molecular Biology platform of the IGBMC for the lentivirus production. We acknowledge the IGBMC imaging center, member of the national infrastructure France-BioImaging supported by the French National Research Agency (ANR-10-INBS-04).

Lin, J. Liu X. and Liao, Y. were supported by PhD fellowships from the China Scholarship Council (CSC). Agote-Aran, A., was supported by PhD fellowship from the IMCBio graduate school and a fellowship from the “Ligue Nationale Contre le Cancer”. Cloarec M. was supported by PhD fellowship from the doctoral school University of Strasbourg.

Andronov, L., Schoch, R. and Klaholz, B. P. acknowledge support by Institut National du Cancer (INCa) and by the French Infrastructure for Integrated Structural Biology (FRISBI) ANR-10-INSB-05-01, Instruct-ERIC and iNEXT-Discovery. Golzio, C. is a permanent INSERM investigator. Lemée, M.V. is a doctoral fellow supported by EUR IMCBio funds and Fondation de France (WB-2022-45868). This work was funded by Agence Nationale de la Recherche under the projects (ANR-22-CE12-0011). Research in Sumara, I. laboratory was supported by the grant ANR-10-LABX-0030-INRT, a French State fund managed by the Agence Nationale de la Recherche (ANR) under the frame program Investissements d’Avenir ANR-10-IDEX-0002-02, IGBMC, CNRS, ARC, INCa (PLBIO 2022-082), ANR (AAPG2022, NICE4Nups) and AFM-Telethon.

## Author contributions

The role of RanBP2 in human cytosolic NPCs was initially discovered by A.A.A. and confirmed by J.L., and the research plan and the conceptualization of the results were based on discussions between J.L. and I.S.. J.L. performed most of the experiments and analyzed the data. Y.L. helped with experiments on UBAP2L, L.A. and R.S. helped with splitSMLM analyses, P.R. helped with CLEM, R.Z. helped with gel-filtration chromatography, M.V. L generated the hiPSC derived neurons, E. G. helped with photoconversion experiment, X.L and M.C. helped with FB789 and MRC5 cell lines and M.C. helped with experiments on Climp63. C.K. helped with the reagents and materials. M.R., Y.S., C.G. and B.P.K. helped with the design of the experiments and supervision. I.S. supervised the project and J.L. and I.S. wrote the manuscript with input from all authors.

## Competing interest

The authors declare no competing interests.

## Additional information

The supplementary materials consist of Supplementary Table 1, Supplementary Videos 1-10 and Extended Data Figures 1-17.

## Methods

### Antibodies

The following primary antibodies were used: rabbit monoclonal anti-Nup133 (Abcam, ab155990), mouse monoclonal anti-Nup133 (E-6) (Santa Cruz Biotechnology, sc-376763), mouse monoclonal anti-Nucleoporin p62 (BD Biosciences, 610497), rabbit polyclonal anti-RanBP2 (Abcam, ab64276), rabbit polyclonal anti-Nup214 (Abcam, ab70497), rabbit polyclonal anti-Nup85 (Bethyl, A303-977A), mouse monoclonal anti-Pericentrin 1 (D-4) (Nup85) (Santa Cruz Biotechnology, sc-376111), rabbit monoclonal anti-SEC13 (R&D systems, MAB9055), rabbit polyclonal anti-ELYS (Bethyl, A300-166A), mouse monoclonal Nup205 (H-1) (Santa Cruz Biotechnology, sc-377047), mouse monoclonal anti-Nup93 (E-8) (Santa Cruz Biotechnology, sc-374399), rabbit monoclonal anti-Nup98 (C39A3) (Cell Signaling Technology, 2598), rabbit polyclonal anti-Nup96 (Bethyl, A301-784A), rabbit polyclonal anti-POM121 (GeneTex, GTX102128), rabbit polyclonal anti-Nup153 (Abcam, ab84872), rabbit polyclonal anti-CRM1/Exportin 1 (Novus, NB100-79802), mouse monoclonal anti-NTF97/Importin beta (Abcam, ab2811), mouse monoclonal anti-Ran (BD Biosciences, 610340), rabbit monoclonal anti-SUN1 (Abcam, ab124770), rabbit monoclonal anti-SUN2 (Abcam, ab124916), rabbit polyclonal anti-Lamin A (C-terminal) (Sigma, L1293), rabbit polyclonal anti-Lamin B1 (Abcam, ab16048), rabbit monoclonal anti-Nesprin1 (Abcam, ab192234), rabbit polyclonal anti-Emerin (Abcam, ab40688), rabbit polyclonal anti-LAP2 (Proteintech, 14651-1-AP), rabbit polyclonal anti-Phospho-Rb (Ser807/811) (Cell Signaling Technology, 9308), mouse monoclonal anti-Cyclin B1 (G-11) (Santa Cruz Biotechnology, sc-166757), rabbit polyclonal anti-Cyclin B1 (GeneTex, GTX100911), rat monoclonal anti-HA (Roche, 11867423001), rabbit polyclonal anti-GAPDH (Sigma, G9545), mouse monoclonal anti-β-Actin (Sigma, A2228), mouse monoclonal anti-mCherry (4B3) (Thermo Scientific, MA5-32977), rabbit polyclonal anti-UBAP2L (Abcam, ab138309), mouse monoclonal anti-FXR1 (Millipore, 03-176), rabbit polyclonal anti-α-Tubulin (Abcam, ab18251), mouse monoclonal anti-α-Tubulin (Sigma, T9026), mouse monoclonal anti-Nuclear Pore Complex Proteins (mAb414) (Abcam, ab24609), rabbit polyclonal anti-LC3B (Novus biological, NB100-2220SS), rabbit polyclonal anti-LC3A (Novus biological, NB100-2331), rabbit polyclonal anti-p62 (GeneTex, GTX100685), rat monoclonal anti-GFP (3H9) (ChromoTek, 3h9-100), rabbit polyclonal anti-GFP (Abcam, ab290), rabbit polyclonal anti-Aurora B (Abcam, ab2254), rabbit polyclonal anti-cyclin A (H-432) (Santa Cruz Biotechnology, sc-751), mouse monoclonal anti-PDI (Abcam, ab2792), mouse monoclonal anti-Climp63 (Enzo, ALX-804-604), rabbit polyclonal anti-CKAP4 (Proteintech, 16686-1-AP), rat monoclonal α-tubulin-conjugated to Alexa Fluor® 647 (Abcam, ab195884), rabbit polyclonal anti-RTN4/NOGO (Proteintech, 10950-1-AP), rabbit polyclonal anti-RRBP1 (Proteintech, 22015-1-AP), rabbit polyclonal anti-KTN1 (Proteintech, 19841-1-AP), rabbit polyclonal anti-ATL1(OriGene, ta332595), rabbit polyclonal anti-ATL3 (Proteintech, 16921-1-AP), rabbit polyclonal anti-LNPK (Atlas Antibodies, HPA014205).

Secondary antibodies used were the following: goat anti-mouse IgG antibody (HRP) (GeneTex, GTX213111-01), goat anti-rabbit IgG antibody (HRP) (GeneTex, GTX213110-01) and goat anti-rat IgG antibody (HRP) (Cell Signaling Technology, 7077S), goat polyclonal anti-mouse AF647 (Thermo Fisher Scientific, A-21236), goat polyclonal anti-mouse AF568 (Thermo Fisher Scientific, A-11031), goat polyclonal anti-mouse AF555 (Thermo Fisher Scientific, A-21424), goat polyclonal anti-mouse AF488 (Thermo Fisher Scientific, A-11029), goat polyclonal anti-rabbit AF647 (Thermo Fisher Scientific, A-21245), goat polyclonal anti-rabbit AF568 (Thermo Fisher Scientific, A-11036), goat polyclonal anti-rabbit AF555 (Thermo Fisher Scientific, A-21429), goat polyclonal anti-rabbit AF488 (Thermo Fisher Scientific, A-11034), goat polyclonal anti-mouse CF680 (Sigma, SAB4600199), goat polyclonal anti-chicken CF660C (Sigma, SAB4600458), goat polyclonal anti-rat CF660C (Sigma, SAB4600193).

### Plasmid and siRNA transfections

The following plasmids were used: pX330-P2A-EGFP/RFP and pUC57 was a generous gift from Zhirong Zhang (Romeo Ricci laboratory IGBMC). pQCXIP-mScarlet-ER was a generous gift from Julie EICHLER ^75^ (Catherine-Laure Tomasetto laboratory IGBMC). pEF-HA-MCS2, pEF-HA-RanBP2 (1-3224), pEF-HA-RanBP2 (1-2448), and pEF-HA-RanBP2 (1-1171) were generous gifts from Ralph Kehlenbach (University of Göttingen). pEF-HA-RanBP2 (1-832) and pEF-HA-RanBP2 (FL without FG repeat) was obtained by inserting the target sequence, obtained by PCR using pEF-HA-RanBP2 as a template, into pEF-HA-MCS2. pCDNA 3.1 3XHA NUP358 FL (LIQIML) was generous gifts from André Hoelz (California Institute of Technology). The siRNA-resistant RanBP2 plasmid was obtained by PCR and DpnI (Thermo) treatment. NLS-mCherry-LEXY (pDN122) (Plasmid #72655) and mEos2-N1 (Plasmid #54662) was purchased from Addgene. The primers used for plasmid generation are provided in the Supplementary Table 1.

The following siRNA oligonucleotides were used: Non-targeting individual siRNA (Dharmacon), FXR1 siRNA (Dharmacon), UBAP2L siRNA (Dharmacon), RanBP2 siRNA (Dharmacon), Nup153 siRNA (Dharmacon), ELYS siRNA (Dharmacon), POM121 siRNA (Life Technologies), Nup214 siRNA (Eurogenetec), Nup93 siRNA (Horizon), CLIMP-63(Dharmacon), RTN4 (Horizon), p180/RRBP1 (Horizon), KTN1 (Horizon), ATL1 (Horizon), ATL3 (Horizon), LNPK (Horizon). The sequences of siRNA are provided in the Supplementary Table 1.

X-tremeGENE^TM^ 9 DNA Transfection Reagent (Roche) was used to perform transient plasmid transfections according to the manufacturer’s instructions. Lipofectamine™ RNAiMAX Transfection Reagent (Invitrogen) was used to deliver siRNAs for gene knock-down (KD) according to the manufacturer’s instructions.

### Generation of cell lines and cell culture

HeLa Kyoto (human cervix carcinoma) cells and U2OS (human bone osteosarcoma) cells were purchased from ATCC. HeLa Kyoto GFP-Nup107 derived stable cell were purchased from CSL cell bank. AID-NUP358 DLD-1 (colorectal adenocarcinoma epithelial cells) cells and AID-NUP358 HTC116 (human colon cancer cells) cells were generous gifts from Mary Dasso (National Institutes of Health). hTERT-RPE1 (Human retinal pigment epithelial-1) cells was a generous gift from Juliette Godin (IGBMC). FB789 (Human Fibroblast) cells was a generous gift from PROIETTI Luca (IGBMC). MRC5 (Human Fibroblast) cells was a generous gift from SEROZ Thierry (IGBMC).Control human fibroblasts and FXS patient-derived fibroblasts were previously reported^25^.

mCherry-Nup133 and mEos2-Nup133 knock-in HeLa Kyoto cell lines were generated using CRISPR/Cas9 genome editing system as described previously^76^. The guide RNAs (gRNA) were designed as described previously^77^, and cloned into pX330-P2A-EGFP through ligation using T4 ligase (New England Biolabs). The donor constructs used as templates for homologous recombination to repair the Cas9-induced double-strand DNA breaks were generated by cloning the Nup133 DNA fragments upstream and downstream of the CRISPR target sequences and the sequence for mCherry or mEos2, assembled using ExonucleaseⅢ (Takara) method, into a pUC57 vector. HeLa Kyoto cells were transfected for 24h and GFP positive cells were collected by FACS (BD FACS Aria II), cultured for 2 days and seeded with FACS into 96-well plates. Single-cell clones of mCherry-Nup133 knock-in or mEos2-Nup133 knock-in were validated by Western blot and sequencing of PCR-amplified targeted fragment by Sanger sequencing (GATC). The sequences of gRNA and the primers used for plasmid generation are provided in the Supplementary Table 1.

UBAP2L knock-out (KO) in mCherry-Nup133 knock-in HeLa Kyoto cell lines were generated using CRISPR/Cas9 genome editing system as described previously^76^. Two guide RNAs (gRNA) were designed using the online software Benchling (https://www.benchling.com/), and cloned into pX330-P2A-EGFP/RFP through ligation using T4 ligase (New England Biolabs). mCherry-Nup133 knock-in HeLa Kyoto cells were transfected for 24h and GFP and RFP double positive cells were collected by FACS (BD FACS Aria II), cultured for 2 days and seeded with FACS into 96-well plates. UBAP2L KO single-cell clones were validated by Western blot and sequencing of PCR-amplified targeted fragment by Sanger sequencing (GATC). The sequences of gRNA are provided in the Supplementary Table 1.

All cell lines were cultured at 37°C in 5 % CO2 humidified incubator. HeLa Kyoto and its derived cell lines were cultured in Dulbecco’s Modified Eagle Medium (DMEM) (4.5 g/L glucose) supplemented with 10 % fetal calf serum (FCS), 100 U/ml Penicillin + 100 µg/ml Streptomycin. U2OS cell lines were cultured in DMEM (1 g/L glucose) supplemented with 10 % FCS, Non-Essential Amino Acids + Sodium Pyruvate 1 mM + Gentamicin 40 µg/mL. DLD-1 cell lines were cultured in DMEM (1 g/L glucose) supplemented with 10 % FCS, 100 U/ml penicillin, and 100 µg/ml streptomycin. HTC116 cell lines were cultured in McCoy’s 5A medium + 10 % FCS

+ 100 U/ml penicillin, and 100 µg/ml streptomycin. hTERT-RPE1 cells were cultured in Dulbecco’s modified Eagle medium (DMEM) F-12 supplemented with 10% FCS, 0.01 mg/ml hygromycin B. FB 789 cell lines were cultured in MEM w/Earles’s salts supplemented with 15 % FCS, + Vitamines + AANE + Gentamicine 40 µg/ml + Penicillin 100 UI/ml + Streptomycin 100 µg/ml. MRC5 cell lines were cultured in DMEM (1 g/L glucose) supplemented with 10 % FCS, + Gentamicine 40 µg/ml.

### Culture and differentiation of hiPSCs into neurons

Human induced pluripotent stem cells (hiPSCs, GM8330-8; kindly given by M. E. Talkowski) were cultured in mTeSR1, following the manufacturer’s instructions, and maintained on Cultrex-BME. hiPSCs were cultured at 37°C with 5% CO₂ under humidified conditions. Colonies were passaged every 3 to 4 days by manual cutting.

For the generation of inducible rtTA/Ngn2 hiPSC lines, Lentiviral vectors (PCW57) containing the human NGN2 gene cassette under the control of the TRE promoter, along with a puromycin resistance gene linked to rTetR via a T2A sequence under a constitutive promoter, were used to transduce hiPSCs. The construct was cloned and packaged into lentiviruses by the Molecular Biology Platform of the IGBMC, hereafter named TetO-NGN2 lentivirus. hiPSCs were dissociated using Accutase and seeded in 12-well plates at a density of 6 × 10⁴ cells per well in mTeSR1 supplemented with 10 μM Rocki (STEMCell), and incubated overnight at 37°C. The next day, the medium was removed, and the cells were incubated for 6 hours in DMEM/F12 containing 8 μg/ml Polybrene, 10 μM Rocki, and 4 × 10⁴ TU of TetO-NGN2 lentivirus. After 6 hours, cells were washed with DPBS (Thermo Fisher) and returned to mTeSR1 at 37°C and 5% CO₂. Puromycin selection began on day 1 post-transduction, with 0.5 μg/ml from day 1 to 2, 1 μg/ml from day 3 to 4, and then 0.5 μg/ml again from day 5 to 7, with daily medium changes using mTeSR1. Selected hiPSC colonies were subsequently expanded in new Cultrex-BME-coated plates after Accutase dissociation and maintained under standard conditions.

For the differentiation of rtTA/Ngn2 hiPSCs into induced neurons, the differentiation was adapted from reported protocol without addition of astrocytes^78^. On day 0, TetO-NGN2 hiPSCs were dissociated with Accutase and seeded at a density of 4 × 10⁵ cells/cm² on Poly-L-Ornithine (50 μg/cm²) + Laminin (7 μg/cm²)-coated petri dishes and in mTeSR1 supplemented with 10 μM Rocki and 3 μg/ml doxycycline. On day 1, media was replaced with DMEM/F12 + Glutamax 1X + NEAA 1X + B27 1X + N2 1X + doxycycline 3 μg/ml. On day 3, media was replaced with Neurobasal + Glutamax 1X + NEAA 1X + B27 1X + NT-3 (10 ng/mL) + BDNF (10 ng/mL) + doxycycline 3 μg/ml + cytosine β-D-arabinofuranoside (AraC) 5 μM. From day 5 to day 20, media was half-changed each 3-4 days with Neurobasal + Glutamax 1X + NEAA 1X + B27 1X + NT-3 (10 ng/mL) + BDNF (10 ng/mL) + doxycycline 3 μg/ml. Doxycycline was maintained until day 14 of differentiation. Neurons were fixed/immunostained at 20 DIV.

### Cell treatments and cell cycle synchronization

To induce the formation of the cytoplasmic AL foci by microtubule depolymerization, cells were incubated with 10 μM nocodazole (Sigma M1404-50MG) in culture media for 90 min at 37°C. To inhibit autophagy, cells were incubated with 30 nM Bafilomycin (Sigma, B1793-10UG) in culture media for 12h at 37°C. Cells were synchronized in early G1 by the addition of thymidine (Sigma, T1895) at 2 mM for 16h. For cell cycle synchronizations double thymidine block and release (DTBR) protocol was used. Cells were treated with 2 mM thymidine for 16h, washed out (three times with warm thymidine-free medium), then released in fresh thymidine-free culture medium for 8h, treated with 2 mM thymidine for 16h again, washed out, and then released in fresh thymidine-free culture medium for 9h (mitosis) or 12 hours early G1 phase.

### Immunofluorescence

Cells were plated on 9-15 mm glass coverslips (Menzel Glaser) in 12- or 24-well tissue culture plates. For immunofluorescence, cells were washed twice with PBS before being fixed with 4 % paraformaldehyde (PFA, Electron Microscopy Sciences 15710) in PBS for 15-20 min at room temperature. Cells were washed three times for 5 min in PBS before permeabilization for 5 min with 0.5 % NP-40 (Sigma) in PBS. Cells were washed three times for 5 min in PBS before blocking for 1h at room temperature with 3 % BSA in PBS-Triton 0.01 % (Triton X-100, Sigma, T8787). Cells were then incubated with primary antibody in blocking buffer (3 % BSA in PBS-Triton 0.01 %) for 1h at room temperature. Cells were washed three times with PBS-Triton 0.01 % gently shaking for 10 min before incubated with secondary antibody in blocking buffer for 1h at room temperature in the dark. After incubation, cells were washed 3 times with PBS-Triton 0.01 % for 10 min each time with gentle shaking in the dark, and then glass coverslips were mounted on glass slides using MOWIOL containing 0.75 μg/ml DAPI (Calbiochem) and imaged with 100X, 63X or 40X objectives using Zeiss epifluorescence microscope or confocal microscope Leica Spinning Disk Andor/Yokogawa.

### Live-cell imaging

For live-cell microscopy, cells were grown on 35/10 mm 4 compartment glass bottom dishes (Greiner Bio-One, 627871) or μ-Slide 8 well glass bottom (Ibidi, 80827). Before photography, cells were treated with SiR-DNA (Spirochrome, SC007) according to the manufacturer’s instructions. Live cell microscopy was performed using a confocal microscope Leica/Andor/Yokogawa spinning disc, Leica CSU-W1 spinning disc, or Nikon PFS spinning disc with a 40X or 63X objectives with Live Data Mode equipped with automated temperature.

In particular, for AL foci’ dynamics assays under normal growth conditions, HeLa Kyoto GFP-Nup107 were analyzed by Leica CSU-W1 spinning disk (63X NA 1.4 oil objective) for 9h. Z-stacks (10 μm range, 1 μm step) were acquired every 5 min and Videos were made with maximum intensity projection images for every time point shown at speed of 7 frames per second. For AL foci’ dynamics assays after microtubule depolymerization, HeLa Kyoto GFP-Nup107 cell were arrested in S phase by thymidine block and microtubule depolymerization and nucleoporin aggregation were induced by 10 μM nocodazole addition. For AL foci’ dynamics assays under FXR1 knockdown (KD) or UBAP2L KD conditions, HeLa Kyoto GFP-Nup107 derived stable cell were treated with the indicated siRNAs for 48 h before imaging.

For rapid super-resolution live-cell imaging, HeLa Kyoto GFP-Nup107 were analyzed by Nikon PFS spinning disk (100X NA1.4 oil objective) for 15 min using Live-SR mode. Z-stacks (7 μm range, 1 μm step) were acquired every 3 s and Videos were made with maximum intensity projection images for every time point shown at speed of 7 frames per second. For AL foci and microtubules dynamics assays, cells were treated with SiR-tubulin (Spirochrome, SC002) according to the manufacturer’s instructions.

For AL foci and ER’ dynamics assays, HeLa Kyoto GFP-Nup107 were transfected with the pQCXIP-mScarlet-ER for 24 h and analyzed by Nikon PFS spinning disk (100X NA1.4 oil objective) for 5h where images were acquired every 20s (without Z-stacks) or were analyzed for 9h where images were acquired every 2 min (without Z-stacks).

For photoactivatable nucleocytoplasmic shuttling, AID-NUP358 DLD-1 were transfected with the NLS-mCherry-LEXY (pDN122) for 24h and analyzed by Leica Sp5 confocal inverted microscope (DMI6000) system (40X NA1.3 oil objective). The circular region of interest (ROI; ∼38 µm2) was placed onto single, selected cells. The ROI was scanned with a 458-nm laser beam for 10 min following a 20-min dark-recovery phase. The mCherry signal was imaged in parallel every 10s for 30 min using the 561-nm laser line for excitation.

For photoconversion assays, HeLa Kyoto mEos2-Nup133 were analyzed by Nikon PFS spinning disk (100X NA1.4 oil objective). The modular part developed by the GATACA system was used to set the acquisition of the selected area to be photoconverted with applied repeats of blue light for a period of 200 seconds (typical value, depending on the size of the area) by the 500 mW laser diode at 100%. A 30-second acquisition pre-sequence and with acquisition every 10 seconds in the green and red channels has been perfomrmed. Two post sequences are carried out after acquisition, the first for 30 seconds with an interval every 10 seconds in order to quickly observe the effect of photoconversion and the second sequence for 12 hours every 15 minutes to see the effect over a longer period. The software was set up to use spinning with the 491 nm 50 mW laser diode at 20% power with an acquisition time of 150 ms for the green channel and the 561 nm 50 mW laser diode at 30% power with 150 ms exposure time for the red channel.

### Single molecule localization microscopy

For super-resolution imaging using single molecule localization microscopy (SMLM), cells were plated on 35 mm glass bottom dish with 14 mm micro-well #1.5 cover glass (Cellvis). Samples were prepared as described above for immunofluorescence but were not mounted using MOWIOL and were stored in PBS.

The imaging was performed as previously described, using the splitSMLM method^79^ for multi-color single molecule localization microscopy (SMLM) based on a dichroic image splitter. In brief, the immunolabeled samples were mounted in imaging buffer containing 200 U/ml glucose oxidase (G2133, Sigma), 1000 U/ml catalase (C1345, Sigma), 10 % w/v glucose, 200 mM Tris-HCl pH 8.0, 10 mM NaCl, 50 mM MEA (30080, Sigma), with addition of 2 mM COT (138924, Sigma). The buffer optimizes the performance of the fluorophores AF647, CF660C and CF680, for three-color direct stochastic optical reconstruction microscopy (dSTORM). Imaging was then performed on a modified Leica SR GSD system, with a HCX PL APO 160X/1.43 Oil CORR TIRF PIFOC objective. The acquisitions began with a pumping phase, during which the sample was illuminated with the 642 nm 500 mW fiber laser (MBP Communication Inc.) but the fluorescence was not recorded due to a very high density of fluorophores in a bright state. The image collection started when the density of fluorophores dropped to a level that allowed observation of individual molecules and in case of emitter depletion, the sample was additionally illuminated with a 405 nm diode laser (Coherent Inc.) to increase the emitter density (back-pumping). The resulting emission was split with an Optosplit II (Cairn Research) image splitter, attached to a camera port of the microscope, and containing a Chroma T690LPXXR dichroic mirror. This creates a long and short wavelength channel on the Andor iXon Ultra 897 EMCCD camera attached to the image splitter. Each fluorophore thus shows a characteristic brightness ratio between the two channels, enabling the separation of spectrally different fluorophore species (i.e., spectral demixing)^79^. Localization of the single molecules in both channels was performed using the Leica LAS X software with the “direct fit” fitting method. The localization tables were then exported and further processed using the SharpViSu software^29,80,81^ and the SplitViSu plugin^79^. This accounts for spectral demixing, correction of chromatic errors, drift-correction, and the reconstruction of super-resolution images. A pixel size of 10 nm was used to reconstruct the super-resolution images from 2D-histograms of single-molecule coordinates. For the single-color imaging without image-splitter, the images were recorded with an Andor iXon + EMCCD camera, attached to the second camera port of the microscope. Data was then processed as already described, using the Leica LAS X and SharpViSu software, while omitting the spectral demixing steps in the SplitViSu plugin.

The NPCs area enlarged in the article uses Bilinear interpolation for better observation.

### Correlative light and electron microscopy

Cells were grown on carbon coated sapphire disks (3 mm diameter, 50 mm thickness, Wohlwend GmbH, art. 405) and high pressure frozen (HPM010, AbraFluid) in their culture medium. Freeze substitution was performed in a Leica EM-AFS2 with 0.1 % uranyl acetate in dry acetone for 24h at −90°C. The temperature was then raised to −45°C over 9h (slope 5 degrees / hour) and the samples were further incubated at this temperature for 5h to increase electron contrast. After rinsing in acetone, the samples were infiltrated with increasing concentrations of Lowicryl HM20 (Polysciences Inc.), while raising the temperature to −25°C. Finally, the blocks were polymerized using UV light. After removal of the sapphire disks from the block surface, 300 nm sections were cut parallel to the block surface using an ultramicrotome (UC7, Leica Microsystems). The sections were collected on carbon coated mesh grids (S160, Plano).

Fluorescence imaging of the sections was carried out with a widefield microscope (Olympus IX81), equipped with a 100X 1.40 NA Plan-Apochromat objective, placing the grids in water in a glass bottom dish (Mattek). After light microscopy acquisition, the grids were post-stained with 2 % uranyl acetate in 70 % methanol and Reynold’s lead citrate. Electron microscopy (EM) was performed using a Tecnai F30 transmission electron microscope (Thermo Fisher) at 300kV acceleration voltage. The grid squares that were previously imaged at the light microscope (LM) were mapped at low mag. After registration with the LM image, TEM tomography was performed on the areas of interest using the software package SerialEM^82^. Tomograms were reconstructed with IMOD^83^. Correlation between LM and EM images was done with the plugin ec-CLEM^84^ of the software platform Icy^85^, using features of the samples that could be identified in both imaging modalities.

### Image analysis

Image quantification analysis was performed using ImageJ or CellProfiler software. In particular, the presence of Nups AL foci was confirmed by eyes by two individuals.

For nuclear size or nuclear envelope intensity analysis of Nups, CellProfiler software was used. Briefly, the Prewitt Edge Finder method was used to enhance the edges in a DAPI image, and then a nucleus size threshold is set to automatically identify nuclei. This allowed identification and measurement of nuclei area, morphology factor and mean nuclear intensity for the desired channel. The software’s parameter measurements were exported to an Excel file and statistically analyzed.

For the size of Nups foci, ImageJ software was used. In brief, the Nups channel was thresholded in ImageJ and then the size of the Nups foci was calculated by analyzing particles (size 1-100). AL foci that were considered “large” had a size bigger than 0,3 μm^2^.

For co-localization analysis of RanBP2 and Nups in cytoplasm, CellProfiler software was used. The Prewitt Edge Finder method was used to enhance the edges in a DAPI image, and then nuclei, RanBP2 and different Nups were identified according to the corresponding channels. The cytoplasmic fraction of RanBP2 and Nups was identified using MaskObjects module. Co-localization analysis of RanBP2 and Nups in cytoplasm was conducted by MeasureObjectOverlap module. The software’s parameter measurements were exported to an Excel file and statistically analyzed. For nuclear envelope or cytoplasm intensity of Ran, ImageJ was used. The ROI of cell and nuclei were selected based on tubulin and DAPI channels, and the ROI of cytoplasm was selected by XOR of cell and nuclei. Finally, the mean gray value of the nuclei and cytoplasm was measured.

For AL-NPCs analysis in the cytoplasm, AL-NPCs were recognized by Nup133 signal surrounding Nup62 signals. The number of AL-NPCs was counted visually by two individuals.

For the measurement of NPCs density on the nuclear membrane, ImageJ software was used. Briefly, the images of the Nup62 channel were converted to 32 bits using ImageJ and then filtered using Gaussin blur (2). The nuclear region was manually selected and then find maxima (10) was performed to obtain the area of the selected region and the number of AL-NPCs.

For protein import and export assay analysis, ImageJ software was used. Briefly, the ROI within uniform distribution of NLS-mCherry-LEXY signal was selected in the nuclear region, and the measurement of the mean gray value of the same ROI in each frame (time sequentially) was performed by multi-measure option.

For calculation of the mean fluorescence intensity of Nups as a function of distance from the nucleus a custom Python script was used with help of the scikit-image module^86^. The nucleus and the nuclear envelope were identified using the RanBP2 channel following well-established procedures^87,88^. Similarly, the region corresponding to the ER was determined from the mScarlet-ER channel. Regions corresponding to RanBP2 foci were detected using the scikit-image method blob_log for blob detection. The analyzed region for Climp63 and mScarlet-ER corresponded to the ER region excluding the nucleus. The analyzed region for the RanBP2 foci corresponded to the blob regions that fall within the ER region excluding the nucleus. For each pixel in the analyzed regions, the minimum Euclidean distance to any nuclear envelope pixel was taken as the distance from the nucleus. Given the intensity-distance pairs for each pixel, the pixels were binned for the distance and the mean pixel intensity determined for each bin. The obtained histograms were finally normalized for purpose of comparison.

For Colocalization analysis, to assess pixel colocalization/correlation, we used correlation measurement within CellProfiler. Briefly, we used the “MaskImage” module to obtain Climp63, mScarlet-ER, and RanBP2 located in the cytoplasm. “Measure Colocalization” modules were used to study the colocalization and correlation between intensities in different images (different color channels) on a pixel-by-pixel basis across an entire image. The number of cells measured per condition was listed in the corresponding figure’s legend.

For nuclear height measurement, confocal microscopy was used to perform z-sampling of the DAPI channel of the sample, with a z-step of 0.5 μm. Finally, Fiji analysis was used to obtain an approximate nuclear height based on the Z-Extent.

### Protein sample preparation and Western blotting

After centrifugation at 250 g for 3 minutes at 4°C, cells were washed twice with cold PBS, and lysates were prepared using cell lysis buffer (50 mM Tris-HCl pH 7.5, 150 mM NaCl, 0.5 % sodium deoxycholate, 0.1% SDS, 0.5% sodium deoxycholate, 1 % Triton X-100, 1 mM EDTA, 1 mM EGTA, 2 mM Sodium pyrophosphate, 1 mM Na3VO4 and 1 mM NaF) supplemented with protease inhibitor cocktail (Roche) and incubated on ice for 30-60 min. After centrifugation at 16 000 g for 15 min at 4°C, the supernatant was transferred to the new clean Eppendorf tubes, and the total protein concentration was measured with a Bio-Rad Protein Assay kit (Bio-Rad). Separation of cytoplasmic and nuclear fractions was done using the NE-PER nuclear and cytoplasmic extraction reagent kit (Thermo Scientific™, 78833). Protein samples were boiled for 10 min at 95°C in 1X Laemmli buffer (LB) with β-Mercaptoethanol (BioRad, 1610747).

SDS-PAGE of proteins was done using pre-cast 4-12 % Bis-Tris gradient gels (Thermo Scientific, NW04120BOX) or pre-cast NuPAGE™ 3-8 % Tris-Acetate gradient Gels (Thermo Scientific, EA0378BOX). Proteins were transferred to a polyvinylidene difluoride (PVDF) membrane (Millipore, IPFL00010) using wet transfer modules (BIO-RAD Mini-PROTEAN® Tetra System). Membranes were blocked in 5 % bovine serum albumin (BSA, Millipore, 160069), or 5% non-fat milk powder mixed with 3 % BSA resuspended in TBS-T for 1h at room temperature, followed by incubation with primary antibodies diluted in TBS-T 5 % BSA/5 % milk overnight at 4°C or 1h at room temperature and secondary antibodies diluted in TBS-T 5 % BSA/5 % milk at room temperature. Membranes were imaged using SuperSignal West Pico (Pierce, Ref. 34580). Western blotting images were acquired by GE Healthcare_Amersham Imager 600 or Invitrogen iBright 1500. The grayscale value of protein bands was quantified using ImageJ software.

### Immunoprecipitation

Cell lysates were prepared for immunoprecipitations (IP) as described above. Lysates were adjusted to equal volume and concentration. For endogenous IP experiments, lysates were incubated with IgG or target-specific antibodies overnight at 4°C with rotation, and then incubated with protein G sepharose 4 Fast Flow beads (GE Healthcare Life Sciences) for 4h at 4°C with rotation. Before using, beads were blocked with 3 % BSA diluted in 1X cell lysis buffer and incubated for 2h at 4°C with rotation. The incubated IgG/ specific antibodies-samples-beads were washed with washing buffer (25 mM Tris-HCl pH 7.5, 300 mM NaCl, 0.5 % Triton X-100, 0.5 mM EDTA, 0.5 mM EGTA, 1 mM Sodium pyrophosphate, 0.5 mM Na3VO4 and 0.5 mM NaF) or TBS-T 4 to 6 times.

For GFP-IP/HA-IP experiments, GFP-Trap A agarose beads (Chromotek) or Pierce™ anti-HA Magnetic Beads (Thermo Scientific 88836) were used. Cells expressing GFP- or HA-tagged plasmids for at least 24h before lysis were used. Lysates were incubated with beads for 4h at 4°C with rotation, and then washed with washing buffer (25 mM Tris-HCl pH 7.5, 300 mM NaCl, 0.5 % Triton X-100, 0.5 mM EDTA, 0.5 mM EGTA, 1 mM Sodium pyrophosphate, 0.5 mM Na3VO4 and 0.5 mM NaF) or TBS-T 4 to 6 times.

### Gel fractionation

To calibrate the Superose 6 Increase 3.2/300 column, the gel filtration standard (Bio-Rad, Catalog # 1511901) was rehydrated and diluted to 1.5 mg/mL according to the instruction of the manufacturer. The column was equilibrated using buffer comprising 15 mM pH 7.5 Tris, 15 mM NaCl, 60 mM KCl, 340 mM sucrose, 0.15 mM spermine, 0.5 mM spermidine, 0.5 % Triton X-100, 1 mM NaF, 10 mM DTT. The standard sample (50 µL) was separated by a flow rate of 0.015 ml/min. Kav values for the standards were calculated. The calibration curve of Kav versus the logarithm of their molecular weights was determined as follows:

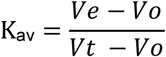

where Ve= elution volume of the protein Vo= column void volume = 0.8 mL

Vt = total bed volume = 2.4 mL

HeLa cell cytosolic and nuclear fractions were prepared as described above. For the gel filtration of the HeLa cell cytosol extraction sample, a 2.4-mL Superose 6 Increase 3.2/300 column (Cytiva) was equilibrated using buffer comprising 15 mM pH 7.5 Tris, 15 mM NaCl, 60 mM KCl, 340 mM sucrose, 0.15 mM spermine, 0.5 mM spermidine, 0.5 % Triton X-100, 1 mM NaF, 10 mM DTT. For the gel filtration of the HeLa cell nuclear fraction sample, a 2.4-ml Superose 6 Increase 3.2/300 column (Cytiva) was equilibrated using buffer comprising 50 mM Tris 7.5, 1 % NP-40, 150 mM NaCl, 1mM EDTA, 0.1 % SDS, 0.5 % sodium deoxycholate. The cytosolic and nuclear fractions (typically 1∼3 mg/ml) were clarified by centrifugation for 15 minutes at 12,000 X g before loading to the equilibrated gel filtration column. Samples were separated by a flow rate of 0.017 ml/min, and 50 µL fractions were collected for Western blot analysis (typically collects from 1.2mL to 2.4mL).

### LC-MS/MS analysis

MS grade Acetonitrile (ACN), MS grade H2O and MS grade formic acid (FA) were from ThermoFisher Scientific (Waltham, MA, USA). Sequencing-grade trypsin/Lys C mix was from Promega (Madison, WI, USA). Trifluoroacetic acid (TFA) and ammonium bicarbonate (NH4HCO3) were from Sigma-Aldrich (Saint-Louis, MO, USA).

For sample preparation prior to LC-MS/MS analysis beads from immunoprecipitation experiments were incubated overnight at 37°C with 20 μL of 50 mM NH4HCO3 buffer containing 1 µg of sequencing-grade trypsin/Lys C mix. The digested peptides were loaded and desalted on evotips provided by Evosep (Odense, Denmark) according to manufacturer’s procedure before LC-MS/MS analysis. For LC-MS/MS acquisition samples were analyzed on a timsTOF Pro 2 mass spectrometer (Bruker Daltonics, Bremen, Germany) coupled to an Evosep one system (Evosep, Odense, Denmark) operating with the 30SPD method developed by the manufacturer. Briefly, the method is based on a 44-min gradient and a total cycle time of 48 min with a C18 analytical column (0.15 x 150 mm, 1.9µm beads, ref EV-1106) equilibrated at 40°C and operated at a flow rate of 500 nL/min. H2O/0.1 % FA was used as solvent A and ACN/ 0.1 % FA as solvent B. The timsTOF Pro 2 was operated with a DIA-PASEF method comprising 12 pydiAID frames with 3 mass windows per frame resulting in a cycle time of 0.975 seconds as described in Bruker application note LCMS 218.

For data analysis MS raw files were processed using Spectronaut 18 (Biognosys, Switzerland). Data were searched against the SwissProt Homo Sapiens database (downloaded 07 2023, 20423 entries). Specific tryptic cleavages were selected and a maximum of 2 missed cleavages were allowed. The following post-translational modifications were considered for identification: Acetyl (Protein N-term) and Oxidation (M) as variables. Identifications were filtered based on a 1 % precursor and protein Qvalue cutoff threshold. The protein LFQ method was set to automatic and the quantity was set at the MS2 level with a cross-run normalization applied. Multivariate statistics on protein measurements were performed using Qlucore Omics Explorer 3.9 (Qlucore AB, Lund, SWEDEN). A positive threshold value of 1 was set to allow a log2 transformation of abundance data for normalization i.e. all abundance data values below the threshold are replaced by 1 before transformation. The transformed data were finally used for statistical analysis i.e. the evaluation of differentially present proteins between two groups using a bilateral Student’s t-test. A p-value better than 0.05 was used to filter differential candidates.

### Experimental design, data acquisition and statistical analysis

All experiments were done in a strictly double-blinded manner. At least three independent biological replicates were performed for each experiment (unless otherwise indicated) and image quantifications were carried out in a blinded manner. Curves and graphs were made using GraphPad Prism and Adobe Illustrator software. Data was analyzed using one-sample two-tailed T-test or two sample unpaired two-tailed T-test (two-group comparison or folds increase relative to the control, respectively) or one-way ANOVA. A p-value less than 0.05 (typically ≤ 0.05) was considered statistically significant and stars were assigned as follows: *P < 0.05, **P < 0.01, ***P < 0.001, ****P < 0.0001.

## Legends of Supplementary Table 1 and Supplementary Videos 1-10

**SUPPLEMENTARY Table 1. List of reagents used for cloning, siRNA, and CRISPR/Cas9 methods**

Table summarizes cloning primers, siRNAs and guide RNAs used in this study.

**Supplementary Video 1. AL foci cluster following MT depolymerization**

2xZFN-mEGFP-Nup107 HeLa cells treated with nocodazole were analyzed by live video spinning disk confocal microscopy. Z-stacks (10 mm range, 0.5 mm step) were acquired every 10 minutes and maximum intensity projection images are shown at speed 5 frames per second. A representative cell is depicted and selected representative frames of the movie are shown in the Fig. S5C.

**Supplementary Video 2. AL foci merge with the NE throughout interphase**

2xZFN-mEGFP-Nup107 HeLa cells were synchronized by double thymidine block, released and analyzed by live video spinning disc confocal microscopy. Timepoint 0 indicates the mitotic exit and entry into a new cell cycle. Z-stacks (10 mm range, 0.5 mm step) were acquired every 5 minutes and maximum intensity projection images are shown at speed 5 frames per second. **Supplementary Video 3. AL foci are highly dynamic under physiological conditions** 2xZFN-mEGFP-Nup107 HeLa cells were analyzed by live video spinning disc confocal microscopy. Z-stacks (10 mm range, 1 mm step) were acquired every 1 second and maximum intensity projection images are shown at speed 5 frames per second. A representative cell is depicted and selected representative frames of the movie are shown in Fig. 2E. **Supplementary Video 4. Smaller AL foci fuse more quickly with the NE than larger AL** 2xZFN-mEGFP-Nup107 HeLa cells were analyzed by live video spinning disc confocal microscopy. Z-stacks (10 mm range, 1 mm step) were acquired every 3 seconds and maximum intensity projection images are shown at speed 5 frames per second. Three representative cells are depicted and selected representative frames of the movie are shown in Fig. 2E.

**Supplementary Video 5. AL occasionally undergo multiple cycles of detachment and re-attachment before ultimately fusing with the NE**

2xZFN-mEGFP-Nup107 HeLa cells were analyzed by live video spinning disc confocal microscopy. Z-stacks (10 mm range, 1 mm step) were acquired every 3 seconds and maximum intensity projection images are shown at speed 5 frames per second. A representative cell is depicted and selected representative frames of the movie are shown in Fig. 2E.

**Supplementary Video 6. NE-NPCs do not diffuse from the nuclear envelope into the cytoplasm.**

mEos2-Nup133 HeLa cells were analyzed by live video spinning disc confocal microscopy. Nup133 on the nuclear envelope was converted from green fluorescence (left channel) to red fluorescence (right channel) by blue light for 30 seconds. Z-stacks (10 mm range, 1 mm step) were used. A 30-second acquisition pre-sequence with acquisition every 10 seconds in the green and red channels was performed. Two post sequences are carried out after acquisition, the first for 30 seconds with an interval every 10 seconds in order to quickly observe the effect of photoconversion and the second sequence for 12 hours every 15 minutes to see the effect over a longer period. The maximum intensity projection images are shown at 5 frames per second.

**Supplementary Video 7. AL-NPCs accumulate at the nuclear envelope over time.** mEos2-Nup133 HeLa cells were analyzed by live video spinning disc confocal microscopy. Nup133 in the cytoplasm was converted from green fluorescence (left channel) to red fluorescence (right channel) by blue light for 30 seconds. Z-stacks (10 mm range, 1 mm step) were used. A 30-second acquisition pre-sequence with acquisition every 10 seconds in the green and red channels was performed. Two post sequences are carried out after acquisition, the first for 30 seconds with an interval every 10 seconds in order to quickly observe the effect of photoconversion and the second sequence for 12 hours every 15 minutes to see the effect over a longer period. The maximum intensity projection images are shown at speed 5 frames per second.

**Supplementary Video 8. AL foci move along ER to merge with the NE**

2xZFN-mEGFP-Nup107 HeLa cells expressing ER marker plasmid mScarlet-ER were analyzed by live video spinning disk confocal microscopy. Timepoint 0 indicates the start of the imaging. Frames were acquired every 20s (without Z-stacks) and images are shown at 5 frames per second. A representative cell is depicted in Fig. 3D and a zoomed region of selected representative frames of the video is shown.

**Supplementary Video 9. Microtubule depolymerization results in the inhibition of AL foci integration into the NE and their clustering**

2xZFN-mEGFP-Nup107 HeLa cells expressing ER marker plasmid mScarlet-ER were analyzed by live video spinning disk confocal microscopy. Timepoint 0 indicates the start of the imaging. Frames were acquired every 120s (without Z-stacks) and images are shown at 5 frames per second. A representative cell is depicted in Fig. 3D and a zoomed region of selected representative frames of the video is shown.

**Supplementary Video 10. AL are present on microtubules**

2xZFN-mEGFP-Nup107 HeLa cells were analyzed by live video spinning disc confocal microscopy. SiR-tubulin was used to label microtubules. Z-stacks were acquired every 2 seconds and images are shown at 5 frames per second.

---

**Extended Data Fig.1.**
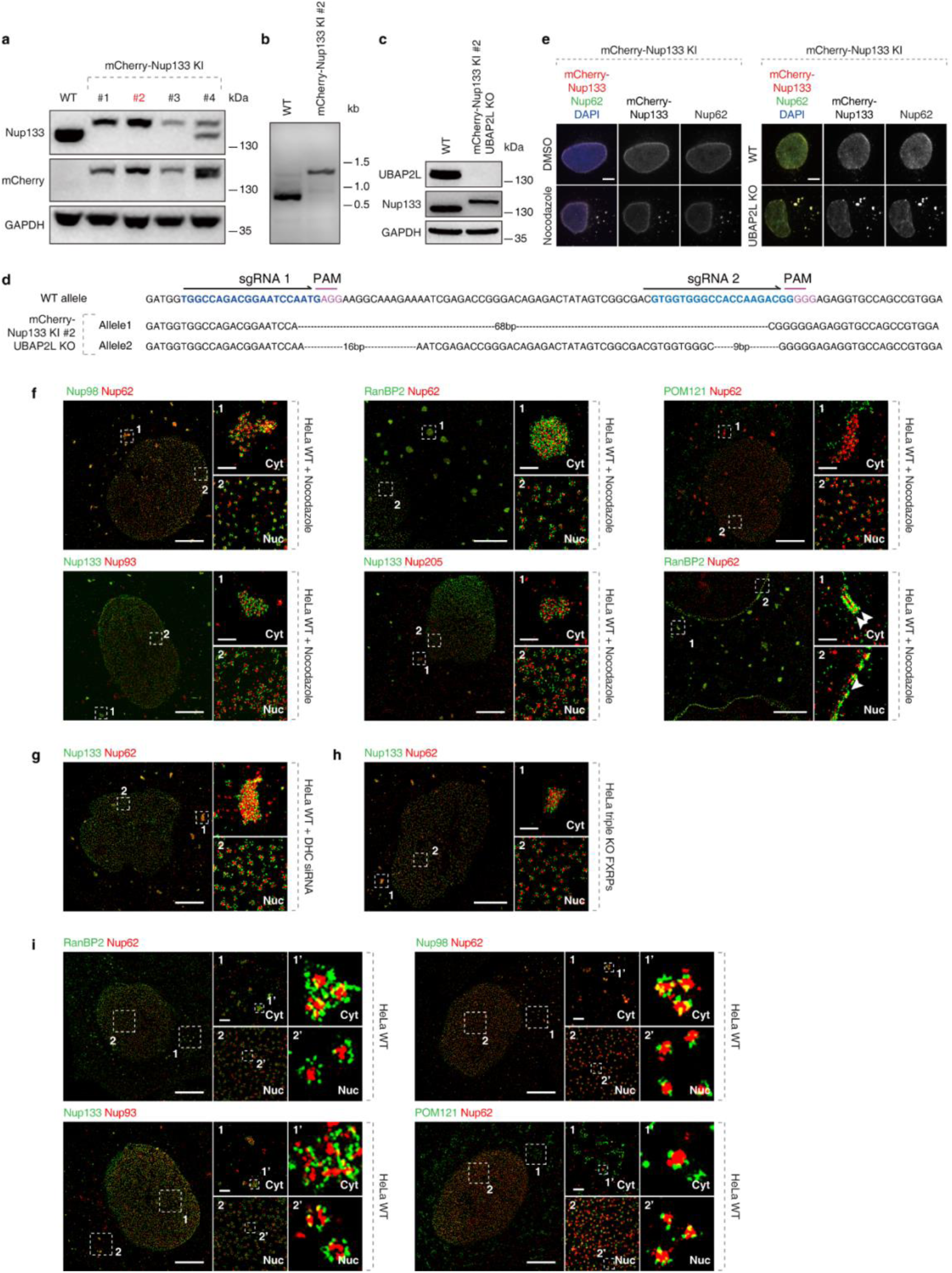
**Single molecule localization microscopy identifies AL in WT and in challenged HeLa cells.** a, b, Validation of CRISPR/Cas9-mediated mCherry-Nup133 KI Hela cell clones by Western blot (a) and PCR (b). c,d, Validation of CRISPR/Cas9-mediated UBAP2L KO mCherry-Nup133 KI HeLa cell clones by Western blot (c) and Sanger sequencing (d). e, Representative images of mCherry-Nup133 KI HeLa cells treated with DMSO or Nocodazole (10 μM) and representative images of UBAP2L KO mCherry-Nup133 KI HeLa. Cells were co-labelled with DAPI (blue), and anti-Nup62 (green) antibody. Scale bars, 5 μm. f, Representative splitSMLM images depicting NPCs on the nuclear (Nuc) surface and AL-NPCs in the cytoplasm (Cyt) in nocodazole-treated (10 μM, 90 min) HeLa cells. Analyzed Nups from different NPC subcomplexes are indicated above each panel. The magnified framed regions are shown in the corresponding numbered panels. Scale bars, 3 μm (entire nuclei), 0.3 μm (zoomed regions). g, h, Representative splitSMLM images depicting NPCs on the nuclear (Nuc) surface and AL-NPCs in the cytoplasm (Cyt) in HeLa cell treated with the dynein heavy chain siRNAs (48h) and triple knock-out of FXRPs HeLa cells. Nup133 signal labels the cytoplasmic and nuclear rings of the NPC and the localization of the central channel is visualized by Nup62. The magnified framed regions are shown in the corresponding numbered panels. Scale bars, 3 μm (entire nuclei), 0.3 μm (zoomed regions). i, Representative splitSMLM images depicting NPCs on the nuclear (Nuc) surface and AL-NPCs in the cytoplasm (Cyt) in WT HeLa cells. Analyzed Nups from different NPC subcomplexes are indicated above each panel. The magnified framed regions are shown in the corresponding numbered panels. Scale bars, 3 μm (entire nuclei), 0.3 μm (zoomed regions).

**Extended Data Fig.2.**
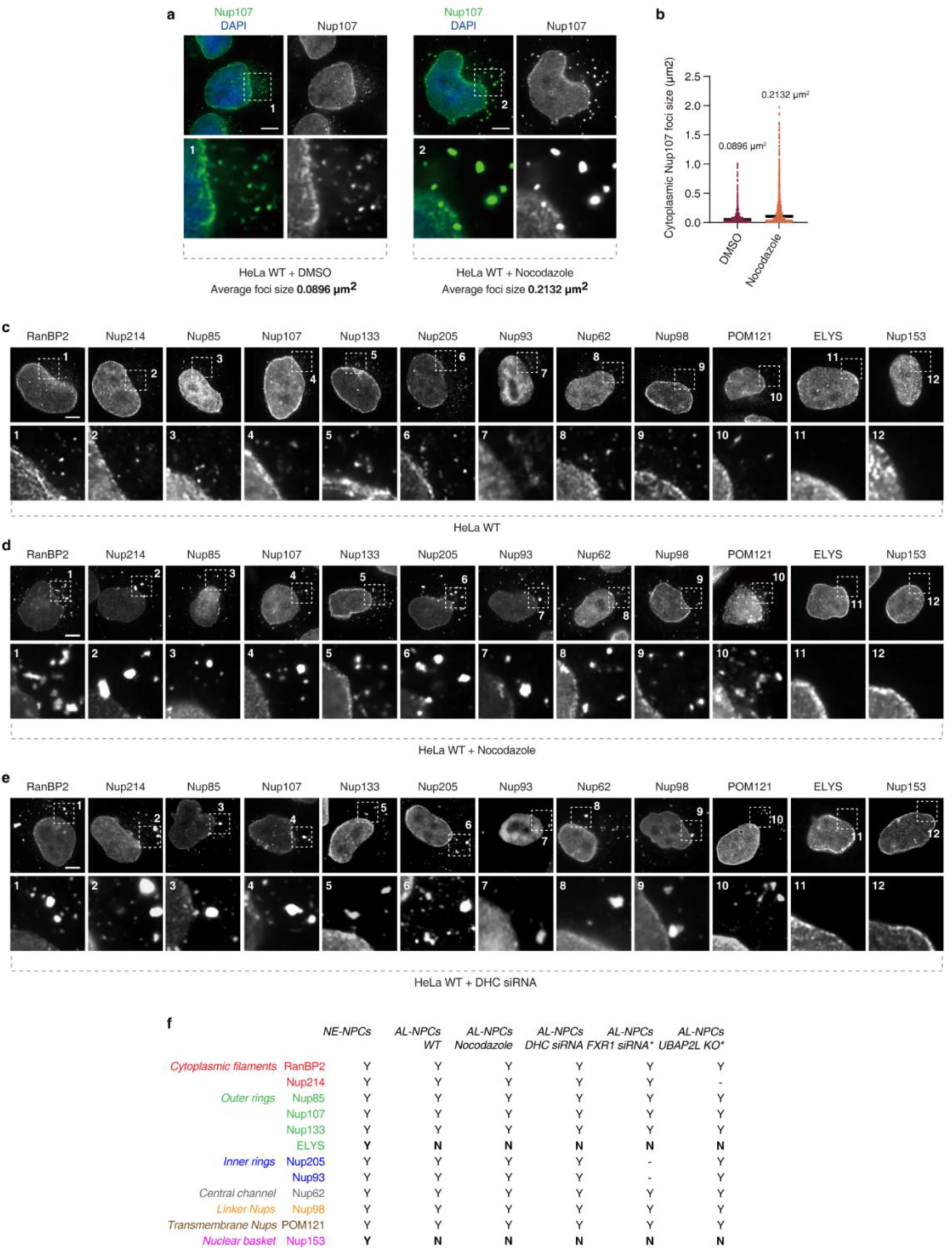
**Systematic analysis of the size and the composition of AL foci.** a, b, Representative images of 2xZFN-mEGFP-Nup107 HeLa cells treated with nocodazole (10 μM) or DMSO (90 min) (a). Cells were co-labelled with DAPI (blue). The magnified framed regions are shown in the corresponding numbered panels. The size of Nup107-positive AL foci in the cytoplasm were quantified in (b), and at least 2000 foci per condition were analyzed (N=3). Scale bars, 5 μm. c, Representative images of WT HeLa cells labelled with Nups from different NPC subcomplexes. Analyzed Nups are indicated above each panel. The magnified framed regions are shown in the corresponding numbered panels. Scale bars, 5 μm. d, Representative images of nocodazole-treated (10 μM, 90 min) HeLa cells labelled with Nups from different NPC subcomplexes. Analyzed Nups are indicated above each panel. The magnified framed regions are shown in the corresponding numbered panels. Scale bars, 5 μm. e, Representative images of dynein heavy chain siRNAs-treated (48h) HeLa cells labelled with Nups from different NPC subcomplexes. Analyzed Nups are indicated above each panel. The magnified framed regions are shown in the corresponding numbered panels. Scale bars, 5 μm. f, Summary of composition of NE-NPCs and AL-NPCs under different conditions. Different colors represent specific NPC subcomplexes. “Y” indicates presence and “N” indicates absence of indicated Nups. “-” represents no available data. “*” indicates published results^25,27,28^.

**Extended Data Fig.3.**
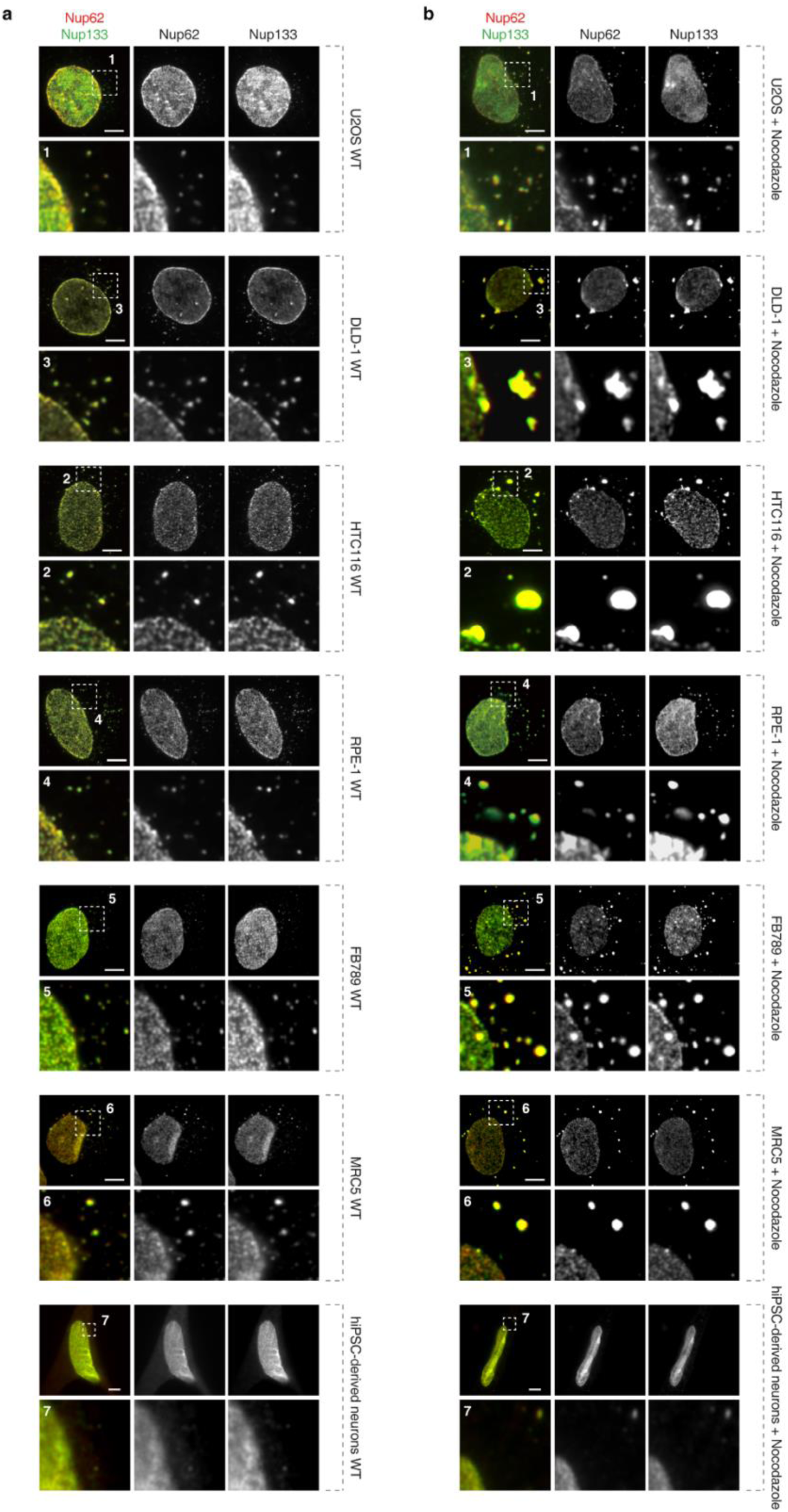
**AL exist in a variety of somatic mammalian cells.** a, Representative images of WT U2OS, DLD-1, HTC116, RPE-1, FB789, MRC5 and hiPSC-derived neuron cells showing AL foci. Cells were co-labelled with anti-Nup62 (red) and anti-Nup133 (green) antibodies to visualize AL-NPCs. Scale bars, 5 μm. b, Representative images of U2OS, DLD-1, HTC116, RPE-1, FB789, MRC5 and hiPSC-derived neuron cells treated with nocodazole (10 μM, 90 min). Cells were co-labelled with anti-Nup62 (red) and anti-Nup133 (green) antibodies to visualize AL-NPCs. Scale bars, 5 μm.

**Extended Data Fig.4.**
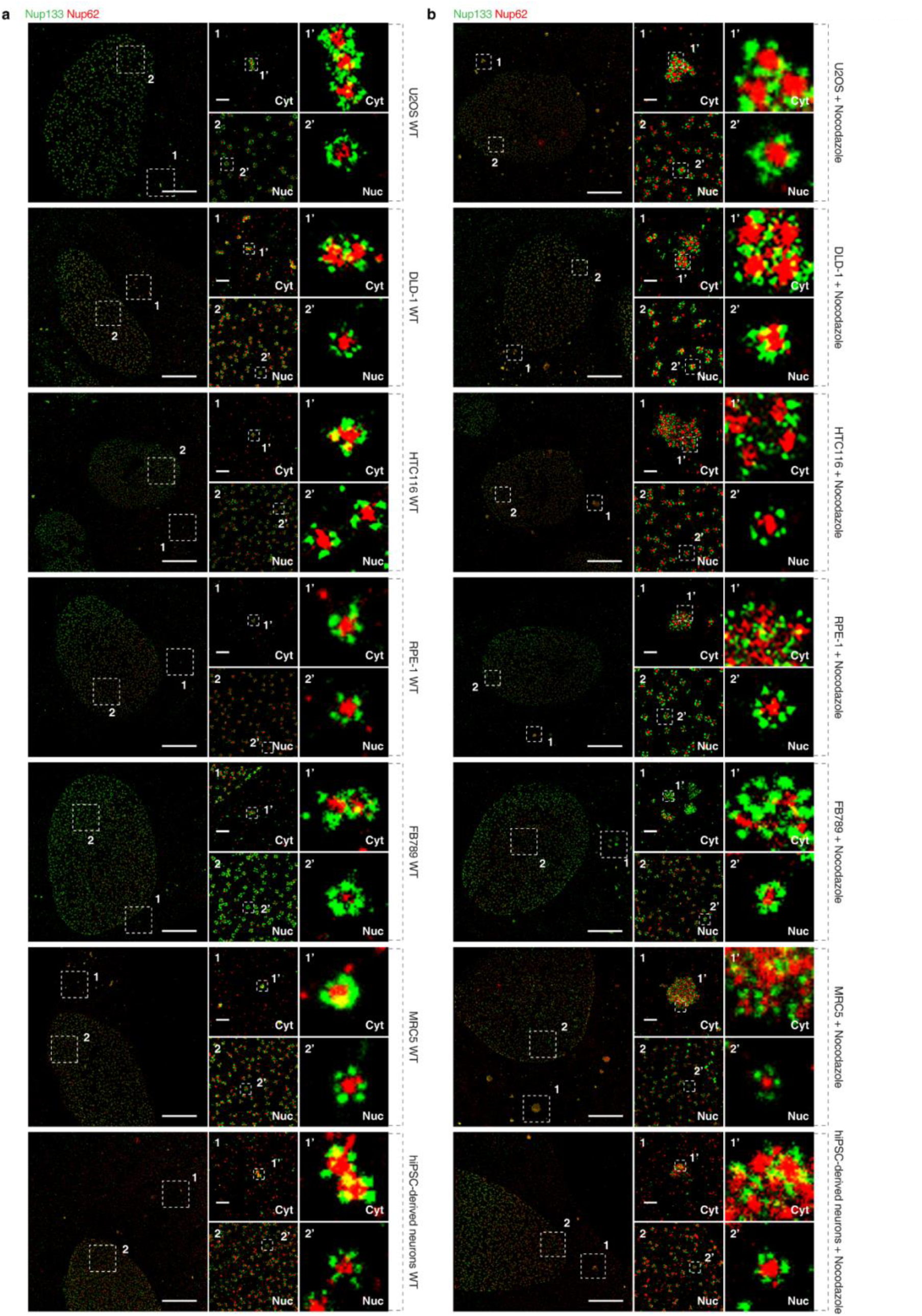
**AL exist in a variety of somatic mammalian cells.** a, Representative splitSMLM images depicting NPCs on the nuclear (Nuc) surface and AL-NPCs in the cytoplasm (Cyt) in WT U2OS, DLD-1, HTC116, RPE-1, FB789, MRC5 and hiPSC-derived neuron cells. Analyzed Nups from different NPC subcomplexes are indicated above each panel. The magnified framed regions are shown in the corresponding numbered panels. Scale bars, 3 μm (entire nuclei), 0.3 μm (zoomed regions). b, Representative splitSMLM images depicting NPCs on the nuclear (Nuc) surface and AL-NPCs in the cytoplasm (Cyt) in nocodazole-treated (10 μM, 90 min) U2OS, DLD-1, HTC116, RPE-1, FB789, MRC5 and hiPSC-derived neuron cells. Analyzed Nups from different NPC subcomplexes are indicated above each panel. The magnified framed regions are shown in the corresponding numbered panels. Scale bars, 3 μm (entire nuclei), 0.3 μm (zoomed regions).

**Extended Data Fig.5.**
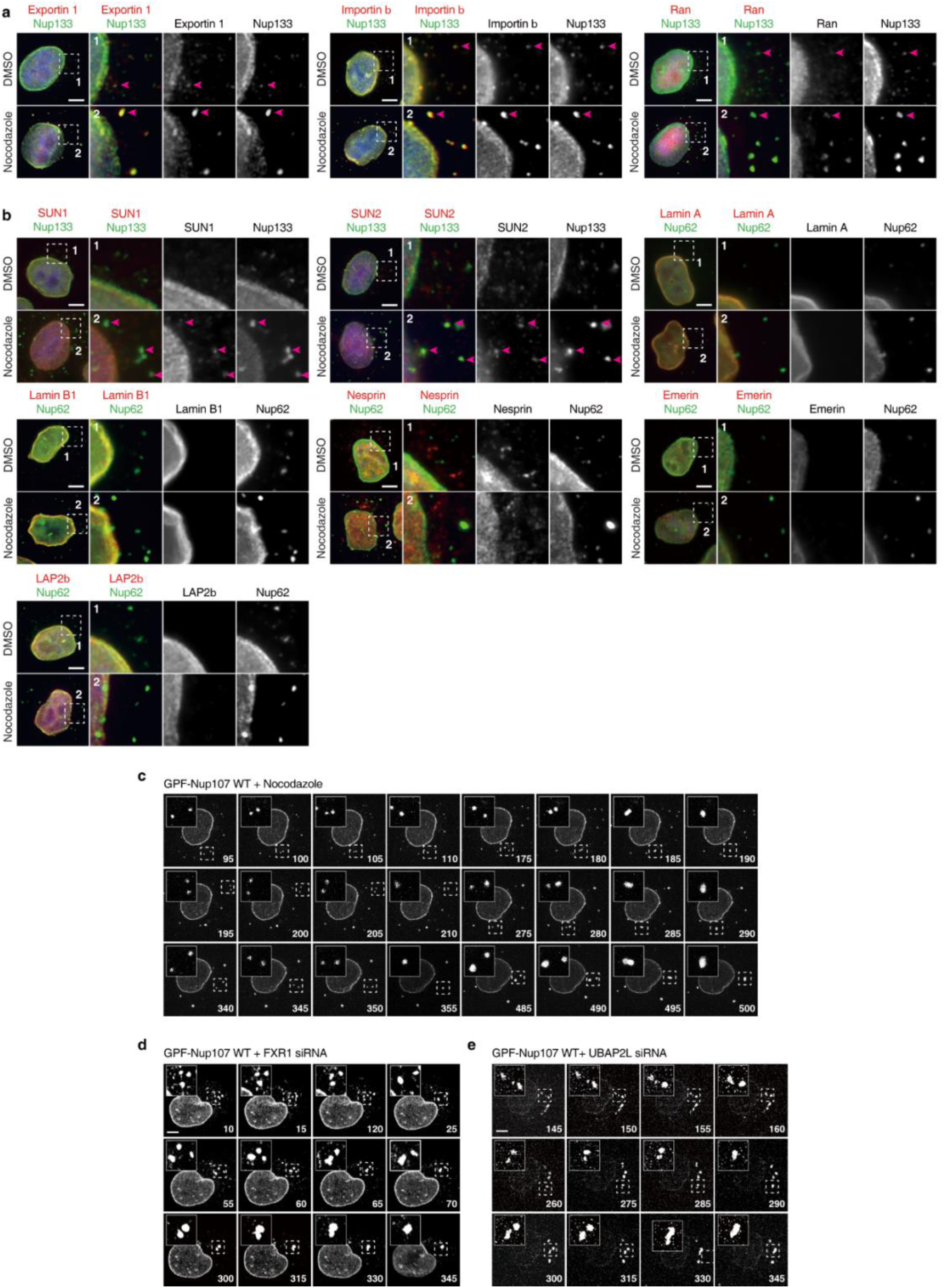
Systematic analysis of AL composition and clustering of AL-NPCs. a, Representative images showing co-localization of nucleocytoplasmic transport factors (NTF) with AL-NPC foci in HeLa cells treated with nocodazole (10 μM) or DMSO (90 min). Cells were co-labelled with anti-NTF (Exportin-1, Importin β, and Ran) (red) and anti-Nup133 (green) antibodies. The magnified framed regions are shown in the corresponding numbered panels. Magenta arrows point to co-localized AL and NTF foci signals. Scale bars, 5 μm. b, Representative images showing co-localization of NE-related proteins with AL-NPC foci in HeLa cells treated with nocodazole (10 μM) or DMSO (90 min). Cells were co-labelled with anti-NE-related proteins (SUN1, SUN2, Lamin A, Lamin B1, Nesprin, Emerin, and Lap2) (red) and anti-Nup62 (green) antibodies. The magnified framed regions are shown in the corresponding numbered panels. Magenta arrows point to co-localized AL and NE-factors foci. Scale bars, 5 μm. c, 2xZFN-mEGFP-Nup107 HeLa cells treated with nocodazole were analyzed by live video spinning disk confocal microscopy. Selected representative frames of the Videos are depicted, and time is shown in minutes after nocodazole addition at the timepoint 0. The magnified framed regions are shown in the upper left corner. Scale bars, 5 μm. d, 2xZFN-mEGFP-Nup107 HeLa cells treated with FXR1 siRNA were analyzed by live video spinning disk confocal microscopy. The selected representative frames of the Videos are depicted, and time is shown in minutes. The timepoint 0 of the Video corresponds to a 12h release time from double thymidine block. The magnified framed regions are shown in the upper left corner. Scale bars, 5 μm. e, 2xZFN-mEGFP-Nup107 HeLa cells treated UBAP2L siRNA were analyzed by live video spinning disk confocal microscopy. The selected representative frames of the Videos are depicted, and time is shown in minutes. The timepoint 0 of the Video corresponds to a 12h release time from double thymidine block. The magnified framed regions are shown in the upper left corner. Scale bars, 5 μm.

**Extended Data Fig.6.**
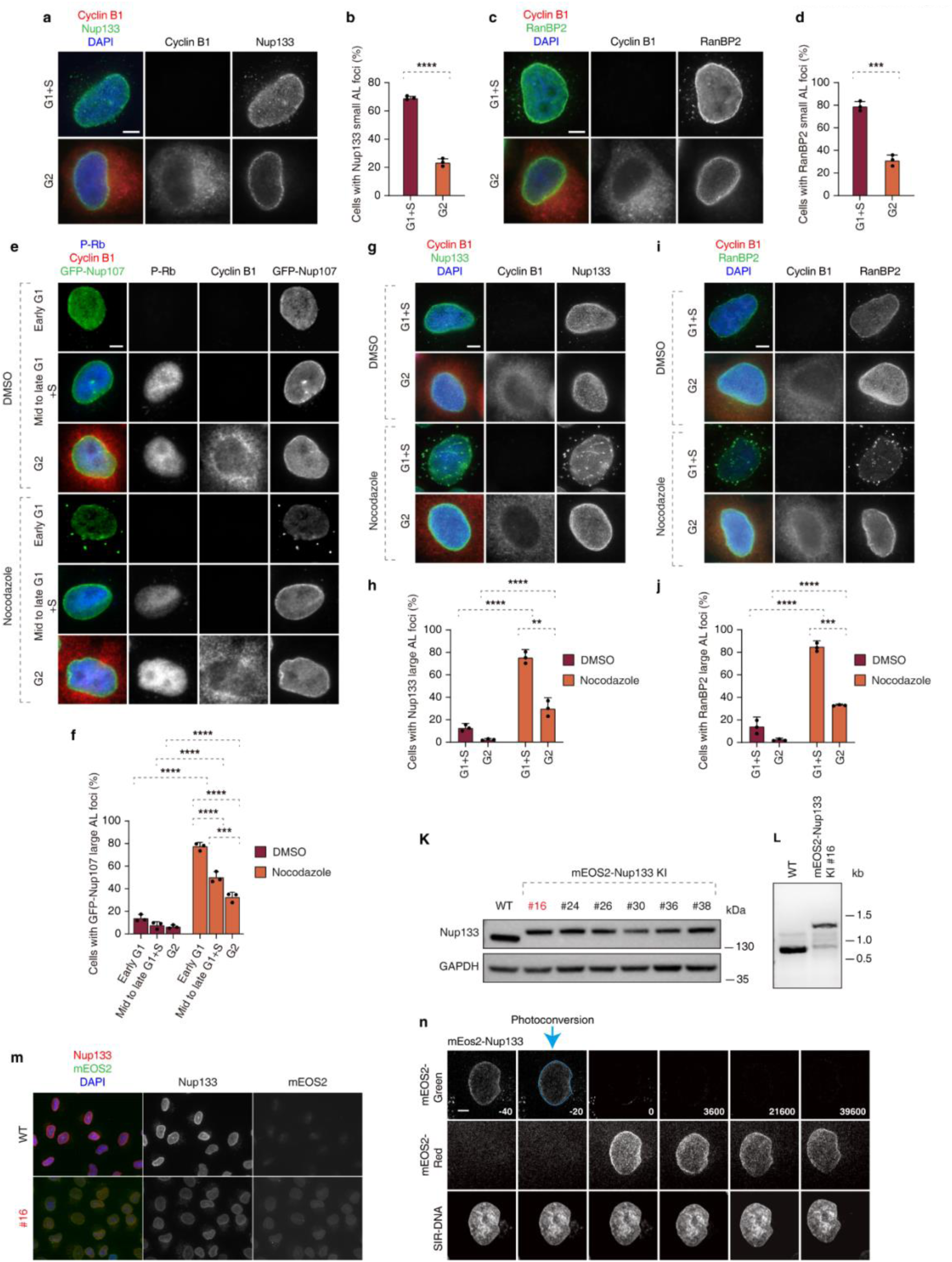
**AL foci are predominantly present in the G1 and S cell cycle phases.** a, b, Representative images of asynchronously proliferating HeLa cells co-labelled with anti-Nup133 (green) and anti-cyclin B1 (red) (a) antibodies. The percentage of cells with Nup133-positive small AL foci in cyclin B1 negative cells (G1+S), and cyclin B1 positive cells (G2) was quantified in (b), and at least 200 cells per condition were analyzed (unpaired two-tailed t test, mean ± SD, ****P < 0.0001; N = 3). Scale bars, 5 μm. c, d, Representative images of asynchronously proliferating HeLa cells co-labelled with anti-RanBP2 (green) and anti-cyclin B1 (red) (c) antibodies. The percentage of cells with RanBP2-positive small AL foci in cyclin B1 negative cells (G1+S), and cyclin B1 positive cells (G2) was quantified in (d), and at least 200 cells per condition were analyzed (unpaired two-tailed t test, mean ± SD, ***P < 0.001; N = 3). Scale bars, 5 μm. e, f, Representative images of asynchronously proliferating 2xZFN-mEGFP-Nup107 HeLa cells treated with nocodazole (10 μM) or DMSO (90 min) and co-labelled with anti-p-Rb (blue) and anti-cyclin B1 (red) (e) antibodies. The percentage of cells with large AL foci in p-Rb and cyclin B1 negative cells (early G1), p-Rb positive and cyclin B1 negative cells (mid to late G1+S), and p-Rb and cyclin B1 positive cells (G2) was quantified in (f), and at least 200 cells per condition were analyzed (one-Way ANOVA test, mean ± SD, ***P < 0.001; ****P < 0.0001; N = 3). Scale bars, 5 μm. g, h, Representative images of asynchronously proliferating HeLa cells treated with nocodazole (10 μM) or DMSO (90 min) and co-labelled with anti-Nup133 (green) and anti-cyclin B1 (red) (g) antibodies. The percentage of cells with large AL foci in cyclin B1 negative cells (G1+S), and cyclin B1 positive cells (G2) was quantified in (h), and at least 200 cells per condition were analyzed (one-Way ANOVA test, mean ± SD, **P < 0.01; ****P < 0.0001; N = 3). Scale bars, 5 μm. i, j, Representative images of asynchronously proliferating HeLa cells treated with nocodazole (10 μM) or DMSO (90 min) and co-labelled with anti-RanBP2 (green) and anti-cyclin B1 (red) (i) antibodies. The percentage of cells with large AL foci in cyclin B1 negative cells (G1+S), and cyclin B1 positive cells (G2) was quantified in (j), and at least 200 cells per condition were analyzed (one-Way ANOVA test, mean ± SD, ***P < 0.001; ****P < 0.0001; N = 3). Scale bars, 5 μm. k, l, Validation of CRISPR/Cas9-mediated mEos2-Nup133 KI HeLa cell clones by Western blot (k) and PCR (l). m, Representative images of mEos2-Nup133 KI HeLa cells. Cells were co-labelled with DAPI (blue), and anti-Nup133 (red) antibody. Scale bars, 5 μm. n, The photoconversion of NE-Nup133 in mEOS2-Nup133 HeLa cells were analyzed by live video spinning disk confocal microscopy. The selected representative frames of the Videos are depicted, and time is shown in seconds. The magnified framed regions of mEOS2-Red signal are shown in the lower panels. The blue dashed box indicates the photoconversion region, where mEOS2-Nup 133 is converted from a green to a red fluorescent state. Scale bars, 5 μm.

**Extended Data Fig.7.**
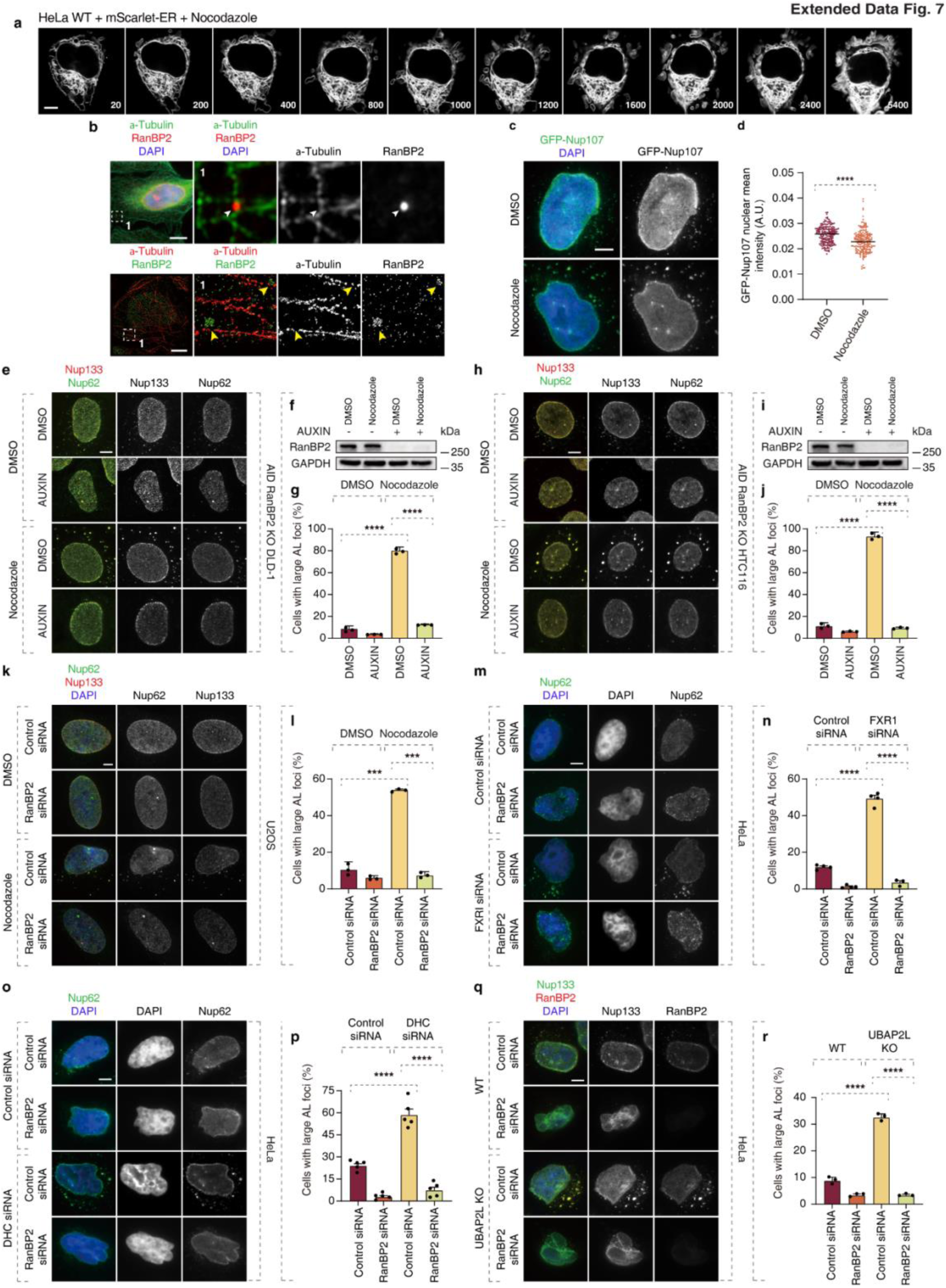
**RanBP2 is required for the accumulation and clustering of large AL in challenged somatic cells.** a, HeLa cells expressing ER marker plasmid mScarlet-ER were treated with nocodazole (10 μM) and were analyzed by live video spinning disk confocal microscopy. The selected representative frames of the Videos are depicted, and time is shown in second. The timepoint 0 corresponds to nocodazole addition. The magnified framed regions are shown in the upper left corner. Scale bars, 5 μm. b, Representative immunofluorescence (IF) images and splitSMLM analysis depicting co-localization of AL-NPCs and microtubules in the cytoplasm in WT HeLa cells. For the IF at the top, cells were co-labelled with anti-alpha tubulin (green), anti-RanBP2 (red) antibodies and DAPI. For the splitSMLM at the bottom, cells were co-labelled with anti-alpha tubulin (red), and anti-RanBP2 (green) antibodies. The magnified framed regions are shown in the corresponding numbered panels. Arrowheads point to AL-NPCs localized near microtubules. Scale bars, IF, 5 μm; SMLM, 3 μm. c, d, Representative images of 2xZFN-mEGFP-Nup107 HeLa cells treated with Nocodazole (10 μM) or DMSO (90 min) (c). Cells were co-labelled with DAPI (blue). The intensity of EGFP-Nup107 in the NE was quantified in (d), and at least 200 cell per condition were analyzed (unpaired two-tailed t test, mean ± SD, ****P < 0.0001; N = 3). Scale bars, 5 μm. e, f, Representative images of auxin-inducible degron (AID)-RanBP2 KO DLD-1 cells treated with the auxin or DMSO (4h) and subsequently with nocodazole (10 μM) or DMSO (90 min). Cells were co-labelled with anti-Nup62 (red) and anti-Nup133 (green) (e) antibodies. (B) The Western blot analysis of RanBP2 under different conditions shown in (f). The number of cells with large AL foci were quantified in (G), and at least 1000 cells per condition were analyzed (one-Way ANOVA test, mean ± SD, ****P < 0.0001; N = 3). Scale bars, 5 μm. h-j, Representative images of AID-RanBP2 KO HTC116 cells treated with auxin or DMSO (4h) and subsequently with nocodazole (10 μM) or DMSO (90 min). Cells were co-labelled with anti-Nup62 (red) and anti-Nup133 (green) (h) antibodies. (i) The Western blot analysis of RanBP2 under different conditions shown in (j). The number of cells with large AL foci were quantified in (K), and at least 1000 cells per condition were analyzed (one-Way ANOVA test, mean ± SD, ****P < 0.0001; N = 3). Scale bars, 5 μm. k, l, Representative images of U2OS cells treated with the indicated siRNAs (48h) and subsequently with nocodazole (10 μM) or DMSO (90 min). Cells were co-labelled with anti-Nup62 (green) and anti-Nup133 (red) (k) antibodies. The number of cells with large AL foci was quantified in (l), and at least 1000 cells per condition were analyzed ((one-Way ANOVA test, mean ± SD, ***P < 0.001; N = 3). Scale bars, 5 μm. m, n, Representative images of HeLa cells treated with the indicated siRNAs (48h). Cells were co-labelled with anti-Nup62 (green) antibodies and DAPI (blue) (m). The number of cells with large AL foci were quantified in (n), and at least 1000 cells per condition were analyzed (one-Way ANOVA test, mean ± SD, ****P < 0.0001; N = 3). Scale bars, 5 μm. o, p, Representative images of HeLa cells treated with the indicated siRNAs (48h). Cells were co-labelled with anti-Nup62 (green) antibodies and DAPI (blue) (o). The number of cells with large AL foci were quantified in (p), and at least 1000 cells per condition were analyzed (mean ± SD, ****P < 0.0001; N = 3). Scale bars, 5 μm. q, r, Representative images of UBAP2L KO and WT HeLa cells treated with the indicated siRNAs (48h). Cells were co-labelled with anti-Nup133 (green) and anti-RanBP2 (red) (q) antibodies. The number of cells with large AL foci were quantified in (r), and at least 1000 cells per condition were analyzed (one-Way ANOVA test, mean ± SD, ****P < 0.0001; N = 3). Scale bars, 5 μm.

**Extended Data Fig. 8.**
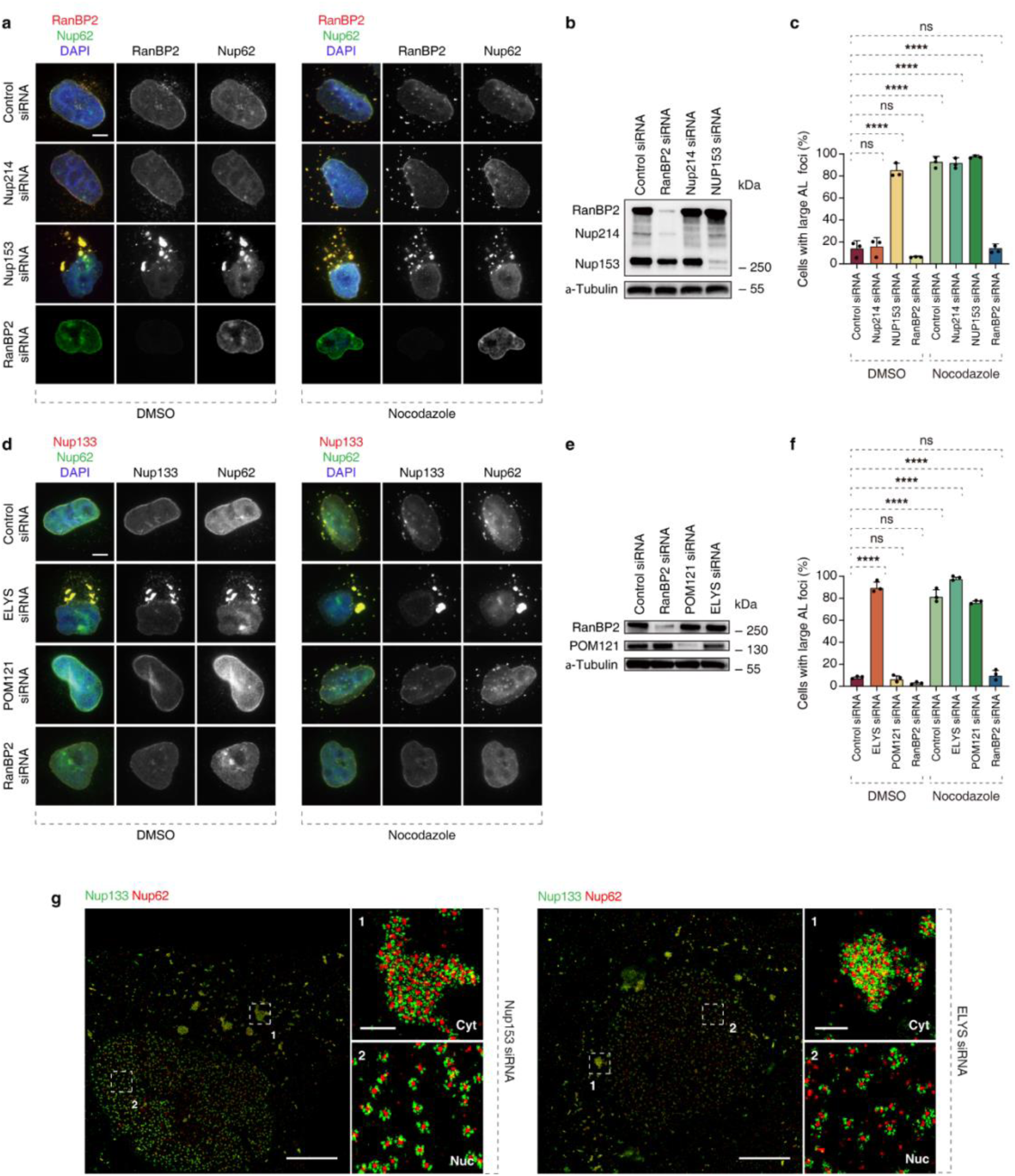
**RanBP2, but not Nup153, Nup214, POM121 and ELYS, is required for the formation of large AL in challenged somatic cells.** a-c, Representative images of HeLa cells treated with the indicated siRNAs (48h) and subsequently with nocodazole (10 μM) or DMSO (90 min). Cells were co-labelled with anti-Nup62 (green) and anti-RanBP2 (red) (a) antibodies. (b) Shows the Western blot analysis of RanBP2, Nup214 and Nup153 under different conditions shown in (a). The number of cells with large AL foci was quantified in (c), and at least 1000 cells per condition were analyzed (one-Way ANOVA test, mean ± SD, ns: not significant, ****P < 0.0001; N = 3). Scale bars, 5 μm. d-f, Representative images of HeLa cells treated with the indicated siRNAs (48h) and subsequently with nocodazole (10 μM) or DMSO (90 min). Cells were co-labelled with anti-Nup62 (green) and anti-Nup133 (red) (d) antibodies. (e) Shows the Western blot analysis of RanBP2, and POM121 under different conditions shown in (d). The number of cells with large AL foci was quantified in (f), and at least 1000 cells per condition were analyzed (one-Way ANOVA test, mean ± SD, ns: not significant, ****P < 0.0001; N = 3). Scale bars, 5 μm. g, Representative splitSMLM images depicting NPCs on the nuclear (Nuc) surface and AL-NPCs in the cytoplasm (Cyt) in Nup153 siRNA or ELYS siRNA-treated (48h) HeLa cells. Nup133 signal labels the cytoplasmic and nuclear rings of the NPC, the localization of the central channel is visualized by Nup62. The magnified framed regions are shown in the corresponding numbered panels. Scale bars, 3 μm, 0.3 μm (zoomed-in).

**Extended Data Fig. 9.**
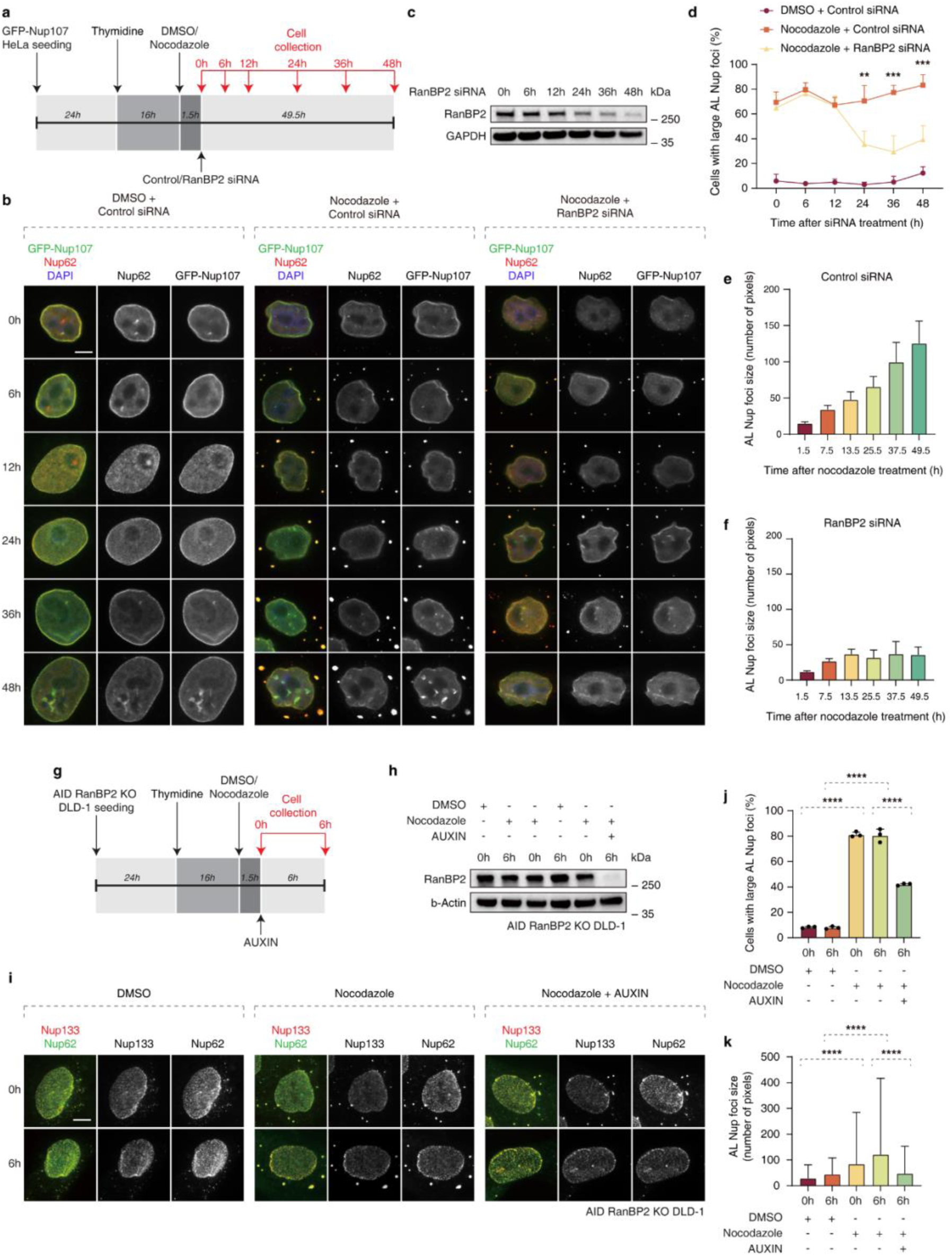
**RanBP2 is required for the maintenance of large AL.** a-f, Scheme of the experimental setup (a). HeLa cells were synchronized and arrested in S phase by thymidine treatment (16h), treated with nocodazole (10 μM) and subsequently with the indicated siRNAs, and fixed at indicated timepoints. Representative images of cells are shown in (b). (c) Shows the Western blot analysis of RanBP2 at indicated timepoints. The number of cells with large AL foci was quantified in (d) and the size of AL foci was quantified in (e, f). At least 200 cells and 1500 foci per condition were analyzed (one-Way ANOVA test, mean ± SD, ****P < 0.0001; N = 3). Scale bars, 5 μm. g-k, Scheme of the experimental setup (g). AID-RanBP2 KO DLD-1 cells were synchronized and arrested at S phase by thymidine treatment (16h), treated with nocodazole (10 μM) and with auxin. Representative images of cells are shown in (i). (h) The Western blot analysis of RanBP2 under different conditions. The number of cells with large AL foci was quantified in (j) and the size of AL foci was quantified in (k). At least 200 cells and 1500 foci per condition were analyzed (one-Way ANOVA test, mean ± SD, ****P < 0.0001; N = 3). Scale bars, 5 μm

**Extended Data Fig. 10.**
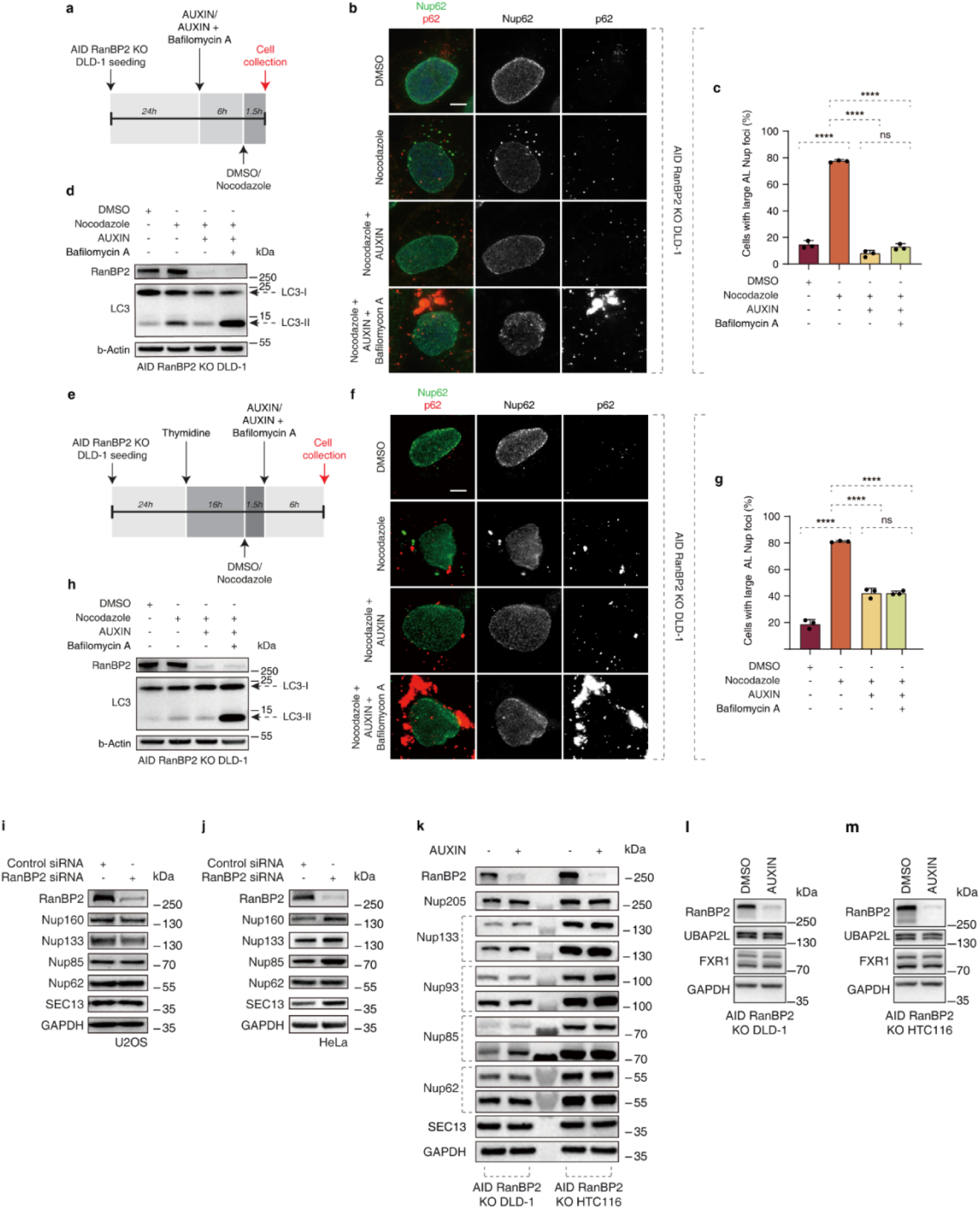
**The role of RanBP2 in the formation and maintenance of large AL is independent of autophagy.** a-d, Scheme of the experimental setup (a). AID-RanBP2 KO DLD-1 cells were treated with nocodazole (10 μM), auxin and bafilomycin at indicated timepoints. Cells were co-labelled with anti-Nup62 (green) and anti-p62 (red) antibodies. Representative images of cells are shown in (b). The number of cells with large AL foci was quantified in (c). (d) Shows the Western blot analysis of RanBP2 and LC3 under different conditions shown in (d). At least 200 cells per condition were analyzed (one-Way ANOVA test, mean ± SD, ns: not significant, ****P < 0.0001; N = 3). Scale bars, 5 μm. e-h, Scheme of the experimental setup (e). AID-RanBP3 KO DLD-1 cells were synchronized and arrested in S phase by thymidine treatment (16h). Subsequently, the cells were treated with nocodazole (10 μM), auxin and bafilomycin at indicated timepoints. Cells were co-labelled with anti-Nup62 (green) and anti-p62 (red) antibodies. Representative images of cells are shown in (f). The number of cells with large AL foci were quantified in (g). The Western blot analysis of RanBP2 and LC3 under different conditions shown in (h). At least 200 cells per condition were analyzed (one-Way ANOVA test, mean ± SD, ns: not significant, ****P < 0.0001; N = 3). Scale bars, 5 μm. i-k, Protein levels of representative nucleoporins were analyzed by Western blot in U2OS, HeLa and AID RanBP2 KO DLD-1 or HTC116 cells treated with RanBP2 siRNA or Auxin to induce depletion of RanBP2. l, m, Protein levels of FXR1 and UBAP2L were analyzed by Western blot in AID RanBP2 KO DLD-1 or HTC116 cells treated with DMSO or auxin to induce depletion of RanBP2 (4h) and synchronized in early interphase.

**Extended Data Fig. 11.**
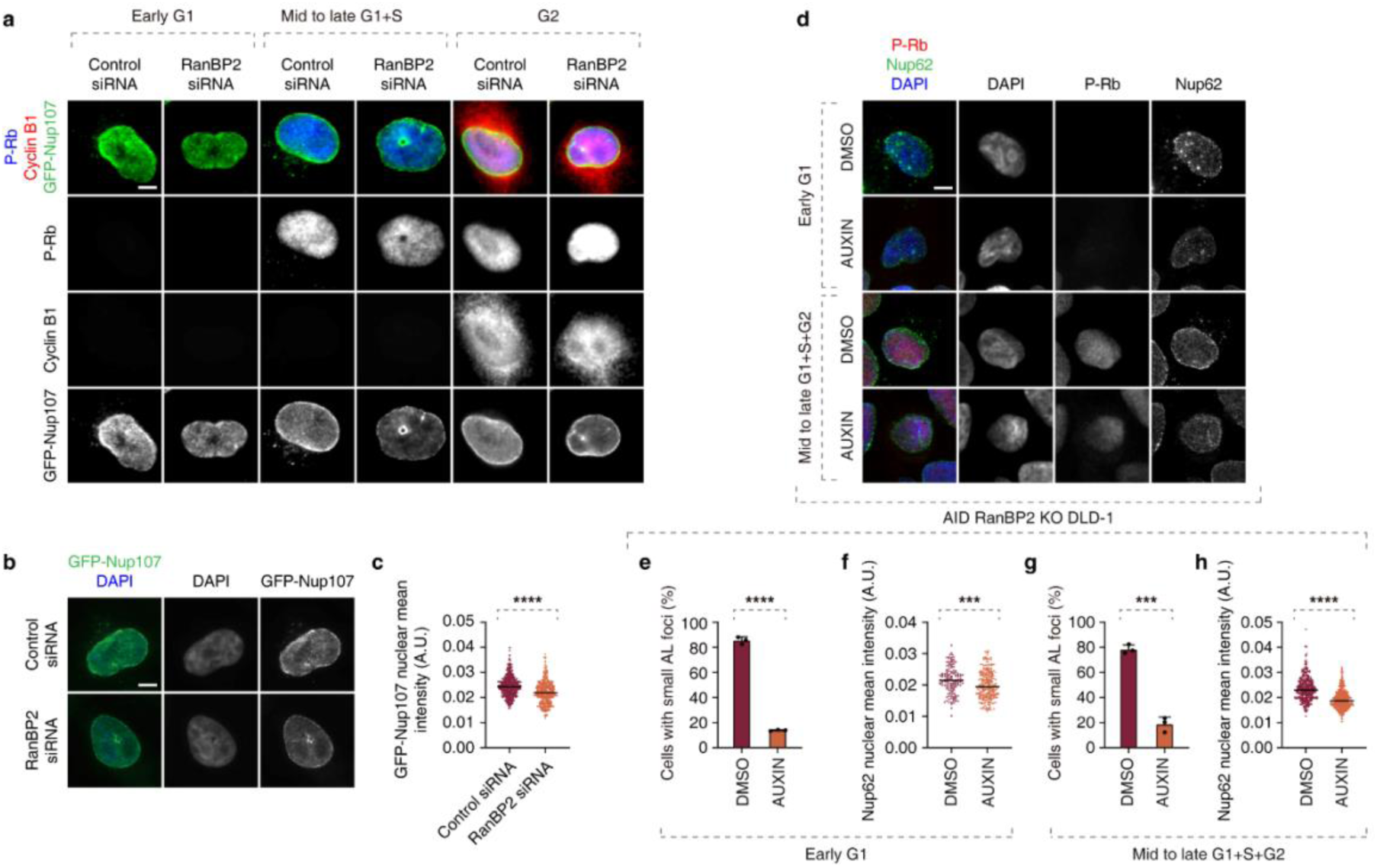
**RanBP2 is required for biogenesis of small AL and localization of Nups to the NE in non-stressed interphasic cells.** a, Representative images of asynchronously proliferating 2xZFN-mEGFP-Nup107 HeLa cells treated with the indicated siRNAs (48h) and co-labelled with anti-p-Rb (blue) and anti-cyclin B1 (red) antibodies. The percentage of cells with small AL foci in p-Rb and cyclin B1 negative cells (early G1), p-Rb positive and cyclin B1 negative cells (mid to late G1+S), and p-Rb and cyclin B1 positive cells (G2) was quantified in Fig, 4m. Scale bars, 5 μm. b, c, Representative images of 2xZFN-mEGFP-Nup107 HeLa cells treated with RanBP2 siRNA (48h) (b). Cells were co-labelled with DAPI (blue). The intensity of GFP-Nup107 at the NE was quantified in (c), and at least 400 cell per condition were analyzed (unpaired two-tailed t test, mean ± SD, ****P < 0.0001; N = 3). Scale bars, 5 μm. d-h, Representative immunofluorescence images depicting the localization of Nup62 in AID RanBP2 KO DLD-1 cells in different cell cycle stages. p-Rb was used to distinguish cells in early G1 (p-Rb–negative cells) and mid-late G1, S, and G2 (p-Rb–positive cells) stages. Nuclei were stained with DAPI (d). The percentage of cells with small AL foci in early G1 (e), and mid-late G1, S, G2 (g) was quantified. At least 300 cells per condition were analyzed (mean ± SD, ***P < 0.001; ****P < 0.0001; unpaired two-tailed t test, n = 3 independent experiments). The NE intensity of Nup62 in early G1 cells (f), and mid-late G1, S, G2 cells (H) was quantified. At least 150 cells per condition were analyzed (mean ± SD, ***P < 0.001; ****P < 0.0001; unpaired two-tailed t test, n = 3 independent experiments). Scale bars, 5 μm.

**Extended Data Fig. 12.**
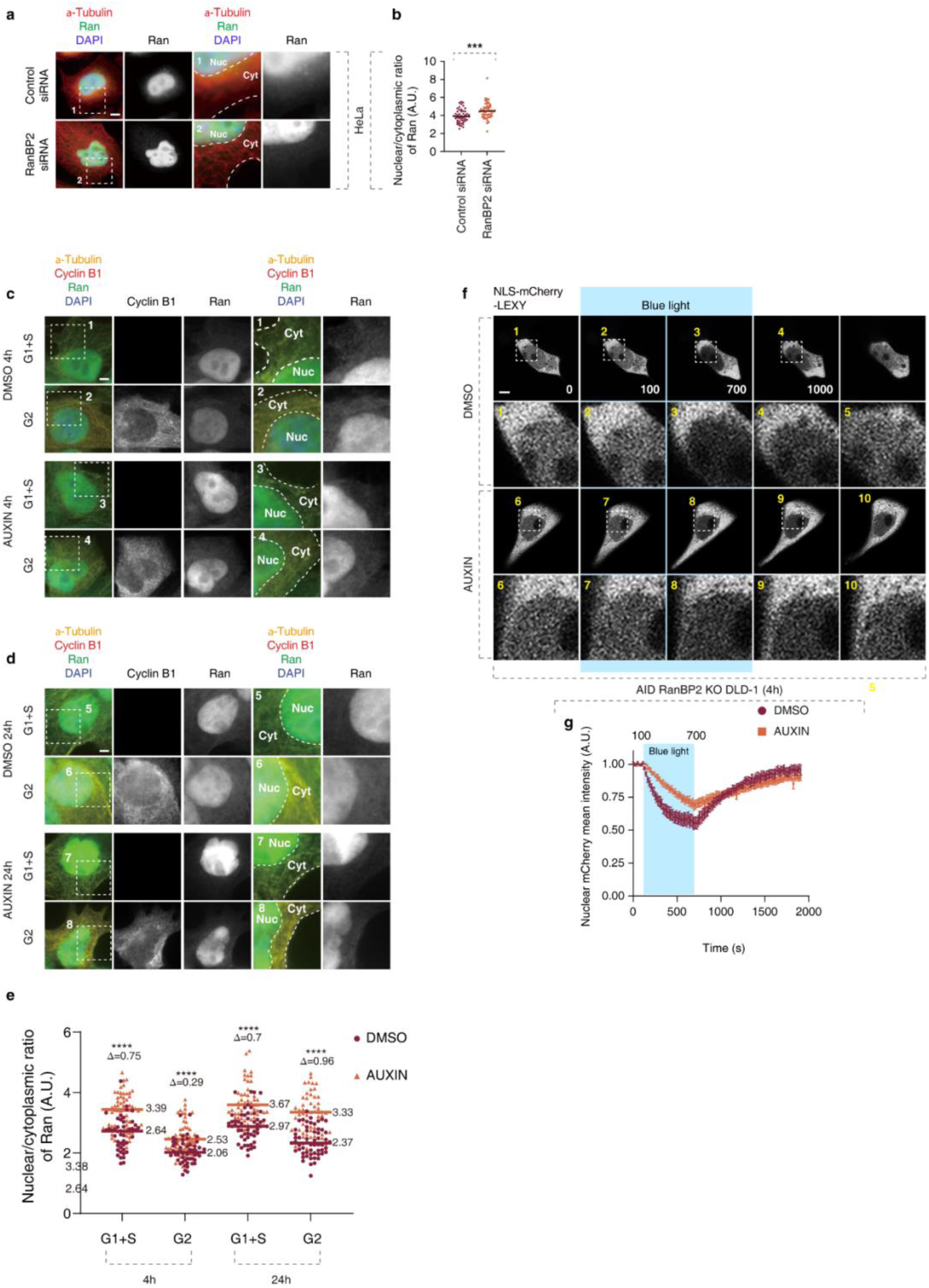
**RanBP2 supports nucleocytoplasmic transport.** a, b, Representative immunofluorescence images depicting the nuclear (Nuc) and cytoplasmic (Cyt) localization of Ran in asynchronously proliferating WT and RanBP2 KD (RanBP2 siRNA 48h) HeLa cells. Cells were co-labelled with anti-Ran (green), anti-alpha-Tubulin (red) antibodies and DAPI (a). The magnified framed regions are shown in the corresponding numbered panels. The N/C ratio of Ran was quantified (b). At least 100 cells per condition were analyzed (mean ± SD, ***P < 0.001; unpaired two-tailed t test, n = 3 independent experiments). Scale bar, 5 μm. c-e, Representative immunofluorescence images depicting the nuclear (Nuc) and cytoplasmic (Cyt) localization of Ran in asynchronously proliferating AID RanBP2 KO DLD-1 cells treated with DMSO or auxin (4h or 24h) to induce depletion of RanPB2. Cells were co-labelled with anti-Ran (green), anti-alpha-Tubulin-647 (yellow), anti-Cyclin B1 antibodies and DAPI (c and e). The magnified framed regions are shown in the corresponding numbered panels. The nuclear/cytoplasmic (N/C) ratio of Ran was quantified in (e). At least 100 cells per condition were analyzed (mean ± SD, ****P < 0.0001; unpaired two-tailed t test, n = 3 independent experiments). Scale bar, 5 μm. f, g, Representative immunofluorescence images depicting nuclear fluorescence NLS-mCherry-LEXY shuttling reporter expressed in AID RanBP2 KO DLD-1 cells treated with DMSO or auxin (4h) to induce depletion of RanPB2 (f). Cells were incubated in the dark for 100 s before blue light irradiation for 600 s was applied, which was followed by a 1200 s dark-recovery phase. Time is indicated in seconds. Relative nuclear fluorescence was quantified in (g). At least 10 cells per condition were analyzed (mean ± SD, n = 3 independent experiments).

**Extended Data Fig. 13.**
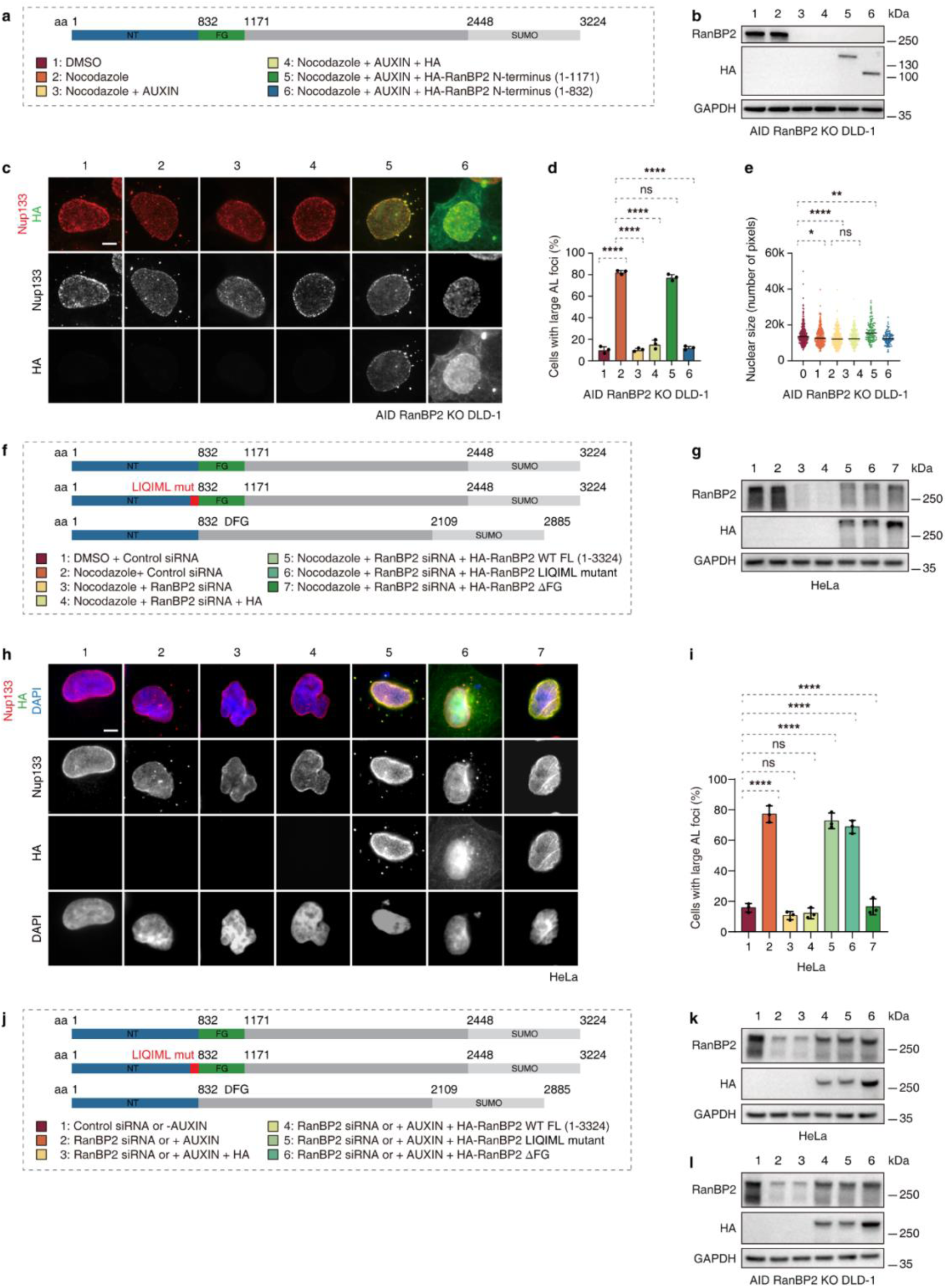
**RanBP2 is specifically required for the formation of large AL through its N-terminal FG repeat region.** a-e Rescue experiments using different RanBP2 protein fragments (a). (b) Shows the Western blot analysis of RanBP2 fragments under different conditions indicated. (c) shows representative images of AID-RanBP2 KO DLD-1 cells treated with the DMSO or auxin, transfected with different RanBP2 fragments and treated with nocodazole (10 μM) or DMSO (90 min). Cells were co-labelled with anti-Nup133 (red) and anti-HA (green) antibodies. The number of cells with large AL foci was quantified in (d), the nuclear size was quantified in (e), and at least 300 cells per condition were analyzed (one-Way ANOVA test, mean ± SD, ns: not significant, *P < 0.05; **P < 0.01; ****P < 0.0001; N = 3). Scale bar, 5 μm. f-I, Rescue experiments using different RanBP2 protein version (f). (g) Shows the Western blot analysis of RanBP2 version under different conditions indicated (numbers in the schematic diagram). (h) shows representative images of HeLa cells treated with the control or RanBP2 siRNAs, transfected with different RanBP2 version and treated with nocodazole (10 μM) or DMSO (90 min). Cells were co-labelled with anti-Nup133 (red) and anti-HA (green) antibodies. The number of cells with large AL foci was quantified in (i), and at least 300 cells per condition were analyzed (one-Way ANOVA test, mean ± SD, ns: not significant, *P < 0.05; **P < 0.01; ****P < 0.0001; N = 3). Scale bar, 5 μm. j-l, Rescue experiments using different RanBP2 protein versions (j). (k and l) The Western blot analysis of RanBP2 fragments under different conditions indicated (numbers in the schematic diagram). Representative images and analysis are shown in Fig 5J and 5K.

**Extended Data Fig. 14.**
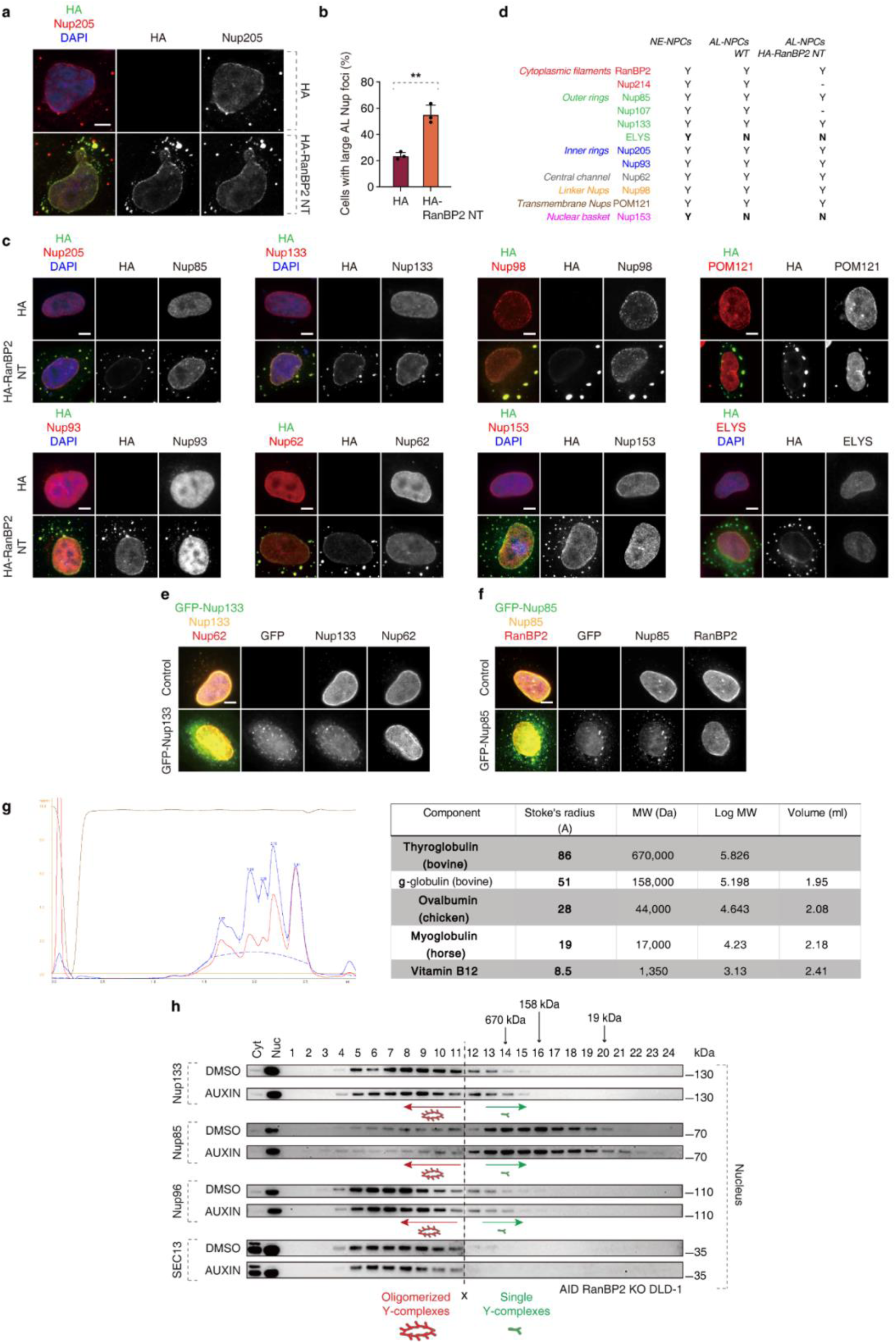
**N-terminal fragment of RanBP2 promotes AL formation.** a, b, Representative images of HeLa cells transfected with HA or HA-RanBP2 N-terminal (NT) fragment. Cells were co-labelled with DAPI (blue), anti-Nup205 (red) and anti-HA (green) antibodies (a). The number of cells with large AL foci was quantified in (b), and at least 300 cells per condition were analyzed (unpaired two-tailed t test, mean ± SD, **P < 0.01; N = 3). Scale bar, 5 μm. c, Representative images of HeLa cells transfected with HA or HA-RanBP2 N-terminal (NT) fragment and labelled for Nups from different NPC subcomplexes as indicated. Cells were co-labelled with DAPI (blue), anti-Nups (red) and anti-HA (green) antibodies. Scale bars, 5 μm. d, Summary of nucleoporin composition in NE-NPCs, ALC-NPCs and AL-NPCs-induced by RanBP2-NT expression. Different colors correspond to specific NPC subcomplexes. “Y” indicates presence and “N” indicates absence of specific Nups and “-” indicates no available data. e, Representative images of HeLa cells transfected with GFP-Nup133. Cells were co-labelled with Nup133 (yellow), and anti-Nup62 (red) antibodies. Scale bars, 5 μm. f, Representative images of HeLa cells transfected with GFP-Nup85. Cells were co-labelled with Nup85 (yellow), and anti-RanBP2 (red) antibodies. Scale bars, 5 μm. Note that expression of RanBP2-NT but not Nup133 or Nup85 induce large AL foci where multiple Nups can co-localize. g, Calibration of the Superose 6 column was carried out with thyroglobulin (bovine), g-globulin (bovine), ovalbumin (chicken), myoglobulin (horse) and vitamin B12. The calibration profile and the molecular weights of the protein markers are indicated h, Auxin-inducible degron (AID) RanBP2 KO DLD-1 cells were treated with DMSO or Auxin for 4h to deplete RanBP2 and nuclear fractions were prepared, separated on a Superose 6 gel filtration column and analyzed by Western blotting (at least 3 independent experiments). Fraction 1 corresponds to the void and the elution of thyroglobulin (670 kDa), g-globulin (158 kDa) and myoglobulin markers 19 kDa are indicated. Green arrows point to fractions preferentially containing single Y-complexes, and red arrows indicate fraction with higher molecular weight, oligomerized Y-complexes. Note that depletion of RanBP2 did not affect the abundance of oligomerized Y-complexes in the nuclear fractions.

**Extended Data Fig. 15.**
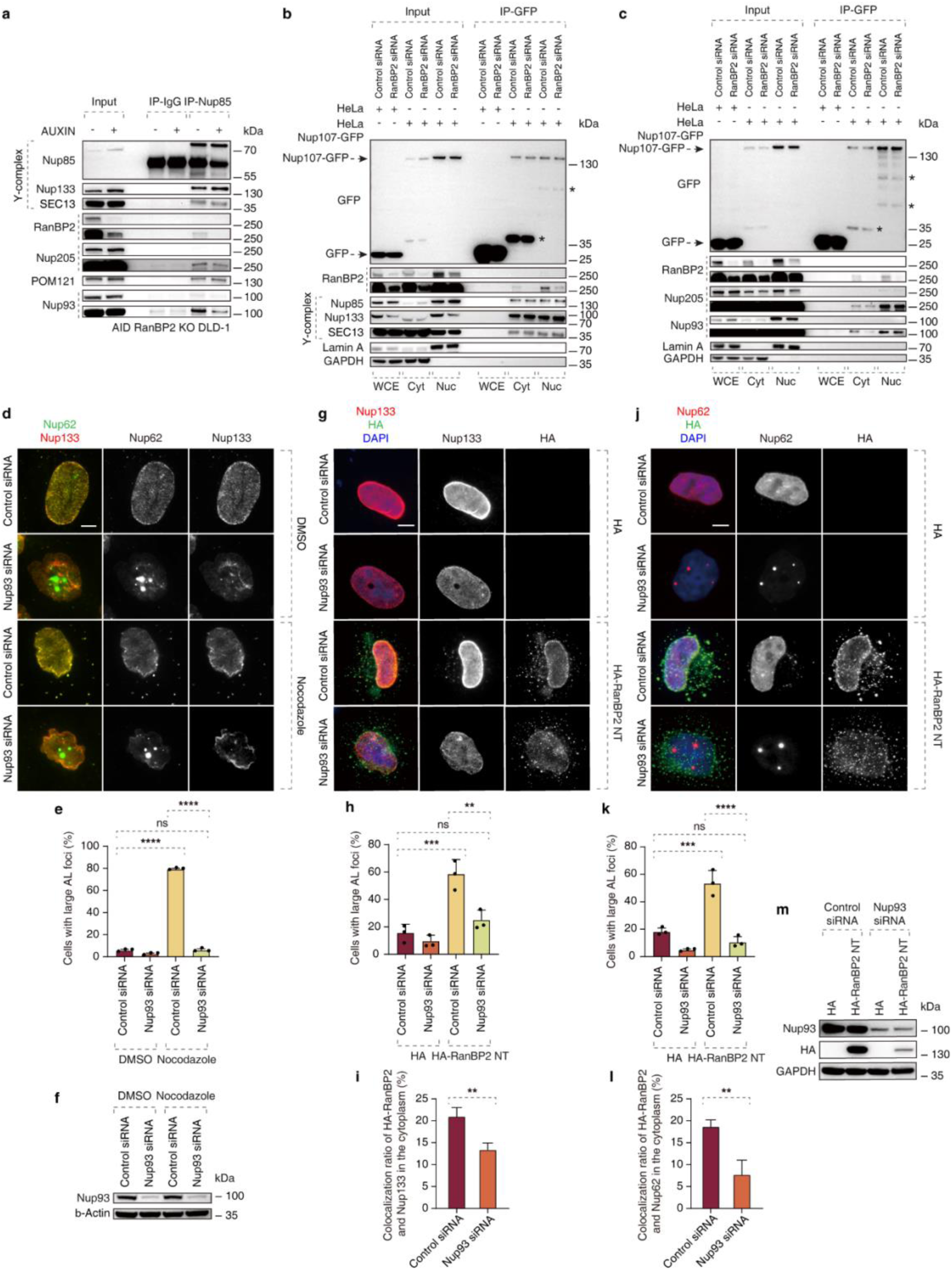
**RanBP2 drives the assembly of cytosolic AL-NPCs by promoting the interaction between Nup93 and the Y complex.** a, AID-RanBP2 KO DLD-1 cells were treated with auxin to induce depletion of RanBP2 and lysates were prepared and immunoprecipitated using Nup85 antibody or IgG and analyzed by Western blot (at least 3 independent experiments). b, c, Cytoplasmic or nuclear extracts of Hela cells expressing GFP or 2xZFN-mEGFP-Nup107 were immunoprecipitated using GFP-Trap A beads (GFP-IP) and analyzed by Western blot (at least 3 independent experiments). d-f, Representative images of HeLa cells treated with the indicated siRNAs (48h) and subsequently with nocodazole (10 μM) or DMSO (90 min). Cells were co-labelled with anti-Nup62 (green) and anti-Nup133 (red) (d) antibodies. The number of cells with large AL foci was quantified in (e) and at least 1000 cells per condition were analyzed (one-Way ANOVA test, mean ± SD, ns: not significant, ****P < 0.0001; N = 3). (f) Shows the Western blot analysis of Nup93 under indicated conditions. Scale bars, 5 μm. g-m, Representative images of HeLa cells treated with the indicated siRNAs and transfected with HA or HA-RanBP2-NT fragment. Cells were co-labelled with anti-Nup133 (red), anti-HA (green) antibodies and DAPI (blue) (g), or co-labelled with anti-Nup62 (red), anti-HA (green) antibodies and DAPI (blue) (j). Non-AL Nup62 foci, which does not contain Nup133, is produced in the cell nucleus(g, j). The number of cells with large AL foci was quantified in (h, k) and the co-localization ratio of HA-RanBP2-NT and Nup62 was quantified in (i, l), and at least 300 cells per condition were analyzed (unpaired two-tailed t test, or one-Way ANOVA test, mean ± SD, **P < 0.01; ***P < 0.001; N = 3). Scale bars, 5 μm. (m) Shows the Western blot analysis of Nup93 and HA-RanBP2-NT under indicated conditions.

**Extended Data Fig. 16.**
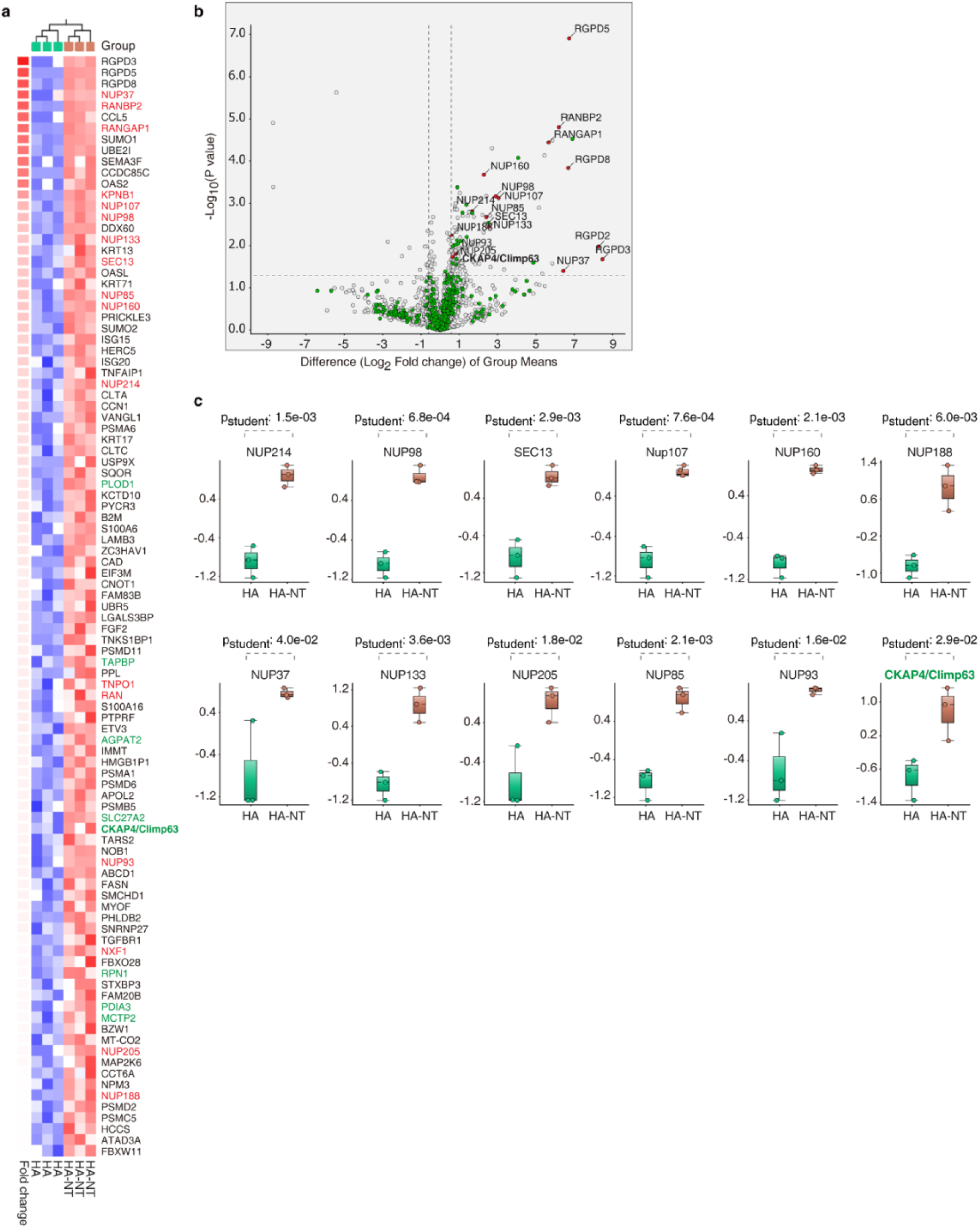
**RanBP2-dependent AL interactome.** a-c, Liquid chromatography-mass spectrometry (LC-MS) was used to identify proteins interacting with HA-tagged NT-RanBP2 and the HA-tag as a negative control. A heatmap displays the proteins that showed a change greater than 1.5-fold with a p-value of less than 0.05 between the two groups (a). Volcano plot for the comparison between the two groups (b). The cutoff values fold change >1.5 and FDR < 0.05 were utilized to identify differentially interacting protein. Red represents a group of NPC-related proteins and green depicted ER-related factors. (c) shows the box plots of selected Nups and Climp63 (CKAP4).

**Extended Data Fig. 17.**
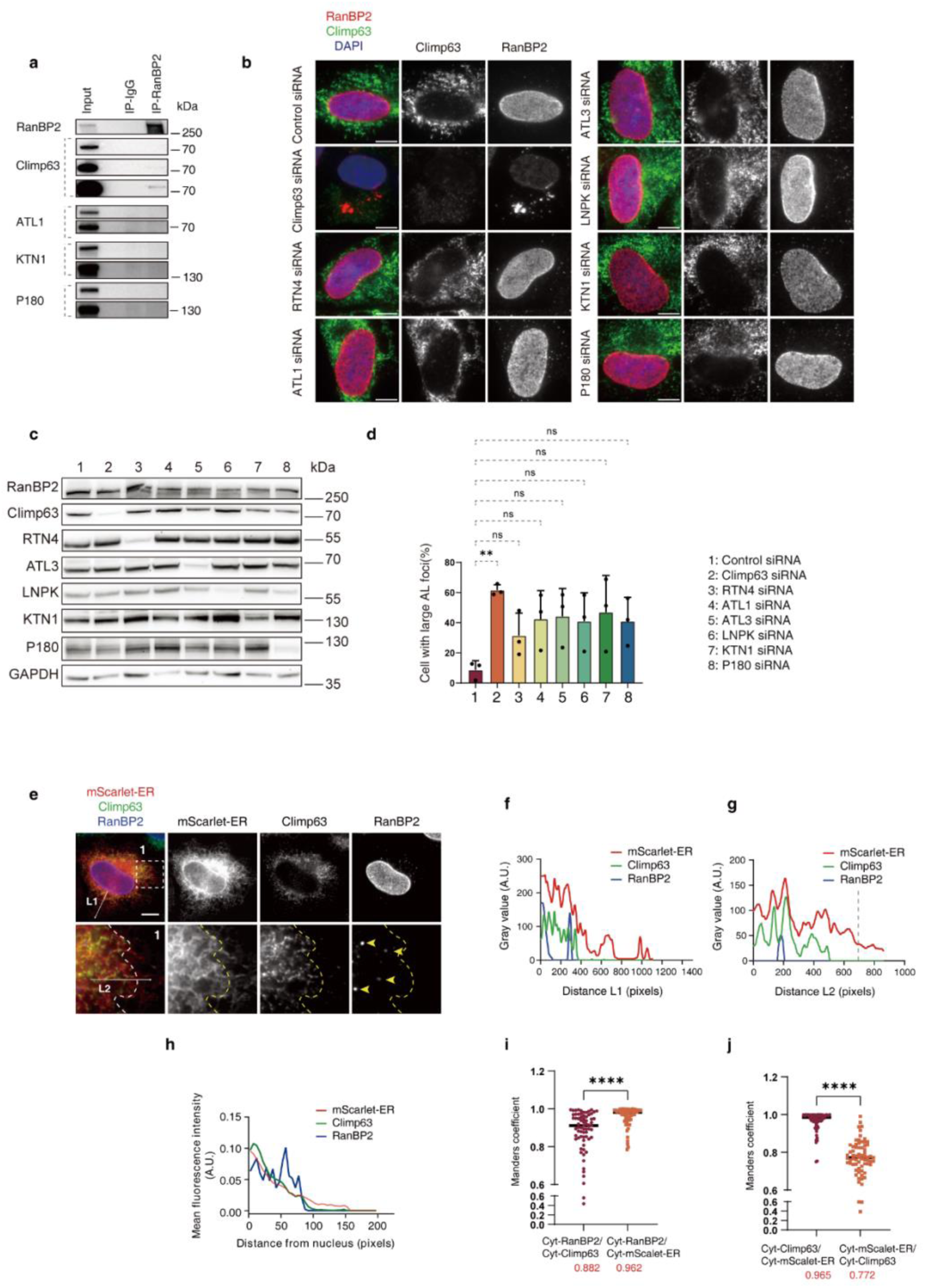
**Climp63 facilitates the localization of AL-NPCs to ER sheets.** a, Lysates of HeLa cells were immunoprecipitated using RanBP2 antibody or IgG and analyzed by Western blot (at least 3 independent experiments). b-d, Representative images of MRC5 cells treated with the indicated siRNAs. Cells were co-labelled with anti-RanBP2 (red), anti-Climp63 (green) antibodies and DAPI (blue) (b). Protein levels of representative nucleoporins were analyzed by Western blot (c). The number of cells with large AL foci was quantified in (d) (one-Way ANOVA test, mean ± SD, ns: not significant; **P < 0.01; N = 3). Scale bars, 5 μm. e-j, Representative images of HeLa cells expressing ER marker plasmid mScarlet-ER. Cells were co-labelled with anti-Climp63 (green) and anti-RanBP2 (blue) antibodies (e). The magnified framed regions are shown in the corresponding numbered panels. The dotted line represents the boundary between ER sheets labeled by Climp63 and more distal ER tubules. Intensity profiles along arrows L1 and L2 showing signals from all three fluorescent channels are shown (f and g). (h) shown the Intensity profiles per distance from the nucleus of three fluorescent channels. The co-localization of Climp63, mScarlet-ER, and RanBP2 located in the cytoplasm was measured by in (i, j) (mean ± SD, ****P < 0.0001, unpaired two-tailed t test; 150 cells were counted)

## Notes

### Competing Interest Statement

The authors have declared no competing interest.

### Summary of Updates

This version of the manuscript has been revised to include updated experimental evidence supporting the role of AL as an additional assembly route for the NPC.

## References

1. Kessel, R. G. Annulate lamellae: a last frontier in cellular organelles. Int Rev Cytol 133, 43–120 (1992).

2. Kessel, R. G. The annulate lamellae--from obscurity to spotlight. Electron Microsc Rev 2, 257–348 (1989).

3. Rawe, V. Y., Olmedo, S. B., Nodar, F. N., Ponzio, R. & Sutovsky, P. Abnormal assembly of annulate lamellae and nuclear pore complexes coincides with fertilization arrest at the pronuclear stage of human zygotic development. Hum Reprod 18, 576–582 (2003).

4. Zybina, E. V. & Zybina, T. G. Modifications of nuclear envelope during differentiation and depolyploidization of rat trophoblast cells. Micron 39, 593–606 (2008).

5. Caruso, R. A. et al. Modifications of nuclear envelope in tumour cells of human gastric carcinomas: an ultrastructural study. Anticancer Res 30, 699–702 (2010).

6. Gil-Perotín, S. et al. Adult neural stem cells from the subventricular zone: a review of the neurosphere assay. Anat Rec (Hoboken) 296, 1435–1452 (2013).

7. Hampoelz, B. et al. Pre-assembled Nuclear Pores Insert into the Nuclear Envelope during Early Development. Cell 166, 664–678 (2016).

8. Underwood, J. M., Becker, K. A., Stein, G. S. & Nickerson, J. A. The Ultrastructural Signature of Human Embryonic Stem Cells. J Cell Biochem 118, 764–774 (2017).

9. Grandi, G., Astolfi, G., Chicca, M. & Pezzi, M. Ultrastructural investigations on spermatogenesis and spermatozoan morphology in the endangered Adriatic sturgeon, Acipenser naccarii (Chondrostei, Acipenseriformes). J Morphol 279, 1376–1396 (2018).

10. Hampoelz, B. et al. Nuclear Pores Assemble from Nucleoporin Condensates During Oogenesis. Cell 179, 671–686.e17 (2019).

11. Dymek, A. M., Pecio, A. & Piprek, R. P. Diversity of Balbiani body formation in internally and externally fertilizing representatives of Osteoglossiformes (Teleostei: Osteoglossomorpha). J Morphol 282, 1313–1329 (2021).

12. McCulloch, D. Fibrous structures in the ground cytoplasm of the Arbacia egg. J. Exp. Zool. 119, 47–63 (1952).

13. Swift, H. The fine structure of annulate lamellae. J Biophys Biochem Cytol 2, 415–418 (1956).

14. Lin, J. & Sumara, I. Cytoplasmic nucleoporin assemblage: the cellular artwork in physiology and disease. Nucleus 15, 2387534 (2024).

15. Ren, H. et al. Postmitotic annulate lamellae assembly contributes to nuclear envelope reconstitution in daughter cells. J Biol Chem 294, 10383–10391 (2019).

16. Raghunayakula, S., Subramonian, D., Dasso, M., Kumar, R. & Zhang, X.-D. Molecular Characterization and Functional Analysis of Annulate Lamellae Pore Complexes in Nuclear Transport in Mammalian Cells. PLoS One 10, e0144508 (2015).

17. Patterson, J. R., Wood, M. P. & Schisa, J. A. Assembly of RNP granules in stressed and aging oocytes requires nucleoporins and is coordinated with nuclear membrane blebbing. Dev Biol 353, 173–185 (2011).

18. Sahoo, M. R. et al. Nup358 binds to AGO proteins through its SUMO-interacting motifs and promotes the association of target mRNA with miRISC. EMBO Rep 18, 241–263 (2017).

19. Rawe, V. Y., Olmedo, S. B., Nodar, F. N., Ponzio, R. & Sutovsky, P. Annulate lamellae assembly in non-inseminated and failed fertilized human oocytes. Fertility and Sterility 78, S163 (2002).

20. Boulware, M. J. & Marchant, J. S. Nuclear pore disassembly from endoplasmic reticulum membranes promotes Ca2+ signalling competency. J Physiol 586, 2873–2888 (2008).

21. Boulware, M. J. & Marchant, J. S. IP3 receptor activity is differentially regulated in endoplasmic reticulum subdomains during oocyte maturation. Curr Biol 15, 765–770 (2005).

22. Jühlen, R. & Fahrenkrog, B. From the sideline: Tissue-specific nucleoporin function in health and disease, an update. FEBS Letters 597, 2750–2768 (2023).

23. Coyne, A. N. & Rothstein, J. D. Nuclear pore complexes - a doorway to neural injury in neurodegeneration. Nat Rev Neurol 18, 348–362 (2022).

24. Kuiper, E. F. E. et al. The chaperone DNAJB6 surveils FG-nucleoporins and is required for interphase nuclear pore complex biogenesis. Nat Cell Biol 24, 1584–1594 (2022).

25. Agote-Aran, A. et al. Spatial control of nucleoporin condensation by fragile X-related proteins. EMBO J 39, e104467 (2020).

26. Weberruss, M. & Antonin, W. Perforating the nuclear boundary - how nuclear pore complexes assemble. J Cell Sci 129, 4439–4447 (2016).

27. Agote-Arán, A., Lin, J. & Sumara, I. Fragile X-Related Protein 1 Regulates Nucleoporin Localization in a Cell Cycle-Dependent Manner. Front Cell Dev Biol 9, 755847 (2021).

28. Liao, Y. et al. UBAP2L ensures homeostasis of nuclear pore complexes at the intact nuclear envelope. Journal of Cell Biology 223, e202310006 (2024).

29. Andronov, L., Vonesch, J.-L. & Klaholz, B. P. Practical Aspects of Super-Resolution Imaging and Segmentation of Macromolecular Complexes by dSTORM. Methods Mol Biol 2247, 271–286 (2021).

30. Lelek, M. et al. Single-molecule localization microscopy. Nat Rev Methods Primers 1, 39 (2021).

31. Andronov, L., Genthial, R., Hentsch, D. & Klaholz, B. P. splitSMLM, a spectral demixing method for high-precision multi-color localization microscopy applied to nuclear pore complexes. Commun Biol 5, 1100 (2022).

32. Davis, L. I. & Blobel, G. Identification and characterization of a nuclear pore complex protein. Cell 45, 699–709 (1986).

33. Kessel, R. G. The structure and function of annulate lamellae: porous cytoplasmic and intranuclear membranes. Int Rev Cytol 82, 181–303 (1983).

34. Sachweh, J. et al. The small GTPase Ran defines Nuclear Pore Complex Asymmetry. Preprint at 10.1101/2024.10.10.617378 (2024).

35. Rasala, B. A., Ramos, C., Harel, A. & Forbes, D. J. Capture of AT-rich chromatin by ELYS recruits POM121 and NDC1 to initiate nuclear pore assembly. Mol Biol Cell 19, 3982–3996 (2008).

36. Walther, T. C. et al. RanGTP mediates nuclear pore complex assembly. Nature 424, 689–694 (2003).

37. Maul, G. G. et al. Time sequence of nuclear pore formation in phytohemagglutinin-stimulated lymphocytes and in HeLa cells during the cell cycle. J Cell Biol 55, 433–447 (1972).

38. McKinney, S. A., Murphy, C. S., Hazelwood, K. L., Davidson, M. W. & Looger, L. L. A bright and photostable photoconvertible fluorescent protein. Nat Methods 6, 131–133 (2009).

39. De Magistris, P. & Antonin, W. The Dynamic Nature of the Nuclear Envelope. Current Biology 28, R487–R497 (2018).

40. Chen, S., Novick, P. & Ferro-Novick, S. ER structure and function. Current Opinion in Cell Biology 25, 428–433 (2013).

41. Deolal, P., Scholz, J., Ren, K., Bragulat-Teixidor, H. & Otsuka, S. Sculpting nuclear envelope identity from the endoplasmic reticulum during the cell cycle. Nucleus 15, 2299632 (2024).

42. Aksenova, V., Arnaoutov, A. & Dasso, M. Analysis of Nucleoporin Function Using Inducible Degron Techniques. Methods Mol Biol 2502, 129–150 (2022).

43. Tomioka, Y. et al. TORC1 inactivation stimulates autophagy of nucleoporin and nuclear pore complexes. J Cell Biol 219, e201910063 (2020).

44. Lee, C.-W. et al. Selective autophagy degrades nuclear pore complexes. Nat Cell Biol 22, 159–166 (2020).

45. Bley, C. J. et al. Architecture of the cytoplasmic face of the nuclear pore. Science 376, eabm9129 (2022).

46. Zhang, K. et al. The C9orf72 repeat expansion disrupts nucleocytoplasmic transport. Nature 525, 56–61 (2015).

47. Niopek, D., Wehler, P., Roensch, J., Eils, R. & Di Ventura, B. Optogenetic control of nuclear protein export. Nat Commun 7, 10624 (2016).

48. Von Appen, A. et al. In situ structural analysis of the human nuclear pore complex. Nature 526, 140–143 (2015).

49. Onischenko, E. et al. Natively Unfolded FG Repeats Stabilize the Structure of the Nuclear Pore Complex. Cell 171, 904–917.e19 (2017).

50. Vollmer, B. & Antonin, W. The diverse roles of the Nup93/Nic96 complex proteins – structural scaffolds of the nuclear pore complex with additional cellular functions. Biological Chemistry 395, 515–528 (2014).

51. Schroeder, L. K. et al. Dynamic nanoscale morphology of the ER surveyed by STED microscopy. Journal of Cell Biology 218, 83–96 (2019).

52. Shibata, Y. et al. Mechanisms Determining the Morphology of the Peripheral ER. Cell 143, 774–788 (2010).

53. Klopfenstein, D. R. Ch., Kappeler, F. & Hauri, H.-P. A novel direct interaction of endoplasmic reticulum with microtubules. The EMBO Journal 17, 6168–6177 (1998).

54. Otsuka, S. & Ellenberg, J. Mechanisms of nuclear pore complex assembly - two different ways of building one molecular machine. FEBS Lett 592, 475–488 (2018).

55. Penzo, A. & Palancade, B. Puzzling out nuclear pore complex assembly. FEBS Letters 597, 2705–2727 (2023).

56. Dultz, E., Wojtynek, M., Medalia, O. & Onischenko, E. The Nuclear Pore Complex: Birth, Life, and Death of a Cellular Behemoth. Cells 11, 1456 (2022).

57. Otsuka, S. et al. A quantitative map of nuclear pore assembly reveals two distinct mechanisms. Nature 613, 575–581 (2023).

58. Otsuka, S. et al. Postmitotic nuclear pore assembly proceeds by radial dilation of small membrane openings. Nat Struct Mol Biol 25, 21–28 (2018).

59. Vitale, J., Khan, A., Neuner, A. & Schiebel, E. A perinuclear α-helix with amphipathic features in Brl1 promotes NPC assembly. Mol Biol Cell 33, ar35 (2022).

60. Kralt, A. et al. An amphipathic helix in Brl1 is required for nuclear pore complex biogenesis in S. cerevisiae. Elife 11, e78385 (2022).

61. Mondal, S., Neuner, A., Khan, A., Vitale, J. & Schiebel, E. Multifunctional Roles of Brr6 and Brl1 in Nuclear Envelope Fusion During Nuclear Pore Complex Biogenesis. Preprint at 10.1101/2025.07.22.665954 (2025).

62. Rampello, A. J. et al. Torsin ATPase deficiency leads to defects in nuclear pore biogenesis and sequestration of MLF2. J Cell Biol 219, e201910185 (2020).

63. Kim, S. et al. TorsinA is essential for neuronal nuclear pore complex localization and maturation. Nat Cell Biol 26, 1482–1495 (2024).

64. Joseph, J. & Dasso, M. The nucleoporin Nup358 associates with and regulates interphase microtubules. FEBS Lett 582, 190–196 (2008).

65. Cui, H. et al. Adapter Proteins for Opposing Motors Interact Simultaneously with Nuclear Pore Protein Nup358. Biochemistry 58, 5085–5097 (2019).

66. Ciccarelli, F. D. et al. Complex genomic rearrangements lead to novel primate gene function. Genome Res 15, 343–351 (2005).

67. Hetzer, M. W. The nuclear envelope. Cold Spring Harb Perspect Biol 2, a000539 (2010).

68. Talamas, J. A. & Hetzer, M. W. POM121 and Sun1 play a role in early steps of interphase NPC assembly. Journal of Cell Biology 194, 27–37 (2011).

69. Eymieux, S., Blanchard, E., Uzbekov, R., Hourioux, C. & Roingeard, P. Annulate lamellae and intracellular pathogens. Cellular Microbiology 23, (2021).

70. Lin, Y.-C. et al. Interactions between ALS-linked FUS and nucleoporins are associated with defects in the nucleocytoplasmic transport pathway. Nat Neurosci 24, 1077–1088 (2021).

71. Gleixner, A. M. et al. NUP62 localizes to ALS/FTLD pathological assemblies and contributes to TDP-43 insolubility. Nat Commun 13, 3380 (2022).

72. Suhr, S. T. et al. Identities of sequestered proteins in aggregates from cells with induced polyglutamine expression. J Cell Biol 153, 283–294 (2001).

73. Grima, J. C. et al. Mutant Huntingtin Disrupts the Nuclear Pore Complex. Neuron 94, 93–107.e6 (2017).

74. Pappas, S. S., Liang, C.-C., Kim, S., Rivera, C. O. & Dauer, W. T. TorsinA dysfunction causes persistent neuronal nuclear pore defects. Hum Mol Genet 27, 407–420 (2018).

75. Zouiouich, M. et al. MOSPD2 is an endoplasmic reticulum-lipid droplet tether functioning in LD homeostasis. J Cell Biol 221, e202110044 (2022).

76. Ran, F. A. et al. Genome engineering using the CRISPR-Cas9 system. Nat Protoc 8, 2281–2308 (2013).

77. Chou, Y.-Y. et al. Inherited nuclear pore substructures template post-mitotic pore assembly. Dev Cell 56, 1786–1803.e9 (2021).

78. Mossink, B. et al. Human neuronal networks on micro-electrode arrays are a highly robust tool to study disease-specific genotype-phenotype correlations in vitro. Stem Cell Reports 16, 2182–2196 (2021).

79. Andronov, L., Genthial, R., Hentsch, D. & Klaholz, B. P. splitSMLM, a spectral demixing method for high-precision multi-color localization microscopy applied to nuclear pore complexes. Commun Biol 5, 1100 (2022).

80. Andronov, L., Lutz, Y., Vonesch, J.-L. & Klaholz, B. P. SharpViSu: integrated analysis and segmentation of super-resolution microscopy data. Bioinformatics 32, 2239–2241 (2016).

81. Andronov, L. et al. 3DClusterViSu: 3D clustering analysis of super-resolution microscopy data by 3D Voronoi tessellations. Bioinformatics 34, 3004–3012 (2018).

82. Mastronarde, D. N. Automated electron microscope tomography using robust prediction of specimen movements. J Struct Biol 152, 36–51 (2005).

83. Kremer, J. R., Mastronarde, D. N. & McIntosh, J. R. Computer visualization of three-dimensional image data using IMOD. J Struct Biol 116, 71–76 (1996).

84. Paul-Gilloteaux, P. et al. eC-CLEM: flexible multidimensional registration software for correlative microscopies. Nat Methods 14, 102–103 (2017).

85. de Chaumont, F. et al. Icy: an open bioimage informatics platform for extended reproducible research. Nat Methods 9, 690–696 (2012).

86. van der Walt, S. et al. scikit-image: image processing in Python. PeerJ 2, e453 (2014).

87. Boni, A. et al. Live imaging and modeling of inner nuclear membrane targeting reveals its molecular requirements in mammalian cells. J Cell Biol 209, 705–720 (2015).

88. Miura, K. Measurements of Intensity Dynamics at the Periphery of the Nucleus. in Bioimage Data Analysis Workflows (eds. Miura, K. & Sladoje, N.) 9–32 (Springer International Publishing, Cham, 2020). doi:10.1007/978-3-030-22386-1_2.

